# Differences in interactions between brain regions across personality types while learning in zebrafish (*Danio rerio*)

**DOI:** 10.64898/2026.09.09.750425

**Authors:** Jamie Corcoran, Ryan Y. Wong

**Affiliations:** Department of Psychology, University of Nebraska at Omaha, Omaha, NE; Department of Biology, University of Nebraska at Omaha, Omaha, NE

**Keywords:** **Keywords**: personality, learning, bold, shy, functional connectivity, neurotransmitter, dopamine, serotonin

## Abstract

Learning is an important process for survival. Many studies find that variation in learning ability is related to personality type. Bold individuals generally learn faster than shy, however, the underlying neural mechanisms are not well understood. Potential mechanisms for differences in learning speed are variation in neurotransmitter activity, specifically dopamine and serotonin, and functional connectivity across the brain. In this study we quantified the number of generally active, active dopaminergic, and active serotonergic neurons using immunohistochemistry across 12 brain regions at two timepoints in bold and shy zebrafish undergoing an associative learning task. There were no interactions between treatment and personality type in any of the cell count measures suggesting that individual region activity and neurotransmitter activity do not explain learning differences. However, analyses of general neural activity networks revealed that the connection from the posterior zone of the dorsal telencephalon (olfactory cortex homolog) to lateral zone of the dorsal telencephalon (hippocampus homolog) was positive in bold but negative in shy fish undergoing learning. Overall, the results suggest that differences in activity involving reward processing in the brain may explain differences in reward learning between personality types.

## Introduction

Learning is an important process that allows individuals to acquire new knowledge about their environment to survive. Individuals consistently differ in their ability to learn and these differences are related to other individual differences such as personality type [1–9]. One common animal personality trait is boldness and shyness [10]. Bold individuals investigate novel stimuli, explore more, and display less behavioral indications of stress while shy individuals tend to be neophobic, explore less and display more behavioral indications of stress [10,11]. The opposing behavioral responses between the personality types have been hypothesized to be linked to general cognitive biases. Specifically, studies show that bold personality types have faster changes in behavior (i.e. faster learning) compared to shy individuals in reward tasks [3–9]. Although studies find similar relationships between personality and learning, the underlying neural mechanisms are not well known.

One neural mechanism that is important for both reward learning and personality type is neurotransmitter activity. It has been hypothesized that changes in dopamine and serotonin activity can explain the link between learning and personality types as both are associated with learning and individual differences in behavior related to personality type [12–15]. Prior studies showed that individual differences in learning are associated with different tyrosine hydroxylase (rate limiting step of dopamine synthesis) levels where high dopamine activity in the nucleus accumbens is associated with enhanced learning in a positive reinforcement task [16–19]. Similarly, serotonin levels are associated with learning where with less serotonin activity enhances learning in positive reinforcement tasks [21]. Therefore, activity involving dopaminergic and serotonergic areas of the brain are key candidates for modulating differences in learning between personality types.

Another potential mechanism for differences in cognition between personality type is variation in functional network activity comprising of brain regions involved in learning. Cognition and personality can be predicted by differences in network activity measurements [23–25]. Changes in connectivity of regions in the frontal cortex and other regions described below are characteristic of individual differences in cognition and personality. Regions implicated in learning and personality networks in teleosts are the medial zone of dorsal telencephalon (Dm), lateral zone of dorsal telencephalon (Dl), posterior zone of the dorsal telencephalon (Dp), ventral hypothalamus (Hv) and habenula (Hb) [26–31]. How general activity or neurotransmitter activity in each of these regions or across a network comprising these regions relate to personality and learning is unclear.

The goal of this study was to identify neural mechanisms linked to differences in learning between personality types. We used zebrafish selectively bred to exhibit bold and shy personality types to assess how general network activity, serotonin and dopamine activity may explain faster learning in bold animals. For neurotransmitter activity, we hypothesized that there would be an increase in serotonin activity and decreased dopamine activity in shy fish compared to bold fish. For network connectivity, we hypothesized that there would be differences in functional connectivity especially in interactions with the Dm, Dl, Dp, Hb and Hv. These differences will provide insight into how these personality types differ in learning ability.

## Methods

### Animals

We used zebrafish from lines that were selectively bred in the lab to exhibit a shy (HSB) or bold (LSB) personality type [32]. The lines are maintained through artificial selection based on freezing behavior in response to exposure to a novel open field environment [32]. These lines consistently differ in behavioral stress responses across 5 other behavioral stress assays [32] and maintain rank order consistency within and between lines in the open field test across 5 weeks [33]. Subsequent generations of these lines similarly show consistent behavioral differences between the lines in response to a novelty stressor and other behavioral assays [31,33–37]. The shy personality type line shows a faster whole-body cortisol release rate in response to a novelty stressor compared to bold individuals, which is consistent with another key trait difference between personality types [34]. Additionally, the lines consistently differ in learning ability with proactive individuals displaying faster reward learning and reactive individuals exhibiting faster fear learning [31,35]. Other distinguishing characteristics that are consistent with a bold or shy phenotype being a trait of the line as opposed to a state are differences in escape-behavior mediated by caudal peduncle musculature, whole-brain neurotranscriptome profiles, and behavioral stress responses to anxiogenic and anxiolytic compounds [32–36,38–40]. In the current study we used fish that were selectively bred for 14 generations, raised in a common garden-like environment, and between 9-15 months old at time of testing. All fish had the same date of birth and sex was balanced across groups. Before testing, we housed the fish together in 40 L tanks and fish were fed twice a day with Tetramin Tropical Flakes (Tetra, USA). One week prior to testing we physically isolated fish into 3-liter tanks on a recirculating water system (Pentair Aquatic Eco-Systems or Aquaneering) using UV and solid filtration on a 14:10 L/D cycle (lights on between 06:00–20:00) and consistent water parameters (temperature of 27 °C, conductivity of 700 uS, and pH of 7.2). Fish had visual and olfactory access to each other and remained in this environment when not being tested for the duration of the experiment. Three days prior to testing, we withheld food from the fish to reduce the possibility of satiation while training.

### Conditioned place preference task

We followed a previously published conditioned place preference (CPP) protocol using the Zantiks automated behavioral research system (Zantiks, UK) [35]. In brief, we tested each fish in the CPP task that differed in number of conditioning days. To habituate each fish to the assay we placed the fish in the tank for 10 min with no training stimulus lights. After habituation, we determined the baseline preference for the light stimuli (grey or checkered pattern) for each fish during a baseline trial where both stimuli were presented for 10 minutes. We determined the conditioned and non-conditioned stimuli as the stimuli where the fish spent the least and most amount of time, respectively. During conditioning days, we sequentially presented each stimulus for 5 min to each fish. The non-conditioned stimulus was presented for the first five minutes followed by the conditioned stimulus. We administered 100 microliters of brine shrimp or distilled water every minute during the presentation of the conditioned stimulus for the treatment and control fish, respectively. We fed control fish an equivalent amount of brine shrimp after each conditioning trial. We tested if preference for conditioned stimulus changed the day after 3 conditioning days by administering a probe trial, which used the same parameters as described for the baseline trial.

We euthanized half of the fish after testing on the third conditioning day (3 total days of conditioning group) and the other half of the fish were euthanized after testing on the seventh conditioning day (7 total days of conditioning group). Sixty minutes after the conditioning trial we euthanized animals by decapitation to prepare brain samples for immunohistochemistry (described below). We selected tissue sampling times of 3 and 7 days based on a prior study demonstrating when bold and shy fish first showed significant changes in preference for conditioned stimulus from baseline, respectively [35]. This facilitated our investigation of neural activity at time points where we have seen each personality type show evidence of learning. We euthanized a third group of fish 60 minutes after receiving food to serve as a control to differentiate brain activity due to feeding in the CPP from brain activity due to learning. The location of conditioned stimulus (left or right side) was consistent within a fish for probe and baseline trials. To control for any bias towards the left or right, the location of conditioned stimulus during probe and baseline trials was cross balanced across groups so each group had an equal number of each stimulus order. We compared the time spent in the conditioned zone at baseline to the time spent in the conditioned zone in probe trials to assess learning. There were 10 total groups: bold and shy after 3 days of conditioning control, bold and shy after 3 days of conditioning treatment, bold and shy after 7 days of conditioning control, bold and shy after 7 days of conditioning treatment, and bold and shy food group. Each group had a sample size of 12 fish for a total of 120 fish, with equal number of males and females in each group. Sex was determined through visualization of ovaries or testes on dissection. All methods in this study were approved by the University of Nebraska at Omaha’s Institutional Animal Care and Use Committee.

### Immunohistochemistry and Image Analysis

After fish were tested in the CPP, we fixed heads in 4% paraformaldehyde and then submersed in 15% and then 30% glucose solutions to cryoprotect. We serial sectioned the heads of each fish on the coronal plane at 16 micrometers using a cryostat (Thermo Scientific, HM525 NX) and stored samples at - 80°C until immunohistochemistry. We performed double-labelled immunohistochemistry following standard protocols [41]. We quantified tyrosine hydroxylase (rate limiting step in dopamine synthesis) as a proxy for dopamine, directly labelled serotonin (5HT), and PS6 as a marker for neural activity [42,43]. In brief, we washed slides in PBS solution, boiled citric acid solution, and then rewashed in PBS. We blocked slides for 1 hour with normal goat serum and incubated with primary antibody solution for 40 hours at 4°C. Afterwards we washed in PBS, incubated with secondary antibody solution for 2 hours, rewashed in PBS, and counterstained with DAPI. The primary antibodies in the first series of slides were 1:500 anti-PS6 conjugated in rabbit (Cell Signaling, 2215) and 1:500 anti-tyrosine hydroxylase conjugated in mouse (Antibodies Incorporated 77-700). The secondary antibodies for the first series were 1:500 Alexa fluor 647 anti-rabbit and 1:500 Alexa fluor 555 anti-mouse, respectively (Thermofisher, US). In the second series of slides, we used 1:500 anti-PS6 conjugated in rabbit (Cell Signaling, 2215) and 1:500 anti-serotonin conjugated in rat (Sigma-Aldrich, MAB352) primary antibodies. The secondary antibodies for the second series were 1:500 Alexa fluor 488 anti-rabbit and 1:500 Alexa fluor 555 anti-rat respectively (Thermofisher, US). We submitted slides to the University of Nebraska Medical Center’s Advanced Microscopy Core Facility for whole-slide imaging using Zeiss Axioscan7 and scanned at 20X with CY5 (Alexa fluor 647), TRITC (Alexa fluor 555), and FITC (Alexa fluor 488) filters. We manually counted the number of positive cells of each protein and DAPI-stained cells using Zeiss ZEN Lite software (Carl Zeiss Microscopy GmbH, Jena, Germany) in the brain region of interest (ROI). We divided positive cell counts by the number of DAPI cells to get proportion of labelled cells within the ROI. The counter was blinded to the ID of each slide. Supplementary Table 2 shows the criteria and ROI box for each brain region. We counted the number of PS6 positive cells in the medial zone of the dorsal telencephalon (Dm), lateral zone of the dorsal telencephalon (Dl), posterior zone of the dorsal telencephalon (Dp), preoptic area (POA), habenula (Hb), ventral zone of the ventral telencephalon (Vv), superior zone of the ventral telencephalon (Vs), ventral hypothalamus (Hv), pretectum (PT), posterior tuberculum (Tpp), superior raphe (SR), and dorsal hypothalamus (Hd) using a zebrafish brain atlas (Wulliman et al., 1996). We counted tyrosine hydroxylase positive cells in the POA, Vv, Vs, PT and Tpp as these are all dopaminergic areas implicated in learning [20]. We counted serotonin positive cells in the SR and Hd because these regions are serotonergic and implicated in learning [22].

### Statistics

We performed a repeated measures ANOVA with a Whiteman’s adjustment for non-normal data of time (baseline or probe), treatment (brine shrimp conditioned or water control), personality type (bold or shy) and all interactions on time spent in the conditioned zone. Due to variations in normality of immunohistochemistry data, we used a quasibinomial generalized linear model. We conducted a quasibinomial generalized linear model to analyze the effects of treatment, time, and personality type and interactions on the proportion of active cells, active tyrosine hydroxylase, or active serotonin labelled cells for each relevant brain area. If there were significant interactions, we ran post hoc tests using emmeans. All statistics were run in R using RStudio [44]. The food only group was not included in the analyses as we did not see any statistical significance between the treatment group compared to controls. This group intended to serve as a control for food-induced increases in PS6, 5HT and TH amounts. However, with no treatment effect, inclusion of this group decreased statistical power. We used general linear mixed effect models in the quasibinomial family to investigate the effects of change in behavior (difference in time spent in the conditioned zone from baseline to probe), treatment, personality type, and time point and all interactions on proportion of active serotonin or dopamine neurons.

We performed structural equation modelling (SEM) of the active cells in the Dm, Dl, Dp, Hb, POA, Hv, and Hd to see if the interactions between brain regions differ across groups following previously established methods [45]. All fish with missing values in brain regions because of inability to accurately count cells (e.g., tissue folding, out of focus, damage to section) were excluded from SEM analyses. Therefore, we excluded the Vs, Vv, Tpp, SR, and PT due to too many fish missing values (over 5 fish) for these regions, which led to the SEM analysis not running. We used a maximum likelihood estimation with robust standard errors due to data not being normally distributed. In brief, first the data was split by control and treatment to allow for comparisons of proactive and reactive individuals within these groups. The treatment group was tested first and an optimal model was created. Then the model was grouped by personality type to compare differences across coping style in the treatment group. When the model was split, paths were constrained (made to be equal) if the change in chi-squared was less than 3.84. The same model was used for the control group and the same procedure was done to compare paths. The same procedure was done to compare within the proactive group across control and treatment and within the reactive group between control and treatment. Sample sizes for SEM analyses: Bold Control After 3 days of conditioning (N= 12) Bold Treatment After 3 days of conditioning (N = 12), Shy Control After 3 days of conditioning (N = 10) Shy Treat After 3 days of conditioning (N = 12) Bold Control After 7 days of conditioning (N= 11), Bold Treat After 7 days of conditioning (N= 10), Shy Control After 7 days of conditioning (N= 7) Shy Treat After 7 days of conditioning (N= 9). Models were tested using R lavaan package (Rosseel, 2012). After testing just control and treatment we split the data by probe and performed comparisons across personality type between control and treatment groups. Due to missing values, we excluded the Hb for these models.

## Results

### Significant relationships between changes in behavior and active serotonin and dopamine neurons

To directly investigate the influence of neurotransmitter activity and general activity on learning, we ran general linear mixed effect models to investigate the relationship between change in behavior and proportion of active neurons or active dopamine or serotonin neurons across groups. In the superior raphe (SR) there was a three-way interaction between treatment, personality type, and the change in behavior on the proportion of double labelled PS6 and serotonin neurons (b = -0.06, t = -2.26, p = 0.031) (Figure 2A). Post-hoc tests revealed that shy control (m =-0.027, p = 0.019), bold treatment (m = -0.019, p = 0.031), and shy treatment (m = -0.095, p<.001) all had significant negative slopes for the relationship between change in behavior and proportion of double labelled PS6 and serotonin neurons. Additionally, shy control has a more negative slope than shy treatment (b = 0.069, p = 0.020) and shy treatment had a more negative slope compared to bold treatment (b = 0.077, p = 0.005). In the posterior tuberculum (Tpp) there was a significant effect of change in behavior on the proportion of PS6 and TH double labelled neurons for all groups (b = 0.02, t = 2.45, p = 0.019) (Figure 2B). All other brain regions had no significant effects for change in behavior, treatment, personality type, and interactions on the proportion of double labelled PS6 and serotonin or TH neurons (Supplementary Tables 3). When investigating the main effects of time, personality, and treatment on the amount of time spent in the conditioned zone, the only significant main effect was time (F(1,88) = 12.11, p =.001). All groups significantly increased their time spent in the conditioned zone from baseline to probe.

**Figure 1.**
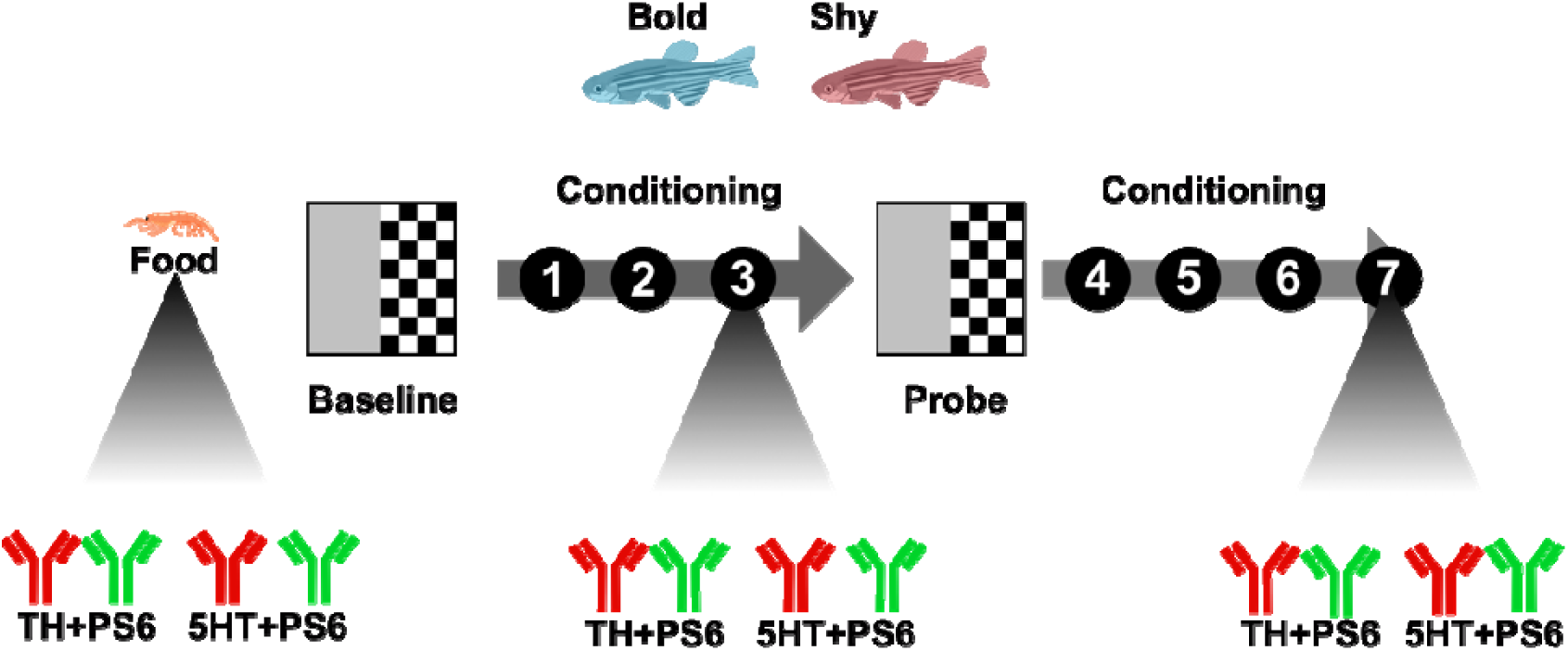
Representation of experimental design.

**Figure 2.**
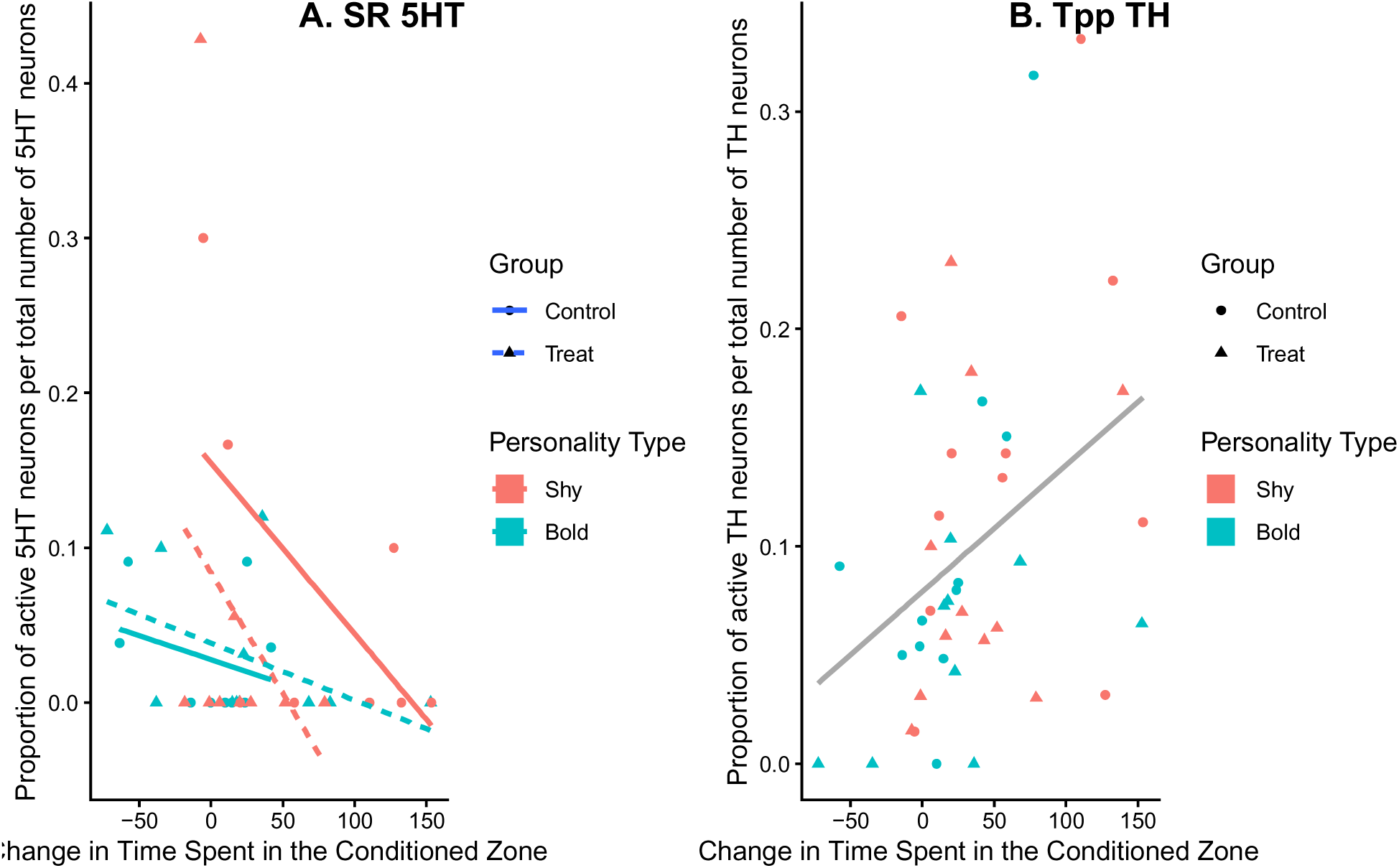
Relationship between change in time spent in the conditioned zone and proportion of active serotonin neurons in the superior raphe (A) and active TH neurons in the Tpp (B) by group. In (A) blue indicates bold, red indicates shy, dotted lines and circles indicate control, and solid lines and triangles indicate treatment. In (B) there are no differences between groups but blue indicates bold, red indicates shy, dotted lines indicate control, and solid lines indicate treatment.

### Differences in proportion of PS6 positive neurons in the Hd, and Dm across groups

In the dorsomedial telencephalon (Dm) there was a significant personality type by treatment interaction on the proportion of PS6 positive cells (b= -.054, t = -0.25, *p* = .049) (Figure 3A). Post-hoc tests revealed that shy treated fish had a lower proportion of PS6 positive cells compared to shy controls (b = 0.92, z = 2.00, p = 0.049). There was no significant difference in the number of active neurons in the Dm for bold treatment compared to control. There was a significant treatment by personality type interaction on the proportion of PS6 labelled neurons in the dorsal hypothalamus (Hd; b = 0.56, t = 2.13, p = 0.036) (Figure 3B). Post-hoc tests revealed that control bold fish had a greater proportion of PS6 labelled neurons compared to control shy fish (b = 0.36, z = 2.63, p = 0.042). All other brain regions had no significant effects for treatment, personality type, time point, and interactions on the proportion of PS6 labelled neurons (Supplementary Tables 2).

**Figure 3.**
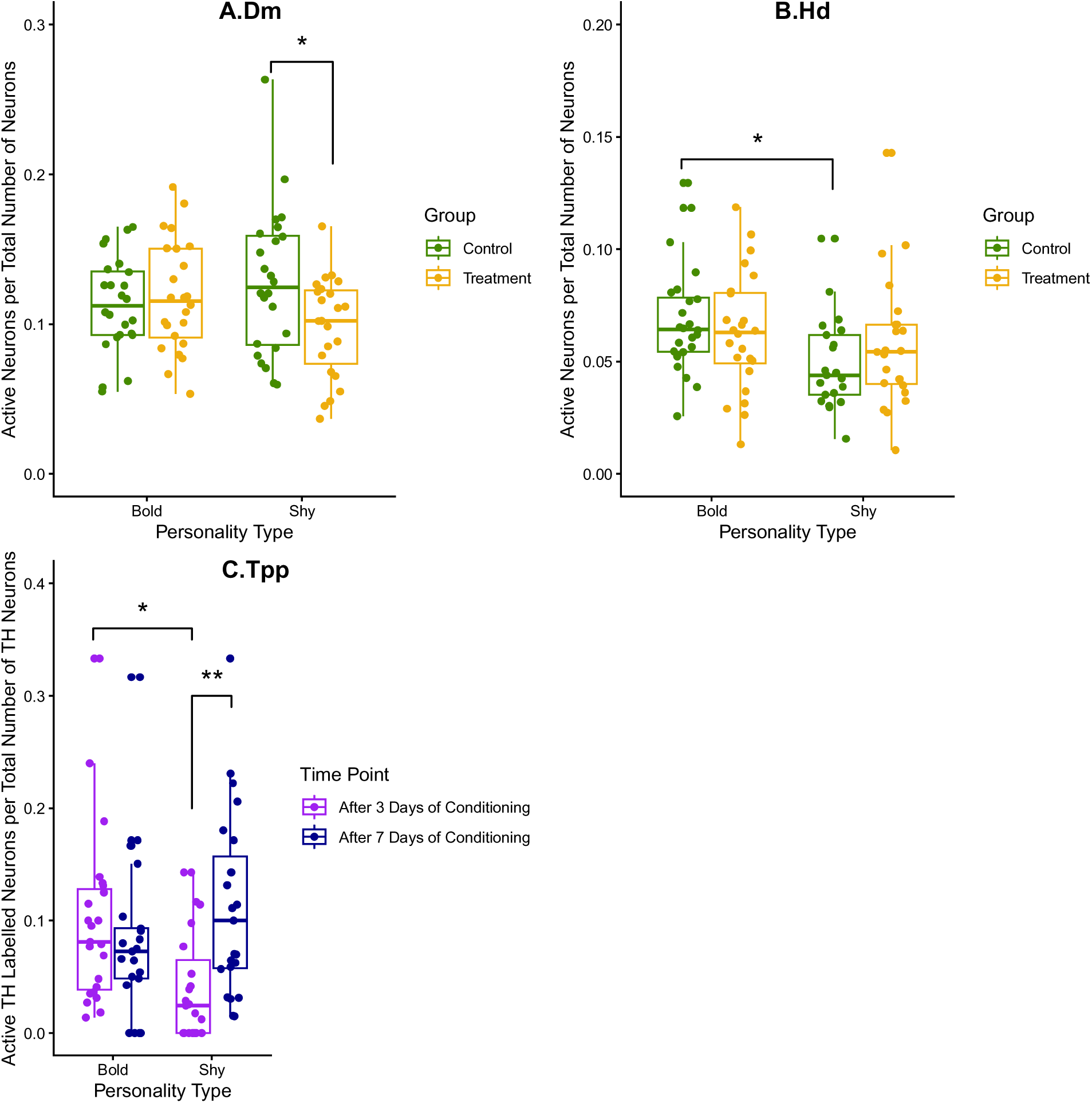
Proportion of active neurons per total neurons compared between groups for the Dm (A), Dl (B), and proportion of active dopamine neurons in the Tpp (C). In (A) and (B) green indicates control, orange indicates treatment. In (C) purple indicates after 3 days of conditioning and navy indicates after 7 days of conditioning Each dot represents one fish. Data is combined for probe 1 and probe 2 or control treatment when there are no differences between those groups. ** *p* <.01.

### Differences in proportion of double labelled TH and PS6 neurons in the Tpp

In the posterior tuberculum (Tpp) there was a significant main effect of personality type (b = - 1.09, t = 2.30, p = 0.024), and significant personality type by time point interaction effect (b = 1.40, t = 2.40, p = 0.019) on the proportion of double labelled TH and PS6 neurons (Figure 3C). Post-hoc tests revealed that all bold fish had a higher proportion of double labelled PS6 and TH neurons than all shy fish after 3 days of conditioning (b = 0.95, z = 2.90, p = 0.020). After 7 days of conditioning, all shy fish significantly increased proportion of double labeled TH and PS6 neurons relative to 3 days of conditioning in the Tpp (b = -1.08, z = -3.33, p = 0.005). All other brain regions had no significant effects for treatment, personality type, time point, and interactions on the proportion of double labelled PS6 and serotonin or TH neurons (Supplementary Tables 2).

### Network activity differs mainly between treatment bold and shy fish

There were only a few differences across groups in interactions between brain areas (Figure 4). We compared across bold and shy groups in the after 3 days of conditioning control, after 3 days of conditioning treatment, after 7 days of conditioning control and after 7 days of conditioning treatment groups. In the after 3 days of conditioning control group there was only one difference between bold and shy fish where there was a negative relationship between the POA and Hd in the shy group and no relationship in the bold group. Fit statistics for this group was acceptable (2(22) = 11.301, p = .34, CFI =0.94, RMSEA = .10, SRMR = .12). There were no differences between bold and shy fish in interactions in the treatment after 3 days of conditioning group and fit statistics for this group was acceptable (2(18) = 8.22 p = .51, CFI =0.99, RMSEA = .10, SRMR = .11). After 7 days of conditioning there were two differences in the control group between bold and shy fish. There was a positive relationship between the Dm and Dl in bold fish that was not present in shy fish. There was also a positive relationship between the POA and Hv in shy fish that was not present in bold fish. Fit statistics for this group was acceptable (/2(24) = 8.582, p = .66, CFI =0.99, RMSEA = .10, SRMR = .10). There were two differences between personality type in the treatment group after 7 days of conditioning. There was a positive relationship between the Dp and Dl in the bold fish that was negative in the shy fish. There was also a stronger connection between the Dp and Hd in shy fish compared to bold fish. Fit statistics for this group was acceptable (/2(19) =4.48 p = .87, CFI =0.99, RMSEA = .08, SRMR = .07).

**Figure 4.**
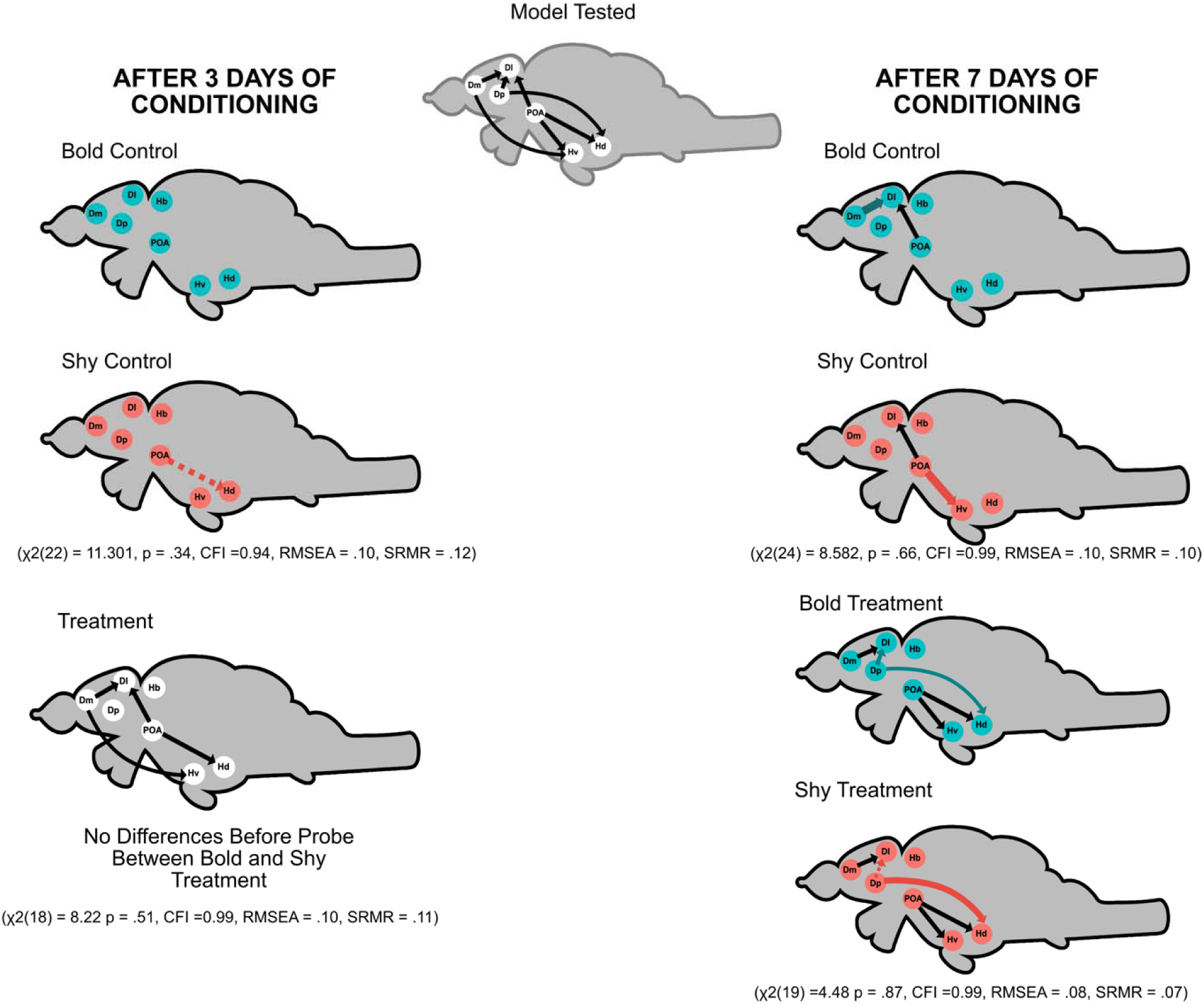
Models of functional connectivity between bold and shy fish at different time points and groups. Red circles indicate shy fish, teal circles indicate bold fish. White circles indicate no differences between groups. Line size indicates strength of connection determined by B value. Colored lines indicate differences across groups. Black lines indicate no difference across group. Solid lines indicate a positive connection and dotted lines indicate a negative interaction.

## Discussion

Individual differences in learning are associated with differences in neural and neurotransmitter activity and personality type. The neural mechanisms by which personality type and learning abilities are related are not well understood. We investigated differences in serotonin, dopamine, and general activity patterns between bold and shy zebrafish while learning. We found that there were no differences in dopamine or serotonin activity (double labelled TH or serotonin with PS6), or general activity (PS6 positive) that explained differences in learning speed between personality types. However, investigating interactions between brain regions revealed differences associated with learning in the teleost homolog of the mammalian olfactory cortex (Dp).

The behavioral results indicate that all fish increased time spent in the conditioned zone (Supplementary Table 4). In contrast to a prior study, the current study showed that after 3 days of conditioning there were no differences in learning between personality types [35]. Additionally there were no differences between control and treatment, suggesting that learning did not occur by this time point. The inconsistency with prior study could be due to smaller samples size and fewer probe trials in this study, which resulted in lower power in the statistical test. Supporting this, when analyzing only the first probe for the data from Corcoran et al., (2025), the only significant effects were the effect of trial and trial by treatment (Supplementary Table 5). It is also possible that none of the fish learned at this time point due to individual differences across our groups. However, the trend of bold individuals learning faster has been seen in multiple studies [3–9].

We investigated the relationship between the change in time spent in the conditioned zone from baseline to probe and the proportion of active cells to determine potential activity related to learning (Figure 2). In the superior raphe fish that increased time spent in the conditioned zone after 3 days of conditioning had decreased serotonin activity in every group except for bold control. The superior raphe is the major serotonergic area in the brain and other studies similarly find a decrease in serotonin activity in the superior raphe associated with learning [21]. However, as this trend was also observed in shy control group, it suggests that learning does not solely drive this relationship with serotonin activity. In the posterior tuberculum, fish that increased time in the conditioned zone had more active dopamine neurons. The posterior tuberculum is the mammalian ventral tegmental area (VTA) analog that acts as a relay between sensory and motor brain regions, which suggests this relationship could simply be related to navigating the sensory environment [46]. Since this relationship is in control fish as well as treatment fish, it is unlikely directly related to learning. In the posterior tuberculum we also saw a difference in active dopamine neurons across groups (Figure 3C). The shy fish after 3 days of conditioning had a decrease in active dopamine neurons compared to all other groups. The decrease in activity in shy fish could be due to an initial delay in learning in shy fish, however as this trend is also in the control fish this may not be the only explanation. Another possibility is that this could indicate an increase in stress suppressing other processing in all of the shy fish. Interestingly, a study on social stress found that dominant zebrafish had more posterior tuberculum activity than subordinates [48]. This could be due to the dominant status or increased stress in the subordinate fish, supported by the fact that subordinate fish had an increased startle response. Our results are not clearly indicative of a difference in dopamine or serotonin activity caused by learning, suggesting that activity of both neurotransmitters are not directly related to learning differences between personality types. We cannot rule out the possibility that effects related to learning are at the level of the receptor as opposed to neurotransmitter activity [49]. Overall, despite observing some linear relationships between serotonergic and dopaminergic activity and behavior in select regions, these relationships do not explain differences in learning speed between personality type.

While there were differences in general neural (number of PS6 cells) activity in some brain regions across groups, these differences were not directly related to learning (Figure 3A-B). This was an unexpected result that might be a consequence of comparing averages between groups, which can mask individual level brain-behavior relationship effects [50]. Additionally this could be due to the method of identifying active neurons as we have previously observed differences in general neural activity (via quantification of immediate early gene expression) in the amygdala (Dm), hippocampus (Dl) and bed nucleus of the stria terminalis (Vs) between bold and shy fish while undergoing fear conditioning [31]. We did see some differences in PS6 activity that were not related to learning. Despite receiving a food reward, there was a decrease in active neurons in the amygdala (Dm) for the shy treatment group compared to shy controls. As we do not see significant differences between bold and shy treatment, it does not appear that the amygdala activity reflects a differing response to reward across bold and shy fish. In the dorsal hypothalamus, control bold fish had more active neurons than control shy fish. This area is primarily involved in sensorimotor activity and could be related to inherent differences in movement across lines [11,51].

After not observing differences related to learning in discrete brain regions, we investigated changes in network interactions involving seven brain regions across groups to assess for personality differences related to learning. In the treatment group there were generally more interactions compared to the control group, suggesting that there is more coordinated activity in these regions that is increasing with learning. Within the control group there were some differences in interactions in brain regions in control fish across personality type at different time points (Figure 4). Shy fish in both time points (after 3 days and after 7 days of conditioning) had a connection from the preoptic area to areas of the hypothalamus. Prior studies show that communication between the POA, ventral telencephalon and hypothalamic areas are linked to stress [52,53]. As individuals with a shy personality type show increased behavior and physiological reactivity to stress, this may explain the presence of shy-specific connections between the POA and hypothalamus. The bold control fish had no connections between brain areas after 3 days of conditioning but a connection from the amygdala (Dm) to the hippocampus (Dl) appeared after 7 days of conditioning. Communication between amygdala and hippocampal regions are associated with increased stress [52]. The appearance of the POA-hypothalamus and Dm-Dl connections in the control groups suggest that the CPP assay may be causing a stress response. Analysis of total distance swam, which is negatively correlated with stress, show that control fish decreased distance swam after 7 days of conditioning [54–57] (Supplementary Table 6). Overall, there was minimal activity in control groups and the interactions that did appear may be linked to stress.

We observed differences in coordinated brain activity between personality types in the learning group after 7 days of conditioning. The treatment group after 7 days of conditioning had a difference in connection from the olfactory cortex (Dp) to the hippocampus (Dl) with an opposite relationship between personality types. The bold fish had a positive interaction between the olfactory cortex and hippocampus while the shy fish had a negative interaction. There are few direct anatomical connections from the olfactory cortex to the hippocampus so this interaction is likely facilitated by another region [58]. The olfactory cortex in zebrafish contains an odor valence map that changes with conditioning [59]. We hypothesize that the opposing interaction between olfactory cortex to the hippocampal homologs between personality types is a result of differences in stimulus valence processing related to learning. A study in pigs showed that increased freezing is associated with fewer rewards eaten and less interactions with rewards, which suggests a decrease in reward sensitivity for shy animals potentially due to increased neophobia [60]. We also observed a stronger connection from the olfactory cortex to dorsal hypothalamus in shy treated animals relative to bold treated animals. There are orexin neurons that go from the olfactory cortex to the dorsal hypothalamus that are associated with odor attraction in mammals, although in zebrafish this connections are mainly from the hypothalamus to the olfactory cortex [61,62]. The increased strength in connection between olfactory cortex and dorsal hypothalamus in shy fish is interesting but could indicate that the shy fish had taken longer to eat the food. The fish were only fed during conditioning and we have previously shown that shy fish will take longer to eat when exposed to an acute stressor [32]. Therefore, the shy fish may not have eaten as much food due to a stressor (e.g., handling and novel environment) that resulted in increased hunger at the later time point.

Overall, there are no clear differences that explain variation in learning due to the neurotransmitters we investigated (dopamine and serotonin). This implies that while there have been differences in expression of dopamine and serotonin receptors and gene expression in general between bold and shy animals [12–14], the activity of these neurotransmitters may not explain variation in learning speed. For general neural activity patterns a key finding related learning differences between personality types is the interaction between the Dp to the Dl. We speculate that that bold and shy fish process the valence of food stimuli in distinct ways involving a pathway between Dp and Dl. Additionally, we do not see differences in general network activity connectivity after 3 days of conditioning, implying that distinct neural activity patterns related to learning form later in the learning process. Overall, results from our study suggest that differences in interactions between brain regions, likely due to distinct reward processing, explains variation in learning speed across personality types.

## Acknowledgments

We thank Amanda Hostert, Brandon Wolfsohn, and Beth Trail for help with fish husbandry. We are grateful to Madison Thurber and Brooklyn Schmidt for helpful discussions and technical assistance. We acknowledge use of the University of Nebraska Medical Center - UNMC Advanced Microscopy Core Facility, RRID:SCR_022467, P20 GM103427 (NIGMS, NE-INBRE), P30 GM106397 (NIGMS, NCS), P20GM130447 (NIGMS, CoNDA), P30 CA036727 (NCI, Buffett Cancer Center), S10RR02730 (NIH), S10OD030486 (NIH), Nebraska Research Initiative, UNMC Vice Chancellor for Research Office. We additionally thank James R. Talaska, B.S of the UNMC AMCF for assistance with whole-slide imaging. This project was funded by the National Science Foundation (IOS-1942202 to RYW) and University of Nebraska at Omaha (Graduate Research and Creative Activity grant to JC).

