## Supplementary Information for "Differences in interactions between brain regions across personality types while learning in zebrafish (*Danio rerio*)"

Jamie Corcoran^1^

Ryan Y. Wong^1,2^

1.	Department of Psychology, University of Nebraska at Omaha, Omaha, NE

2.	Department of Biology, University of Nebraska at Omaha, Omaha, NE

*Ryan Y. Wong

**Table 1. ROI Information for cell counts**

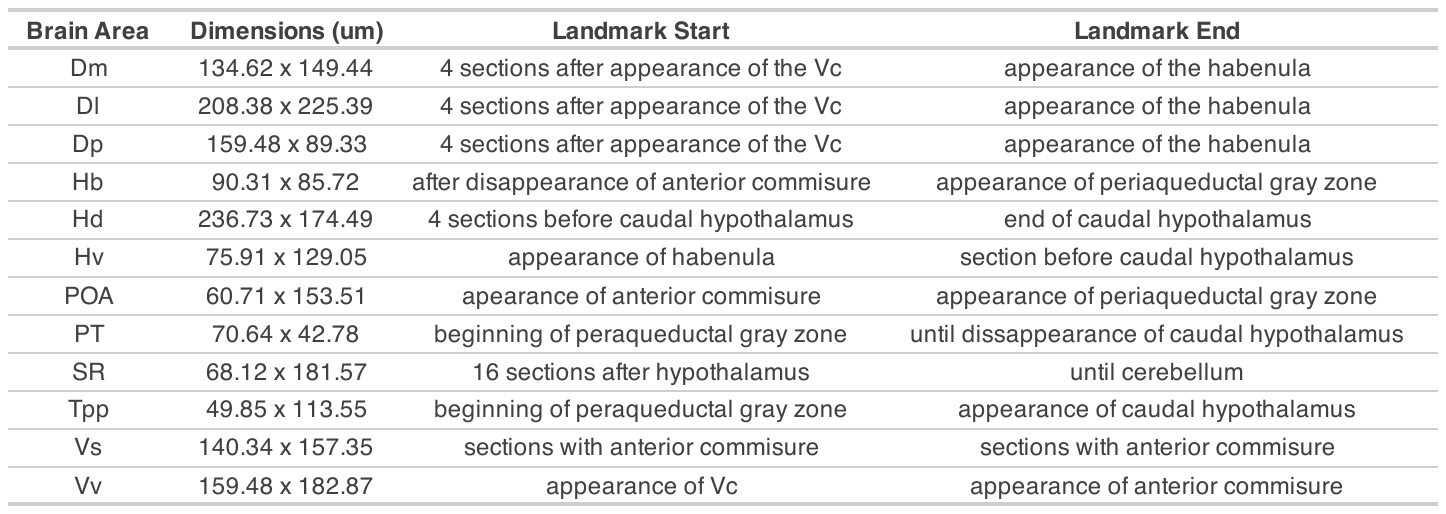

**Table 2. Models of the effects of treatment, personality type, time point, and interactions on the proportion of active cells, active dopamine, and active serotonin cells**

A. Active Dm neurons (ps6 labelled)

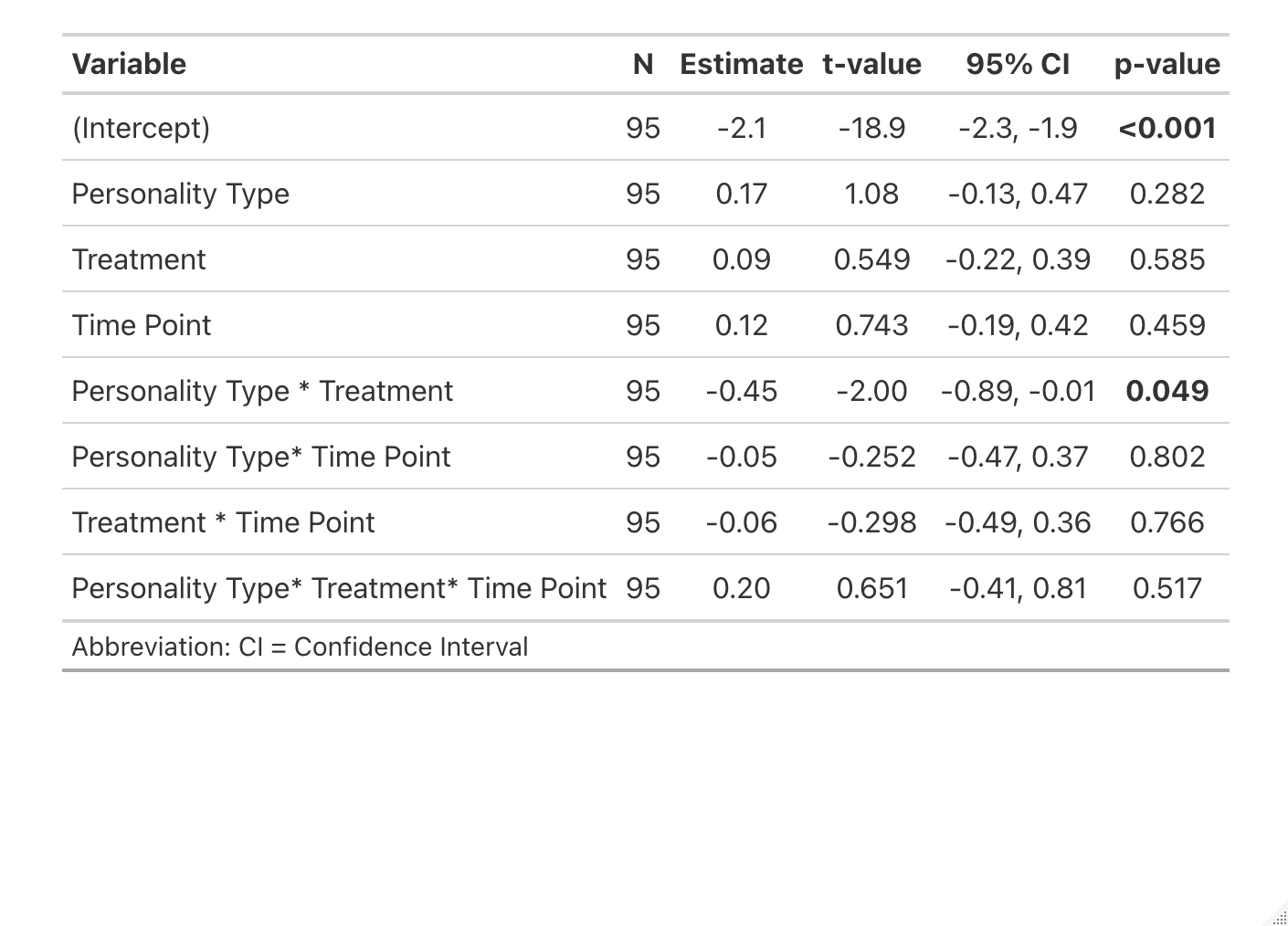

B. Active Dl neurons (ps6 labelled)
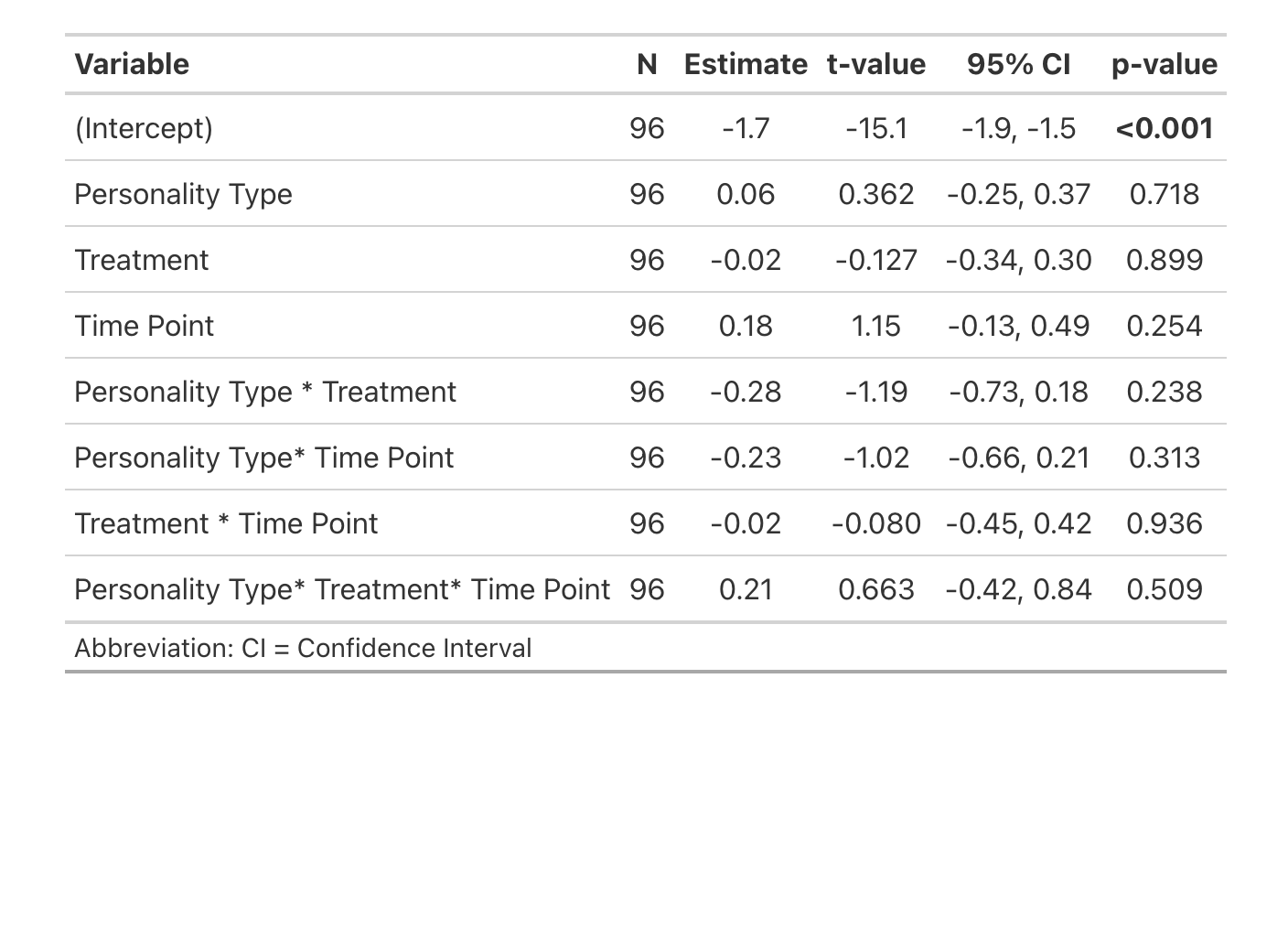

C. Active Dp neurons (ps6 labelled)

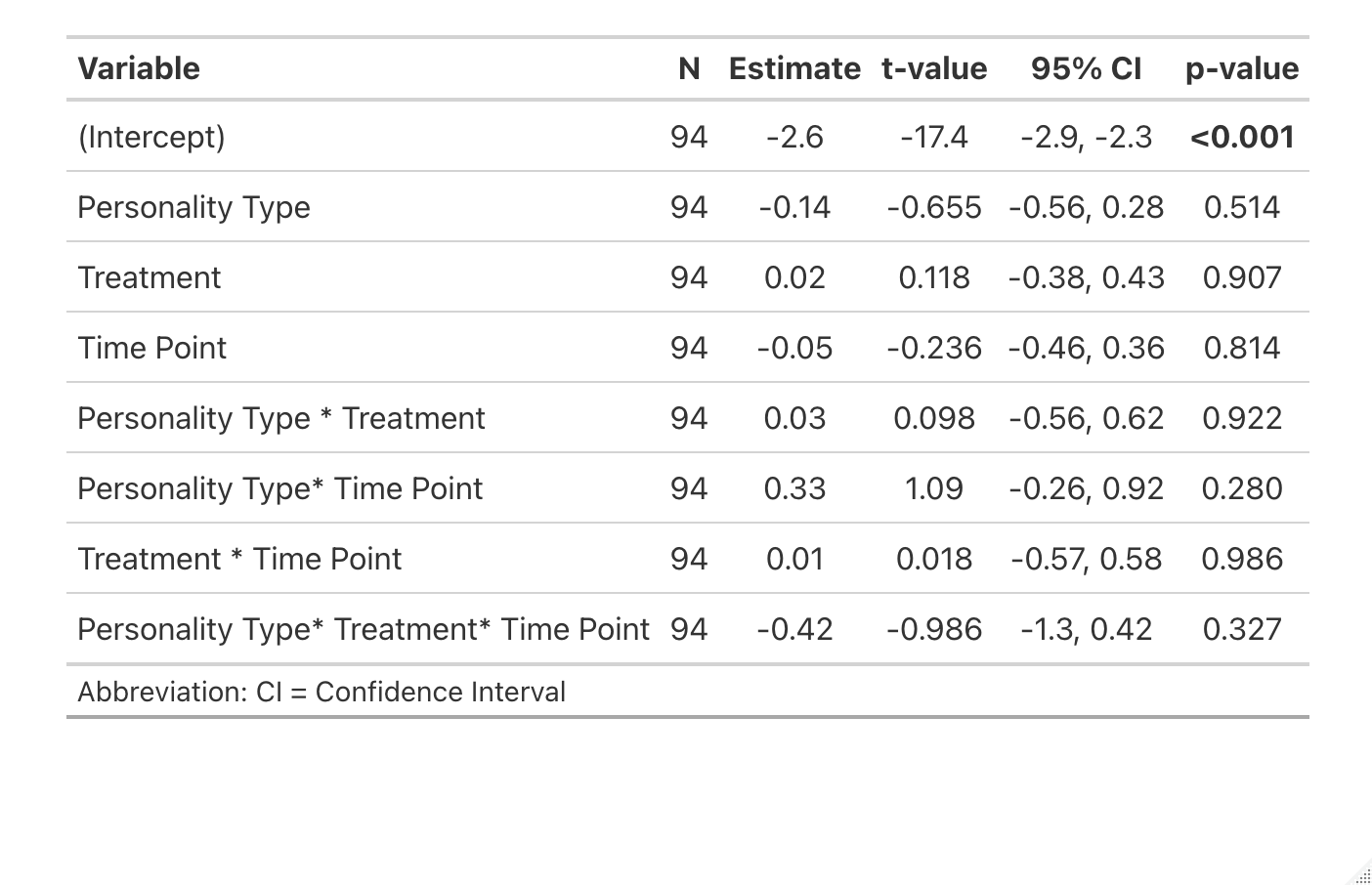

D. Active Hb neurons (ps6 labelled)
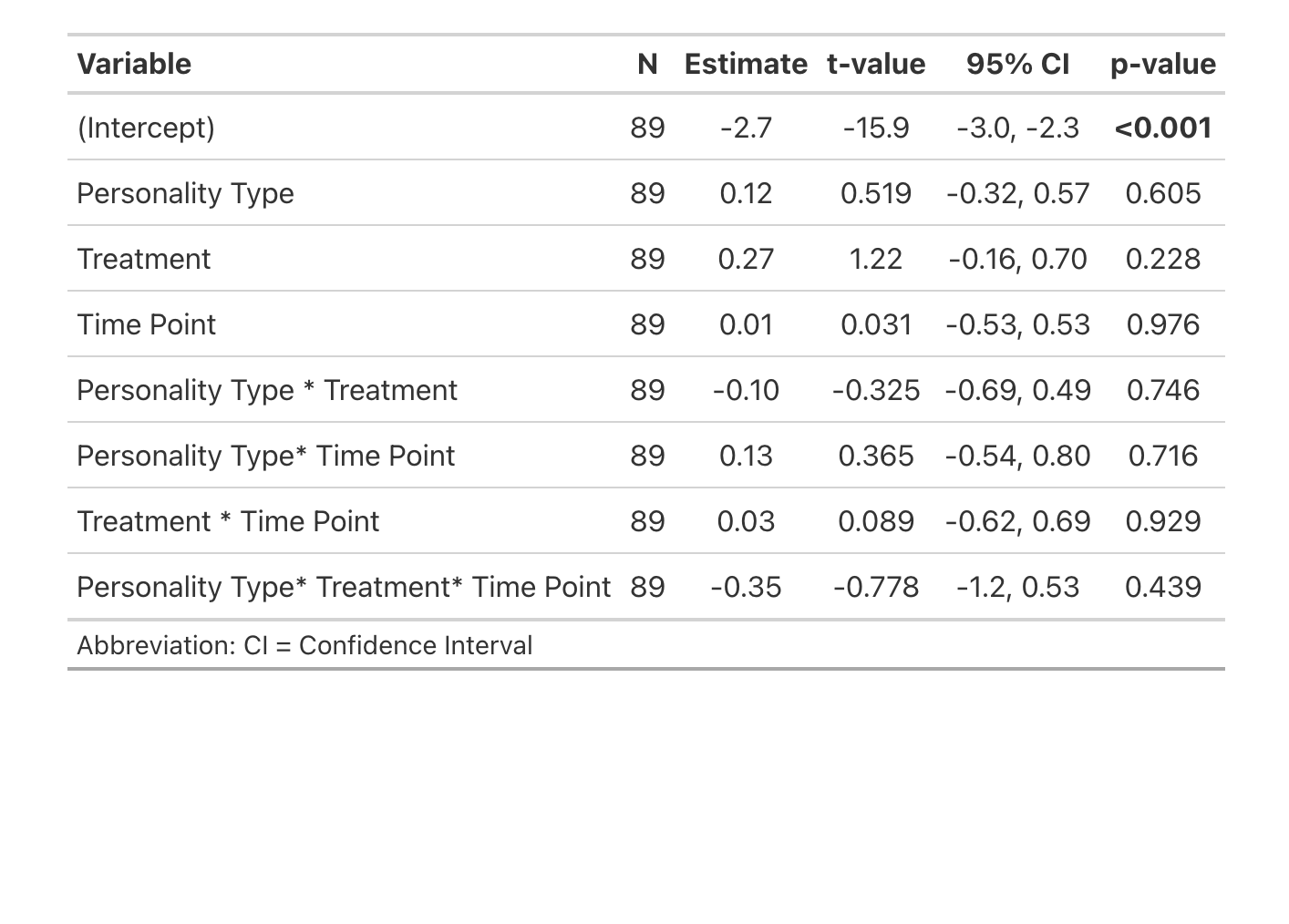

E. Active Hd neurons (ps6 labelled)
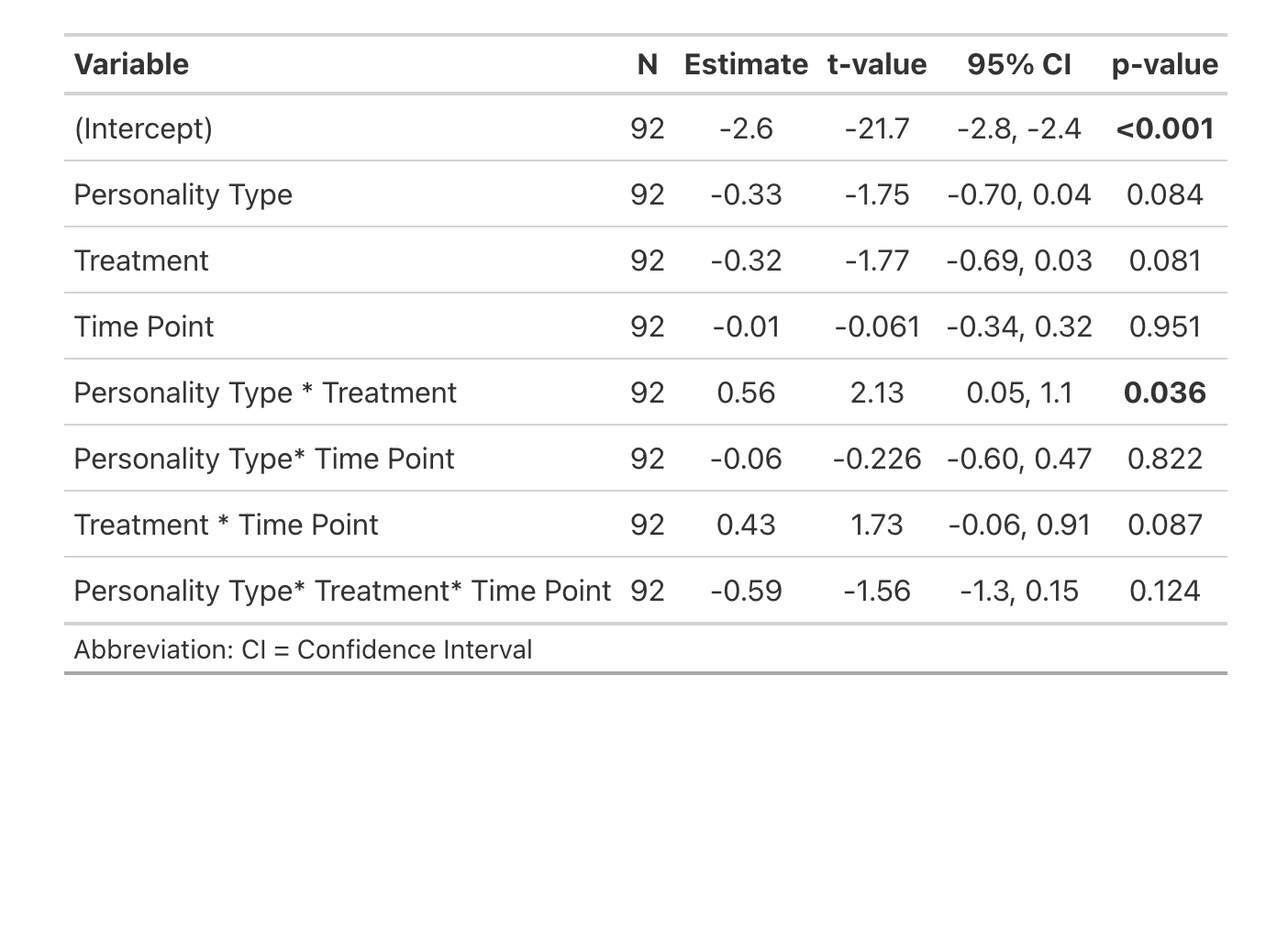

F. Active serotonin Hd neurons (ps6 + 5HT double labelled)
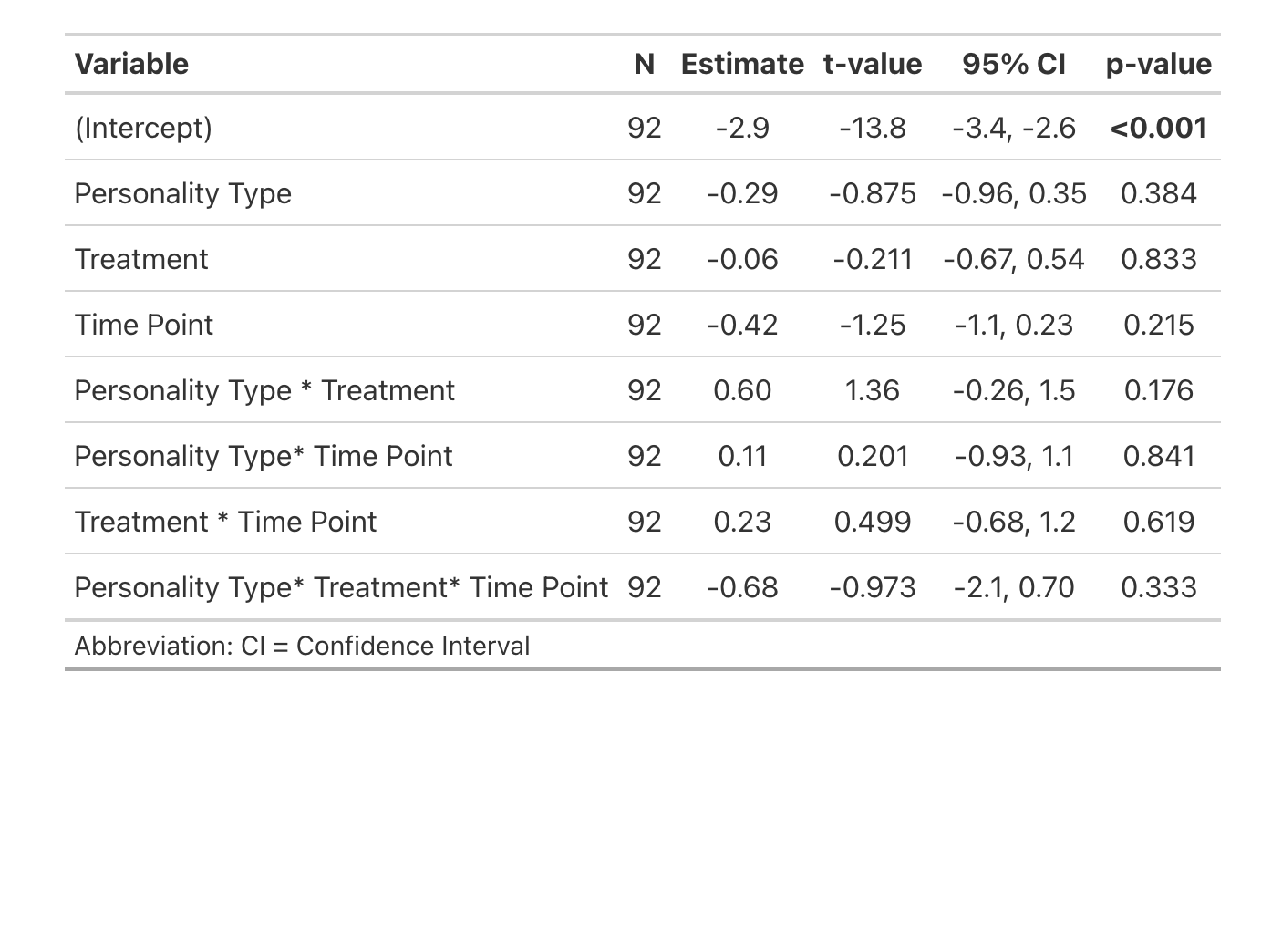

G. Active Hv neurons (ps6 labelled)
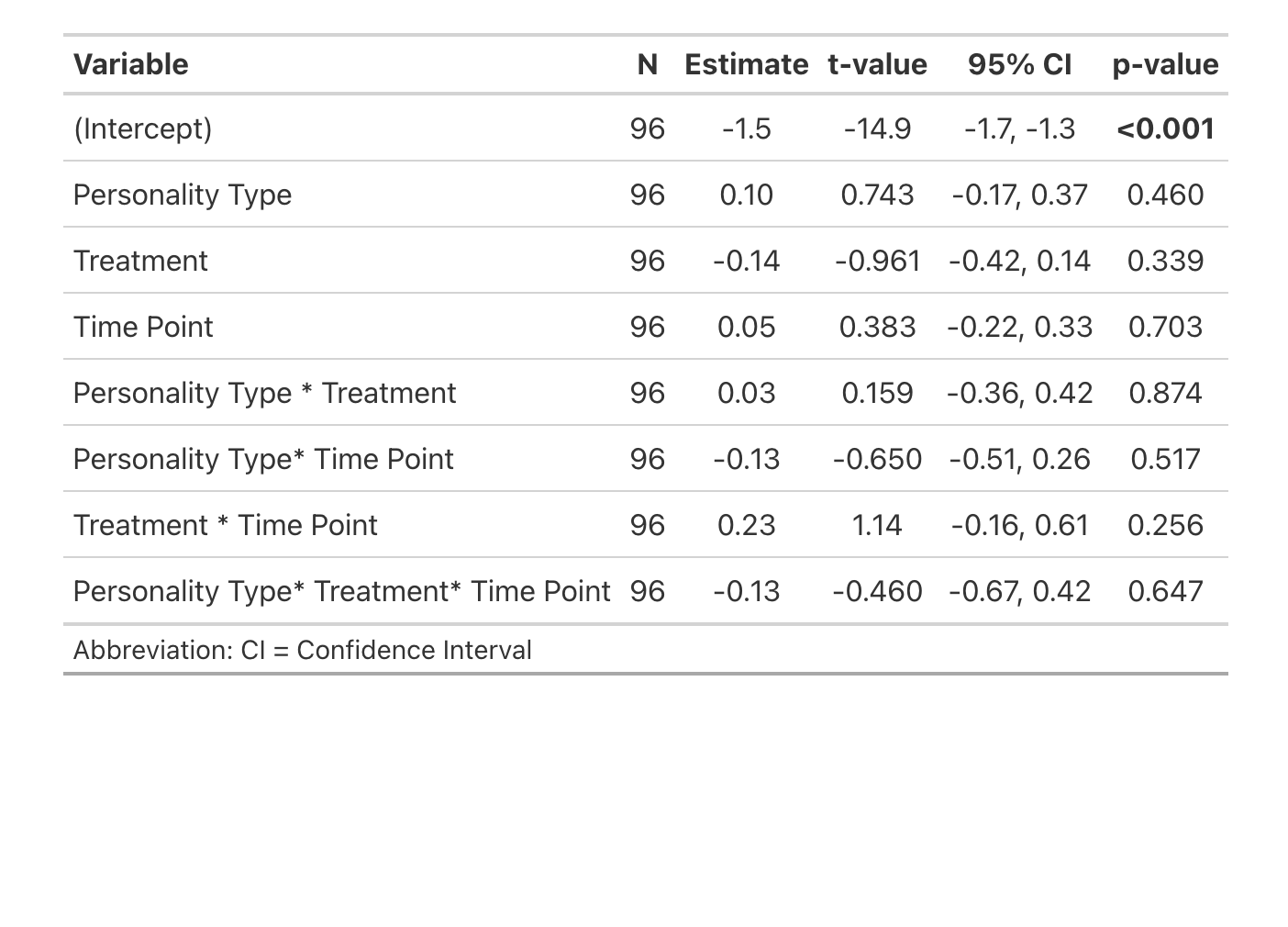

H. Active POA neurons (ps6 labelled)
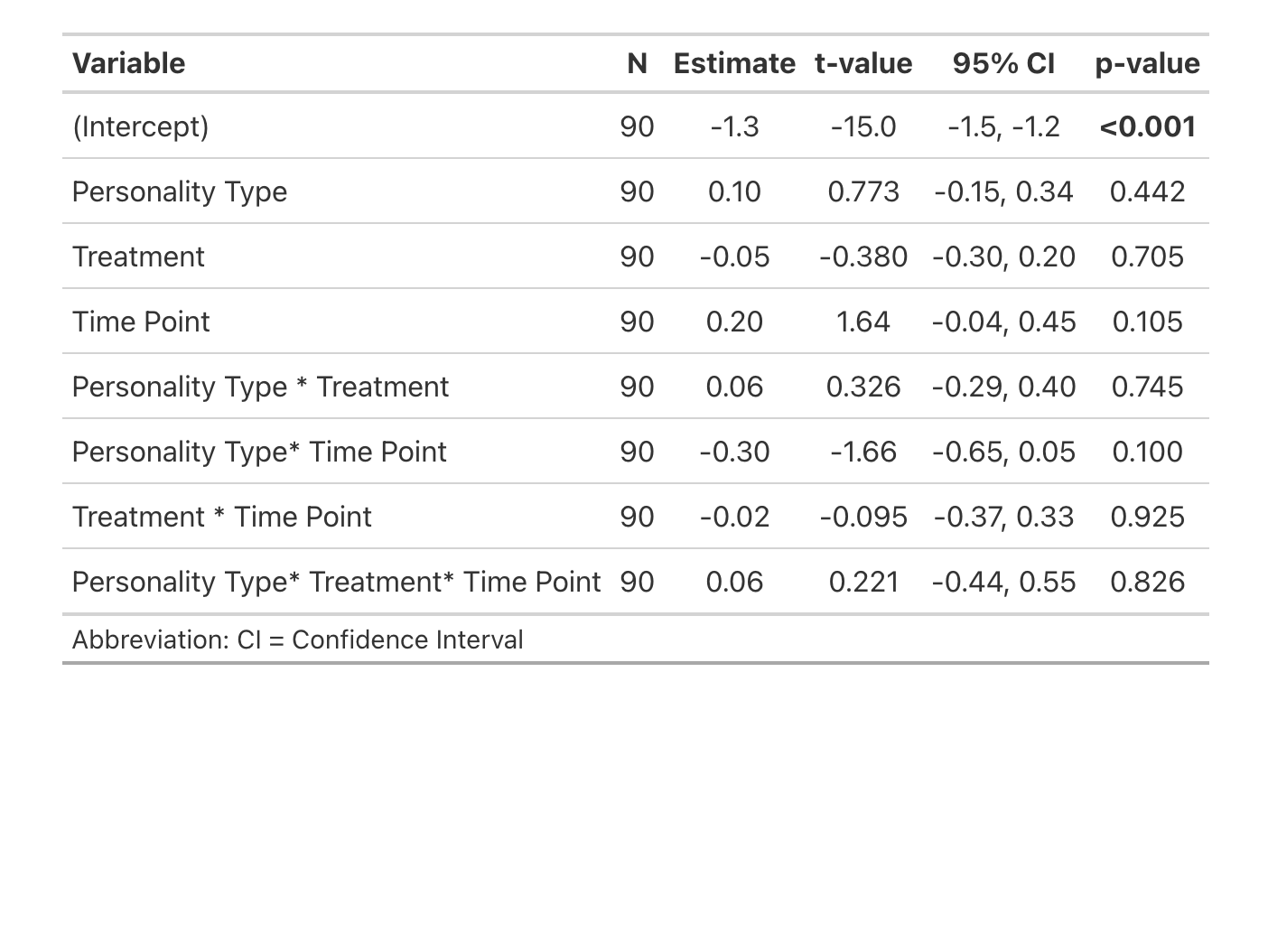

I. Active dopaminergic POA neurons (ps6 + TH double labelled)
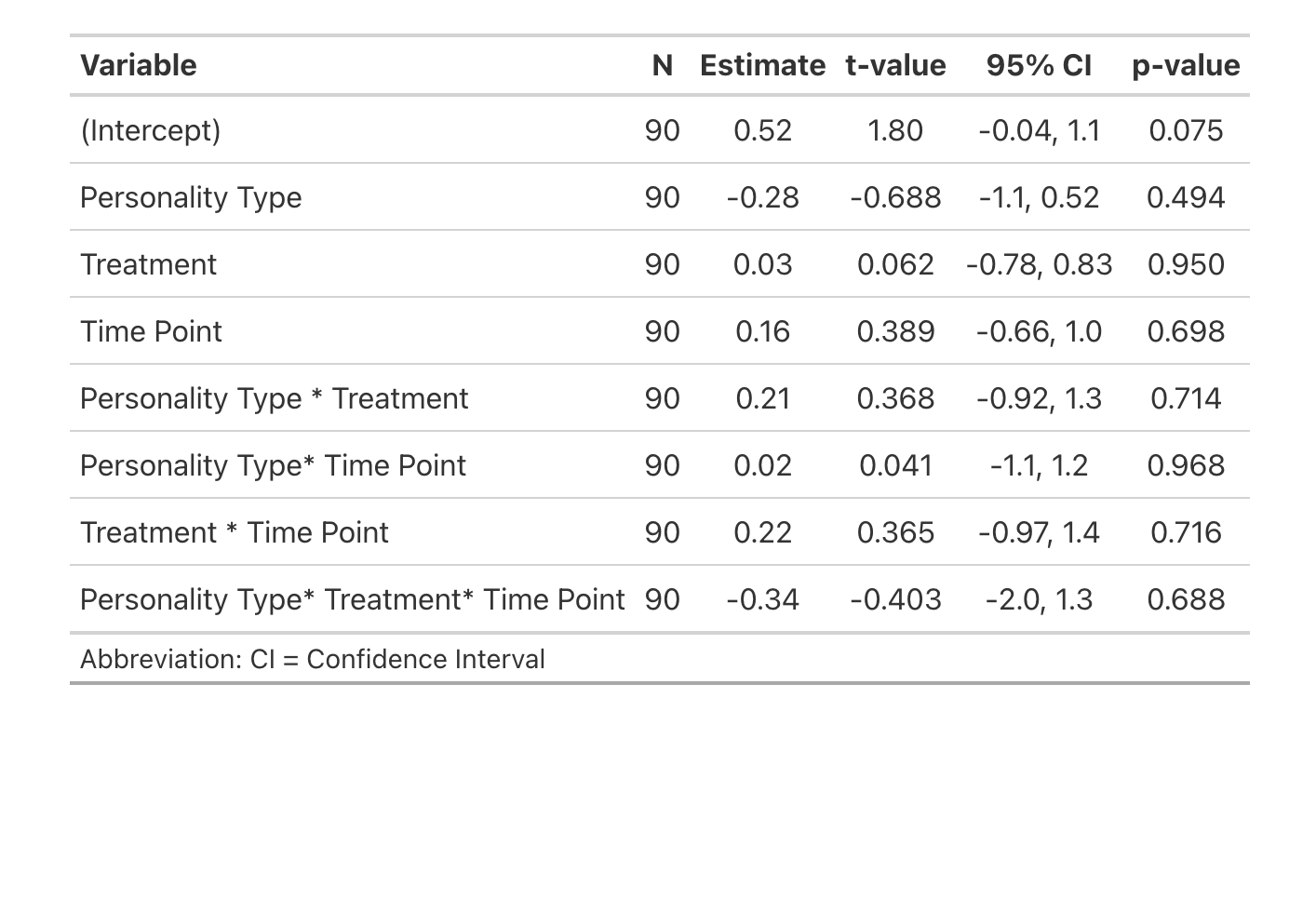

J. Active PT neurons (ps6 labelled)
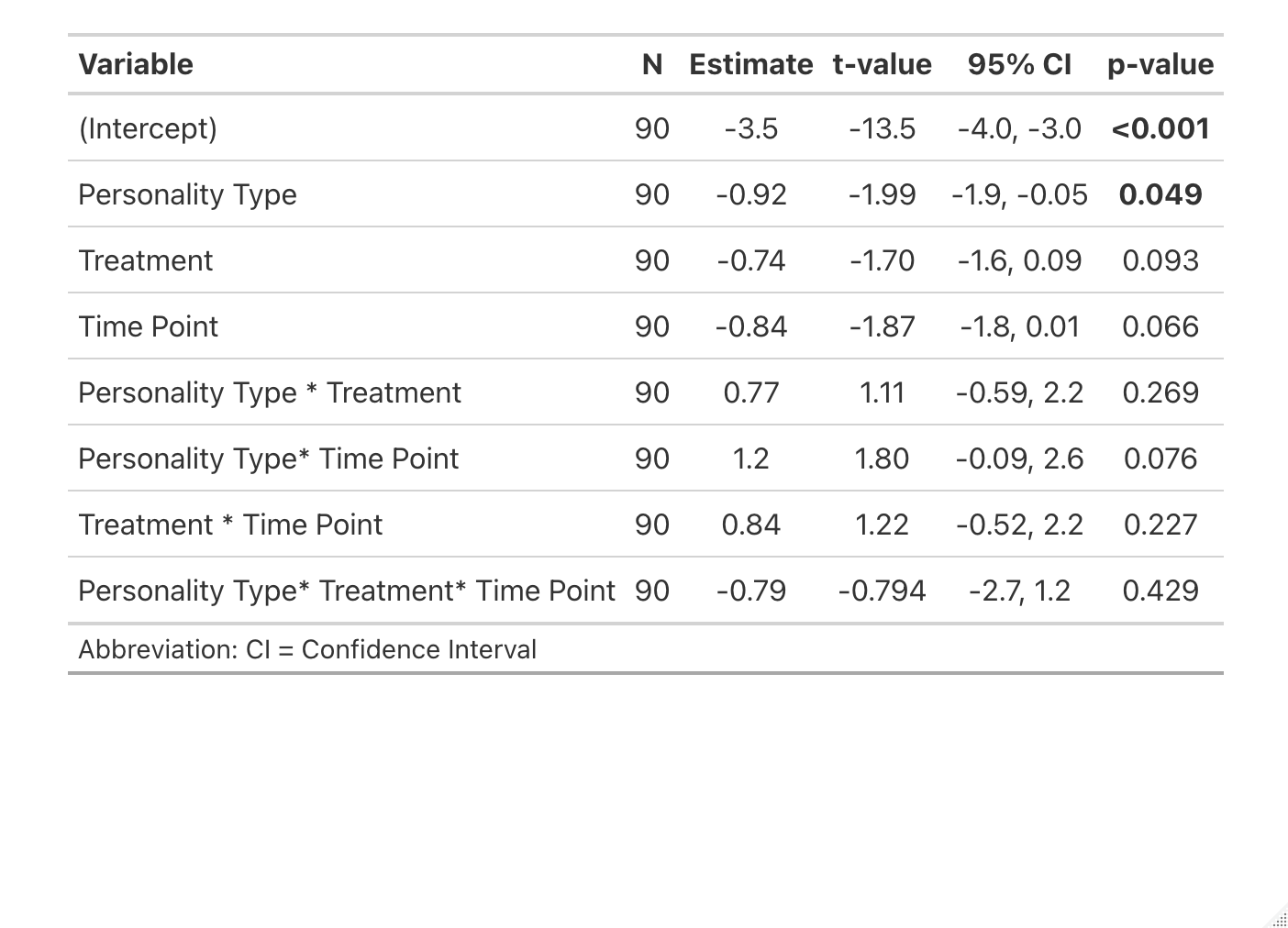

K. Active dopaminergic PT neurons (ps6 + TH double labelled)
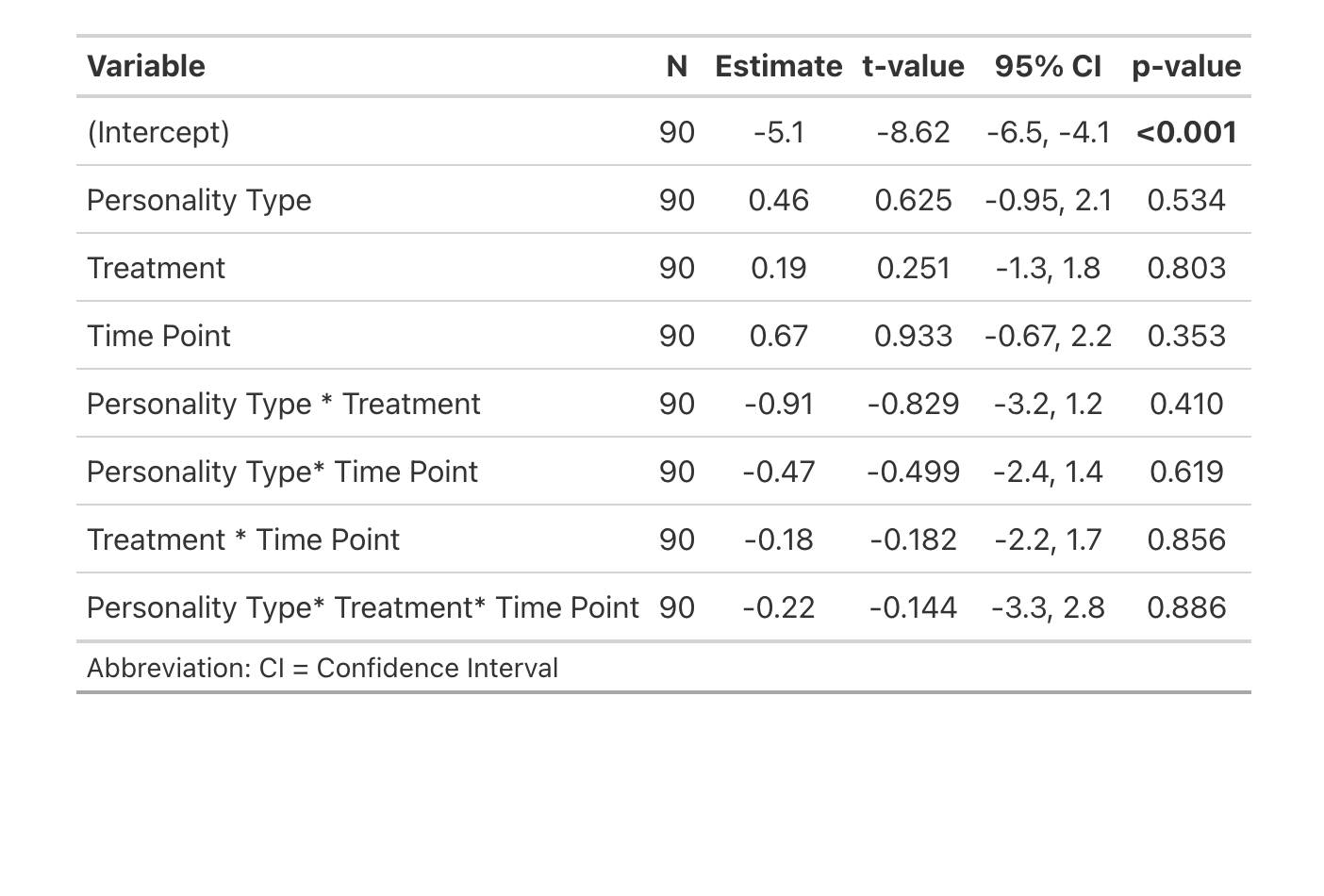

L. Active SR neurons (ps6 labelled)
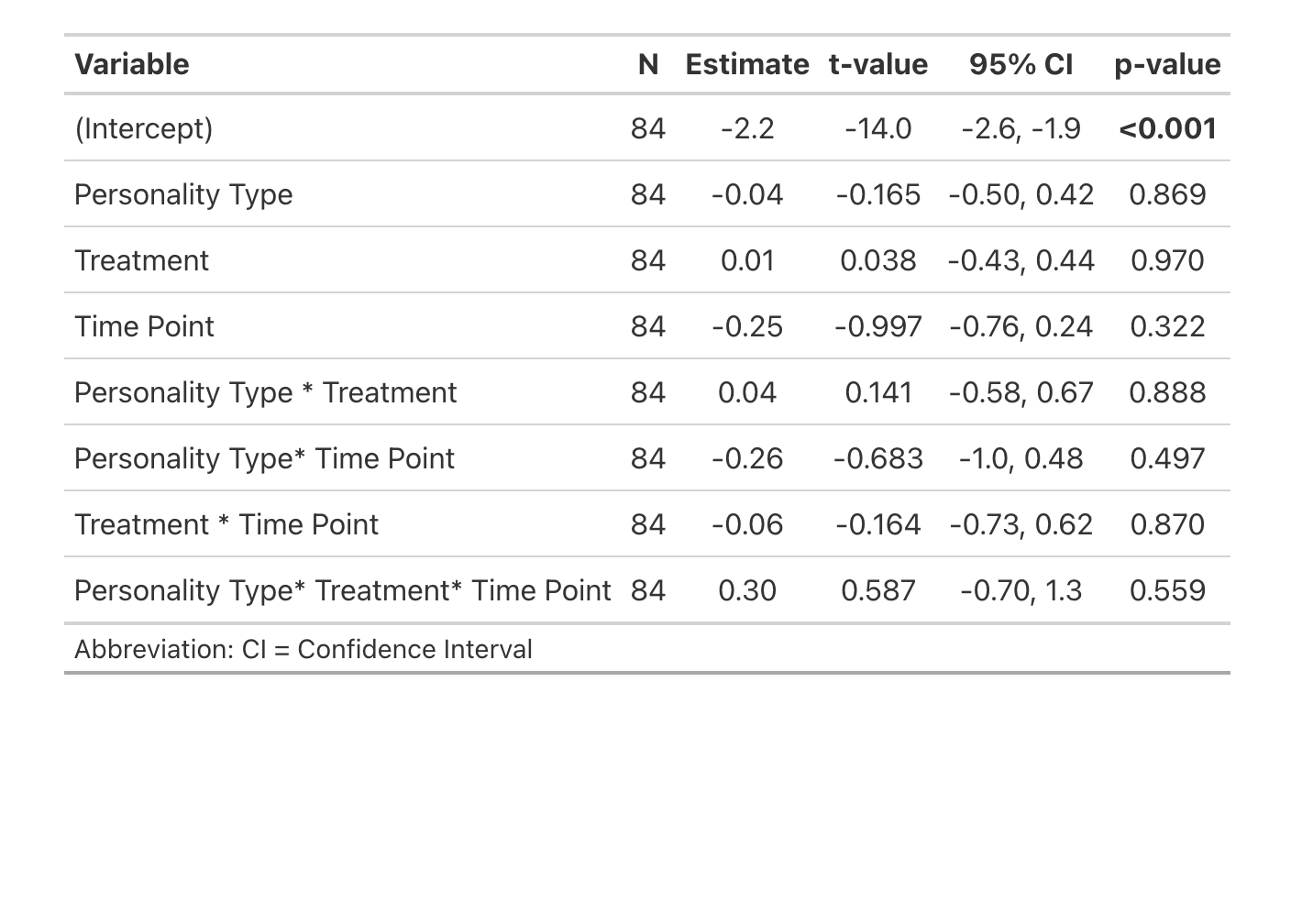

M. Active serotonergic SR neurons (ps6 + 5HT double labelled)
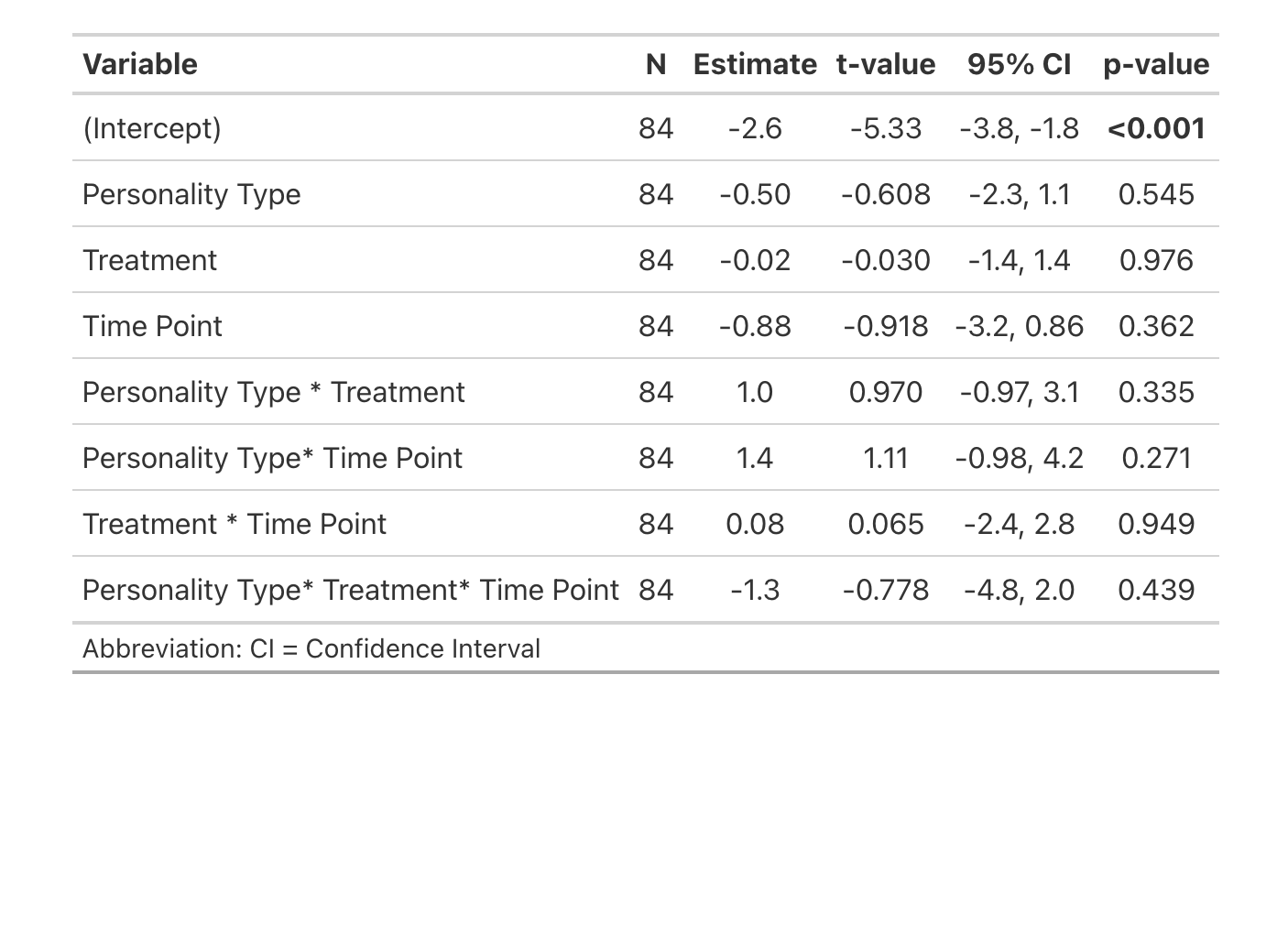

N. Active Tpp neurons (ps6 labelled)
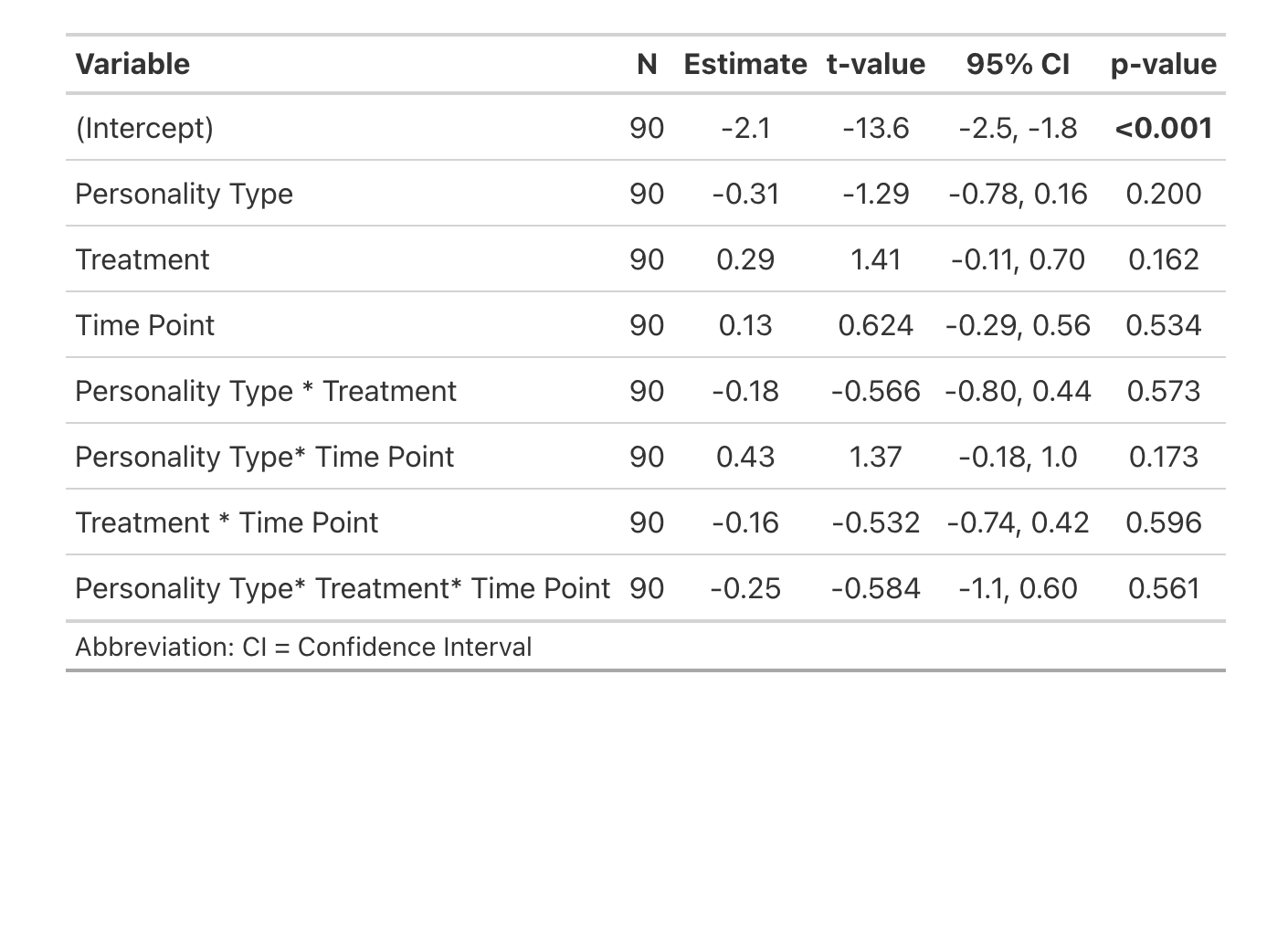

O. Active dopaminergic Tpp neurons (ps6 + TH double labelled)
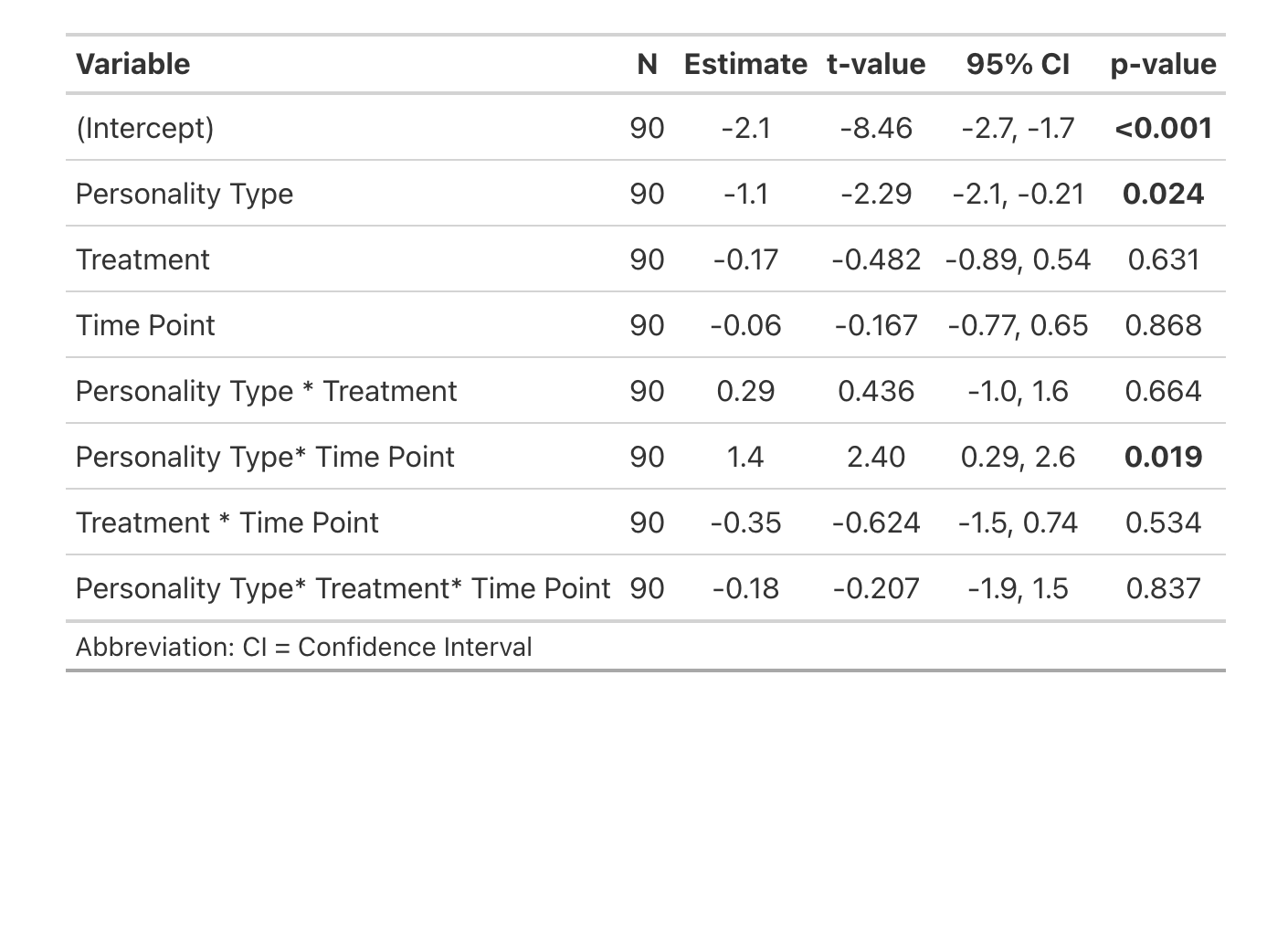

P. Active Vs neurons (ps6 labelled)
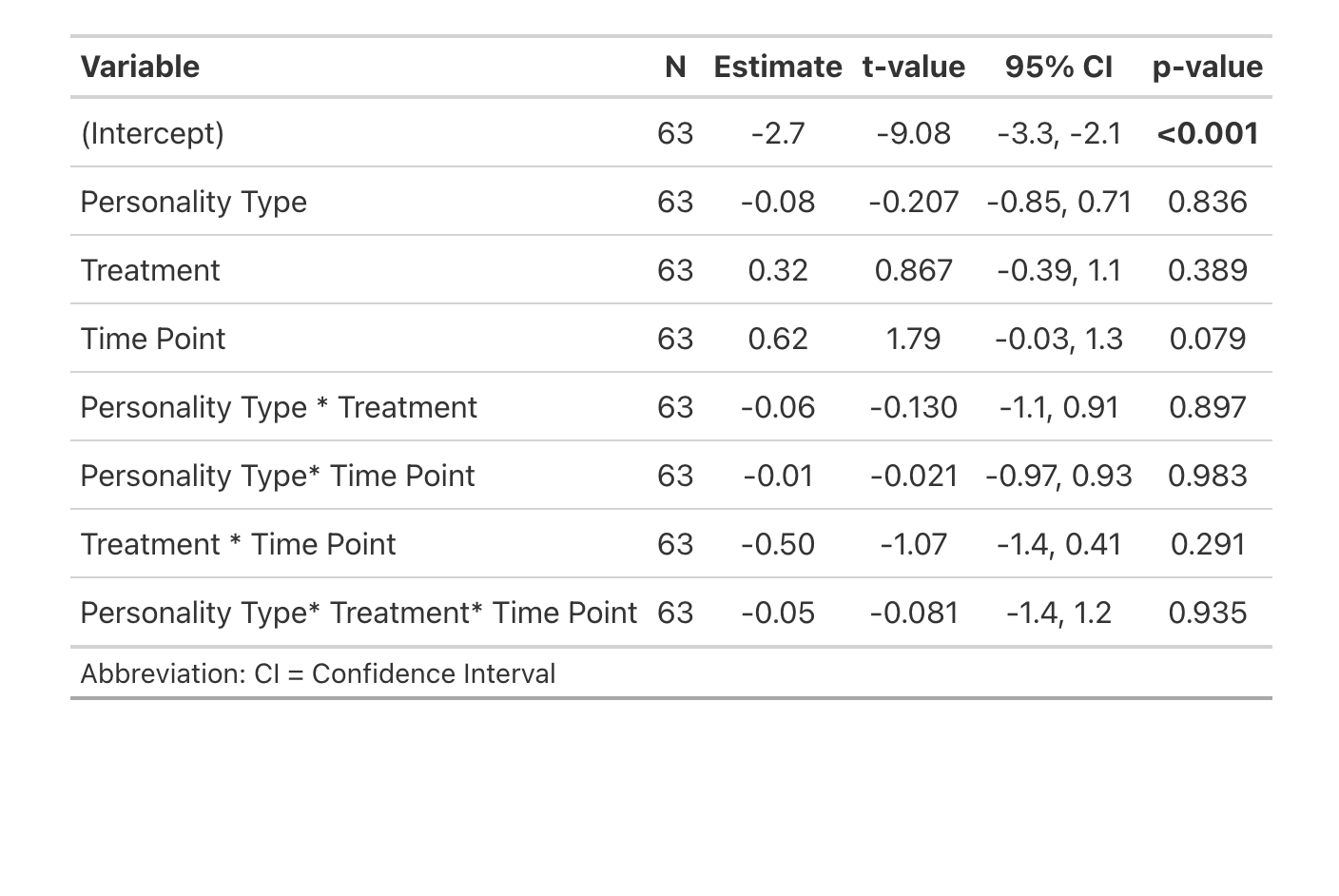

Q. Active dopaminergic Vs neurons (ps6 + TH double labelled)
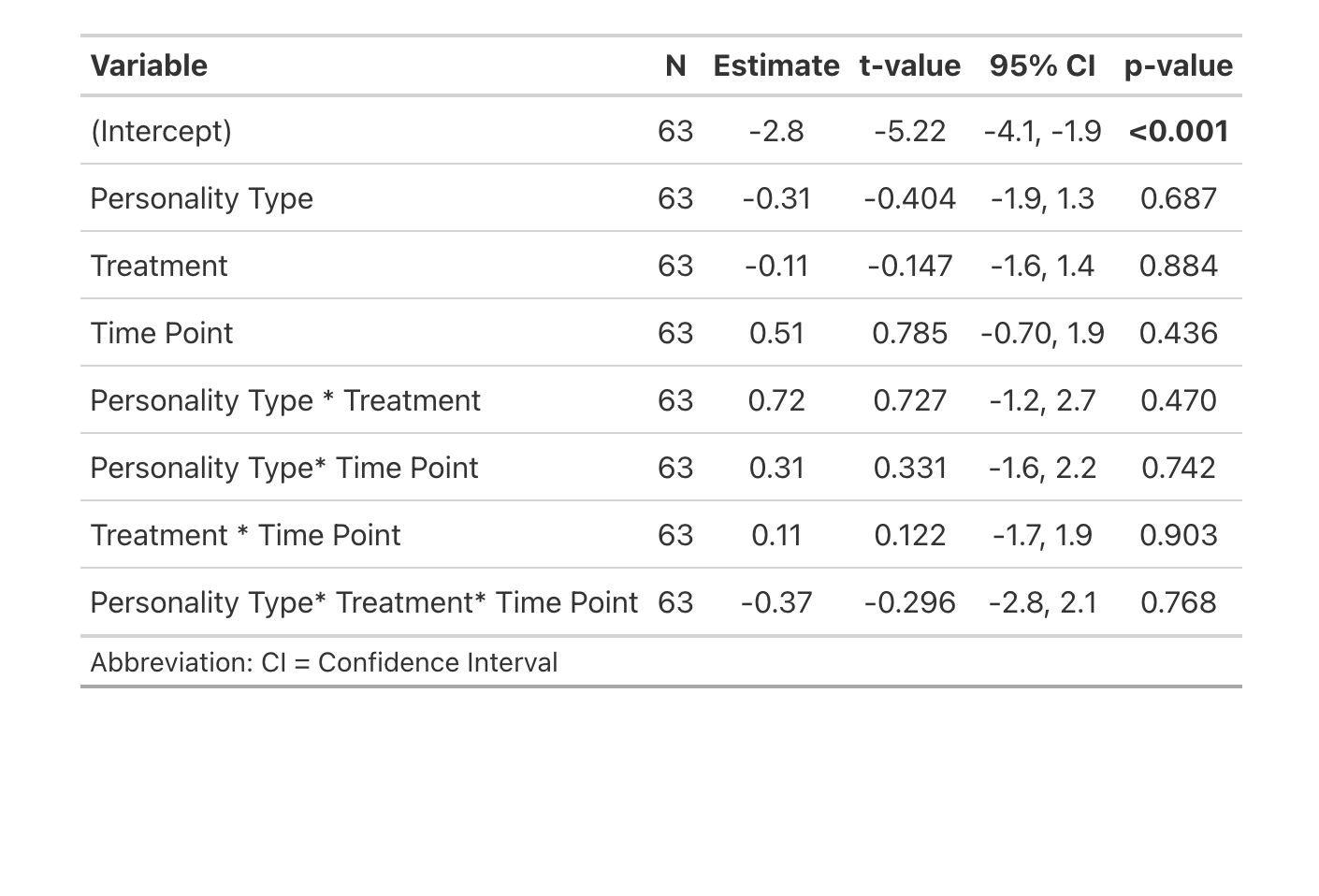

R. Active Vv neurons (ps6 labelled)
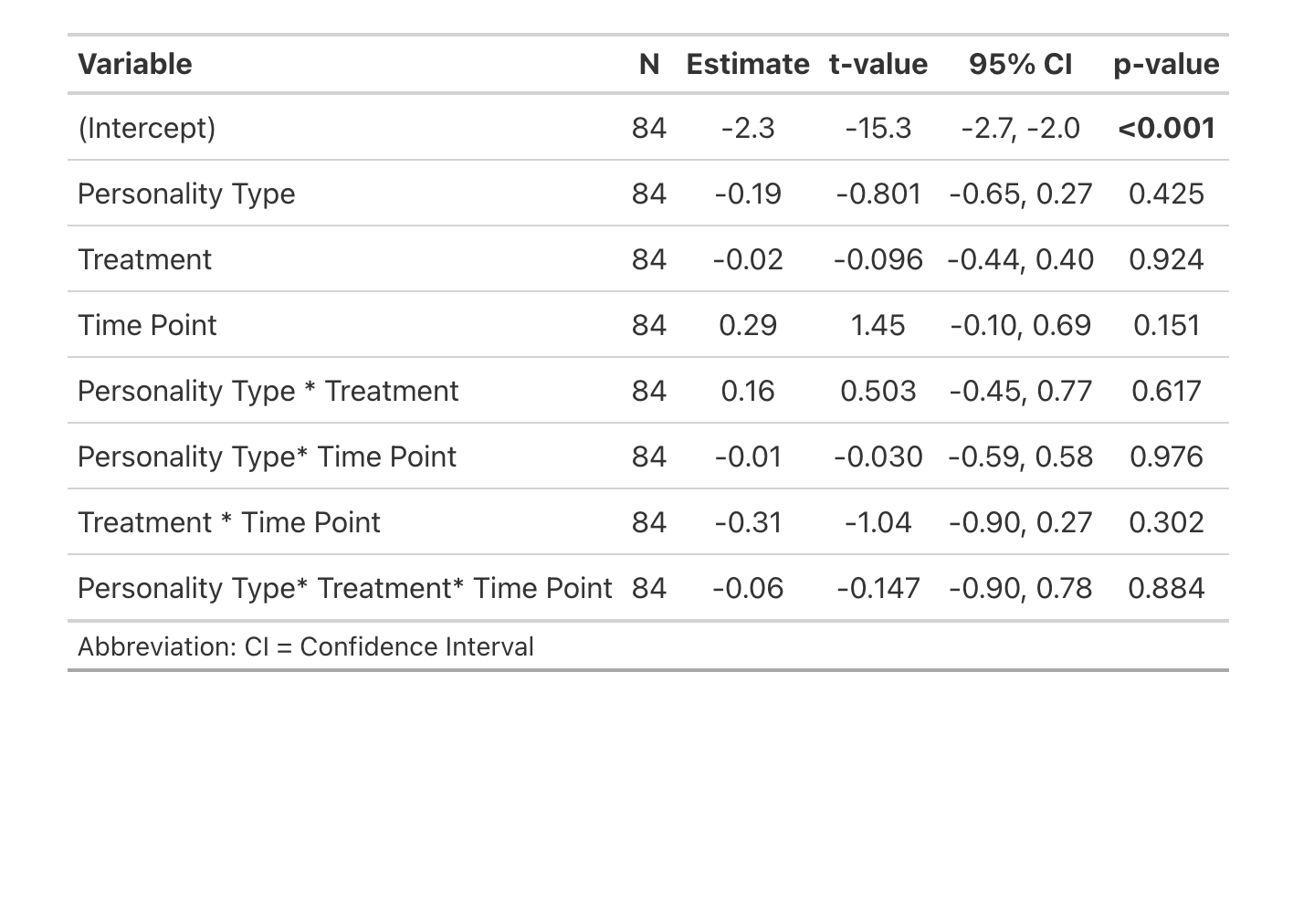

S. Active dopaminergic Vv neurons (ps6 + TH double labelled)
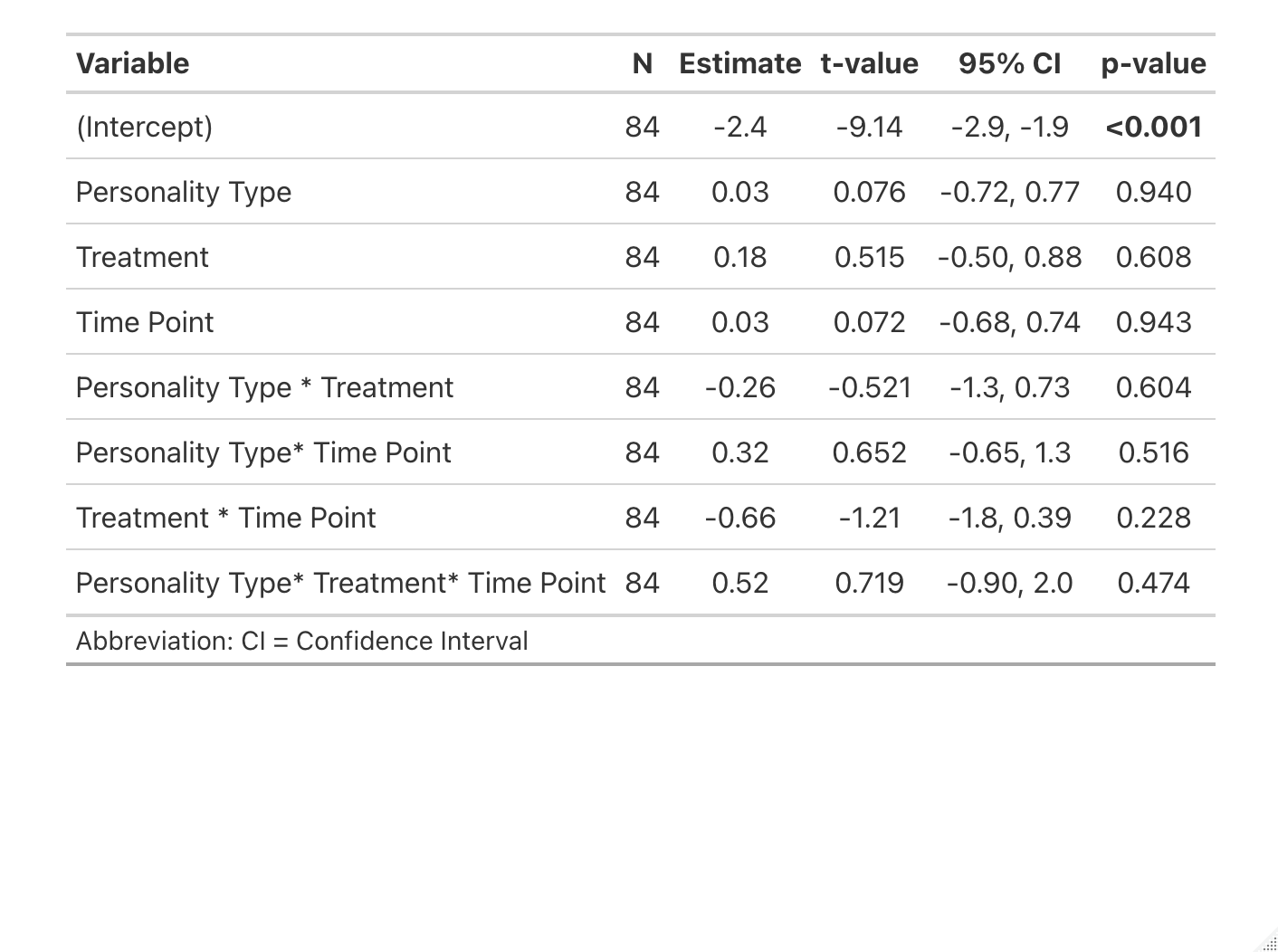

**Table 3. Models of the effects of change in behavior, treatment, personality type, and interactions on the proportion of active cells, active dopamine, and active serotonin cells**

A. Active Dm neurons (ps6 labelled)

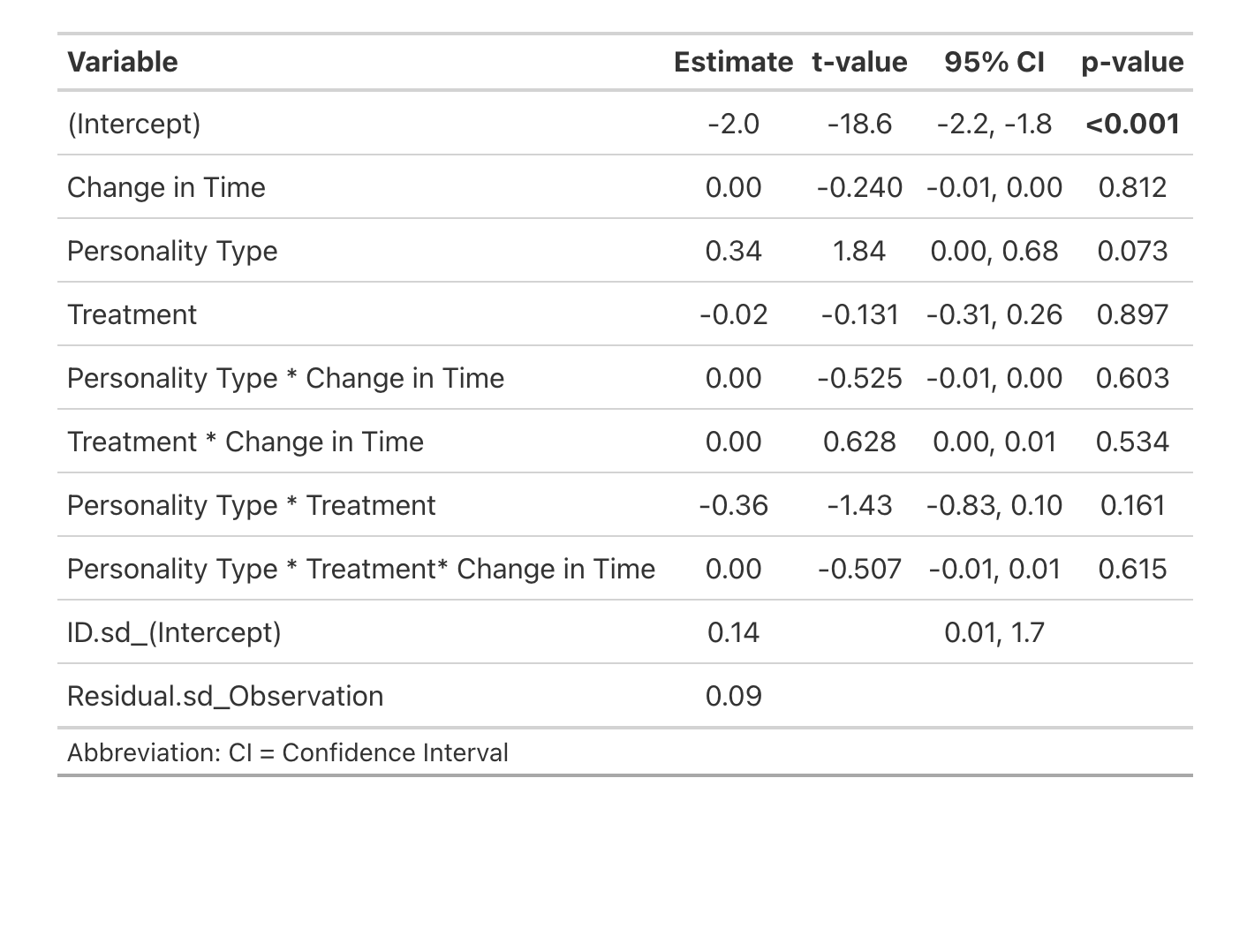

B. Active Dl neurons (ps6 labelled)
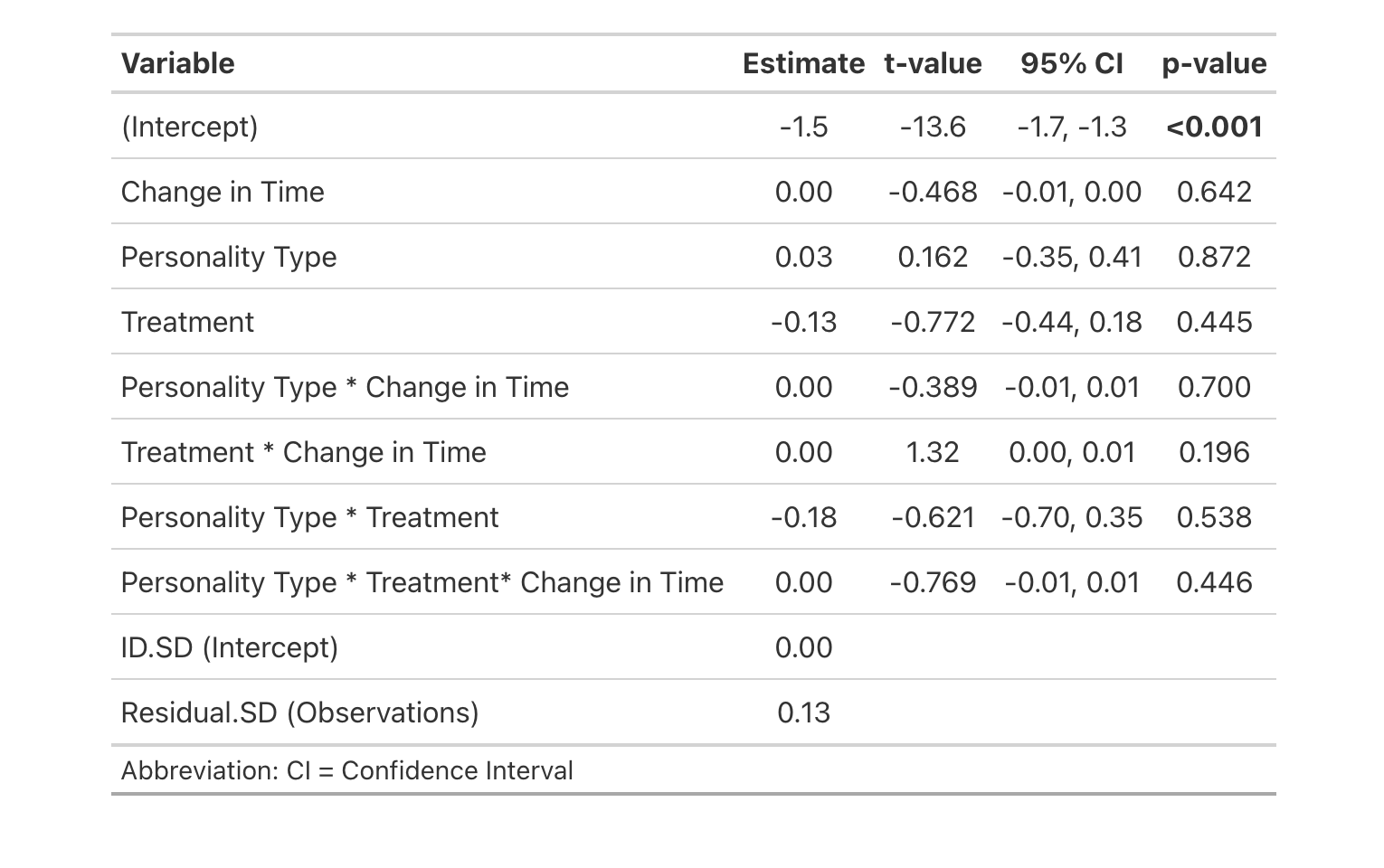

C. Active Dp neurons (ps6 labelled)
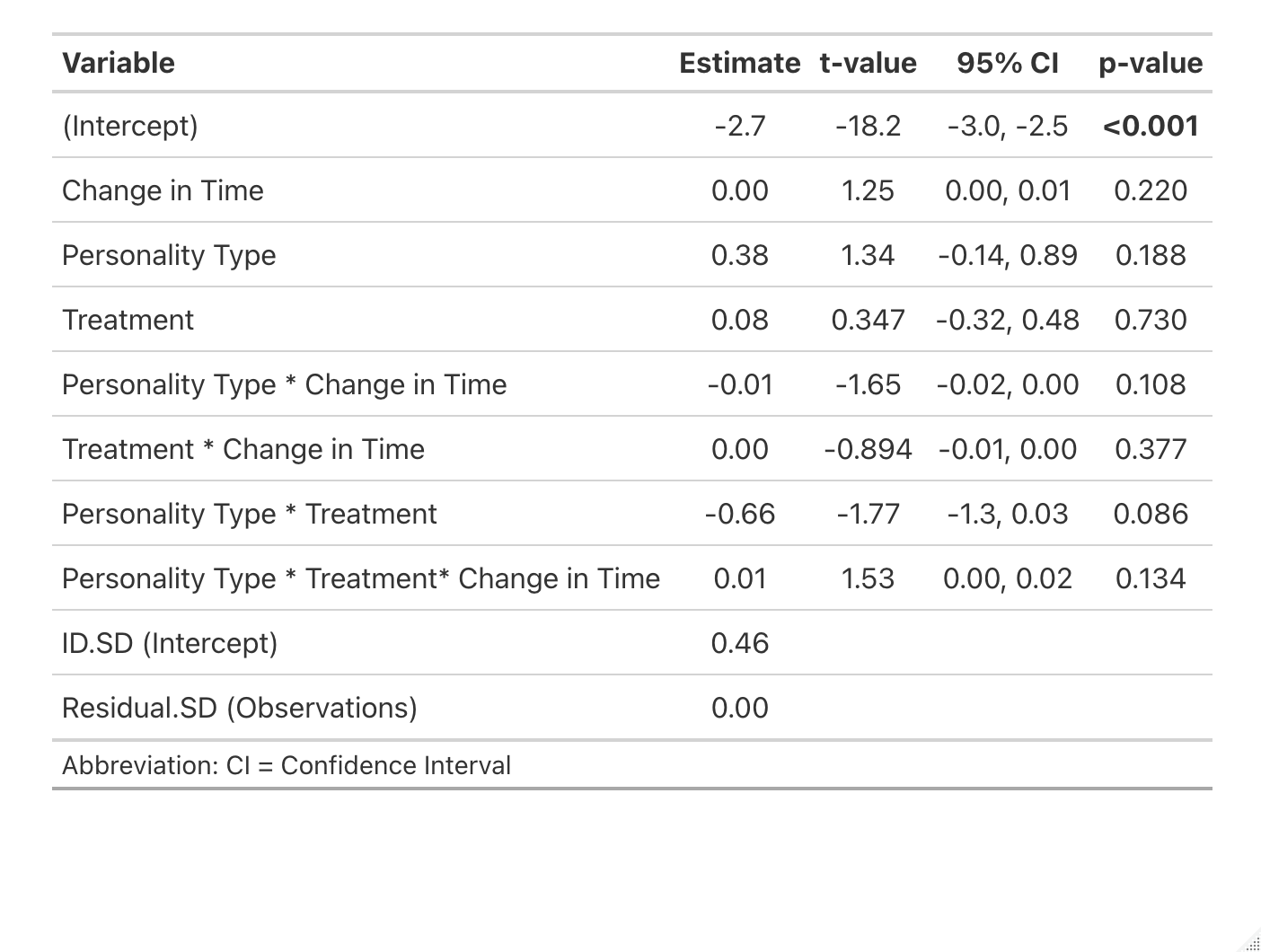

D. Active Hb neurons (ps6 labelled)
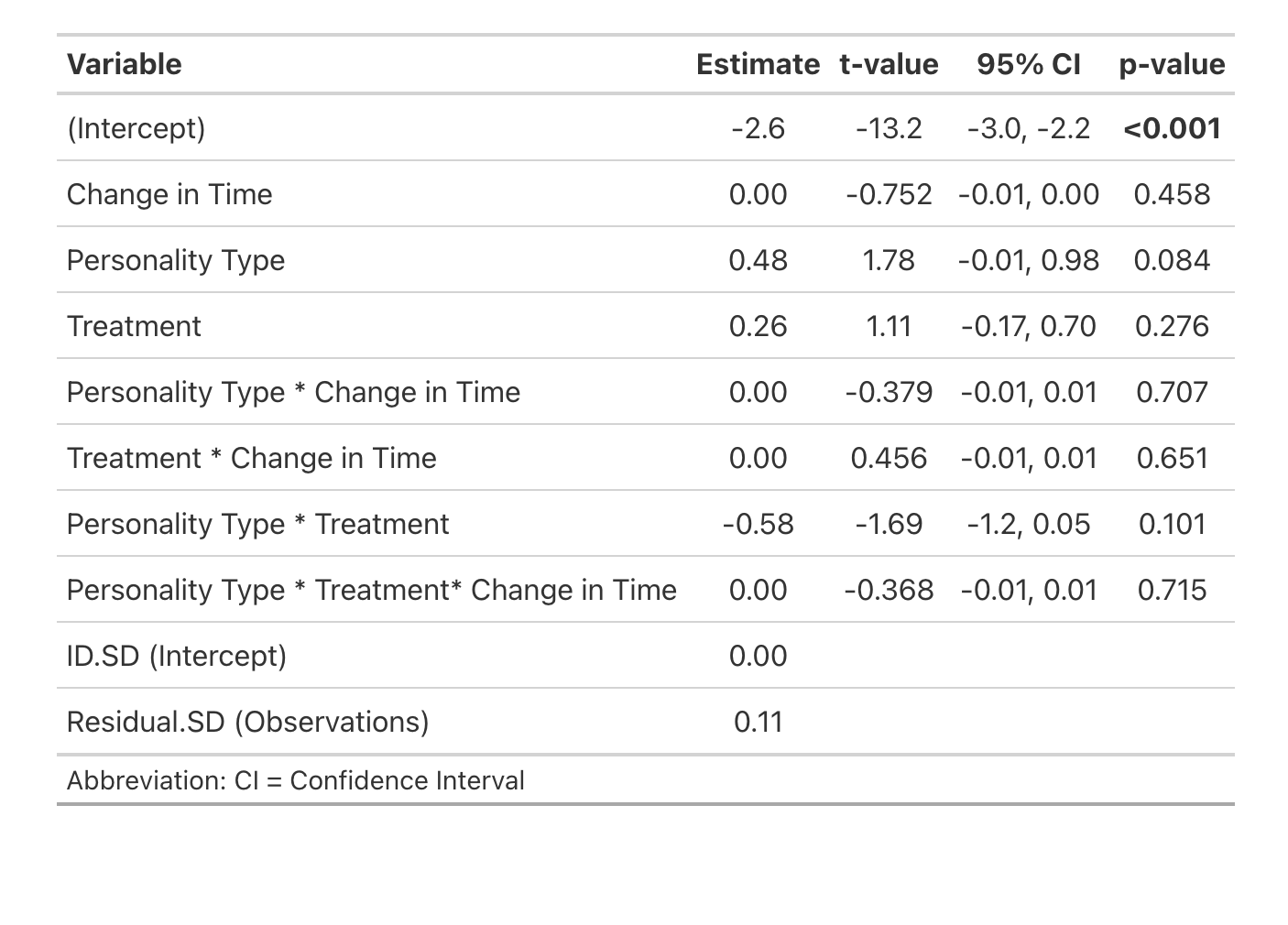

E. Active Hd neurons (ps6 labelled)
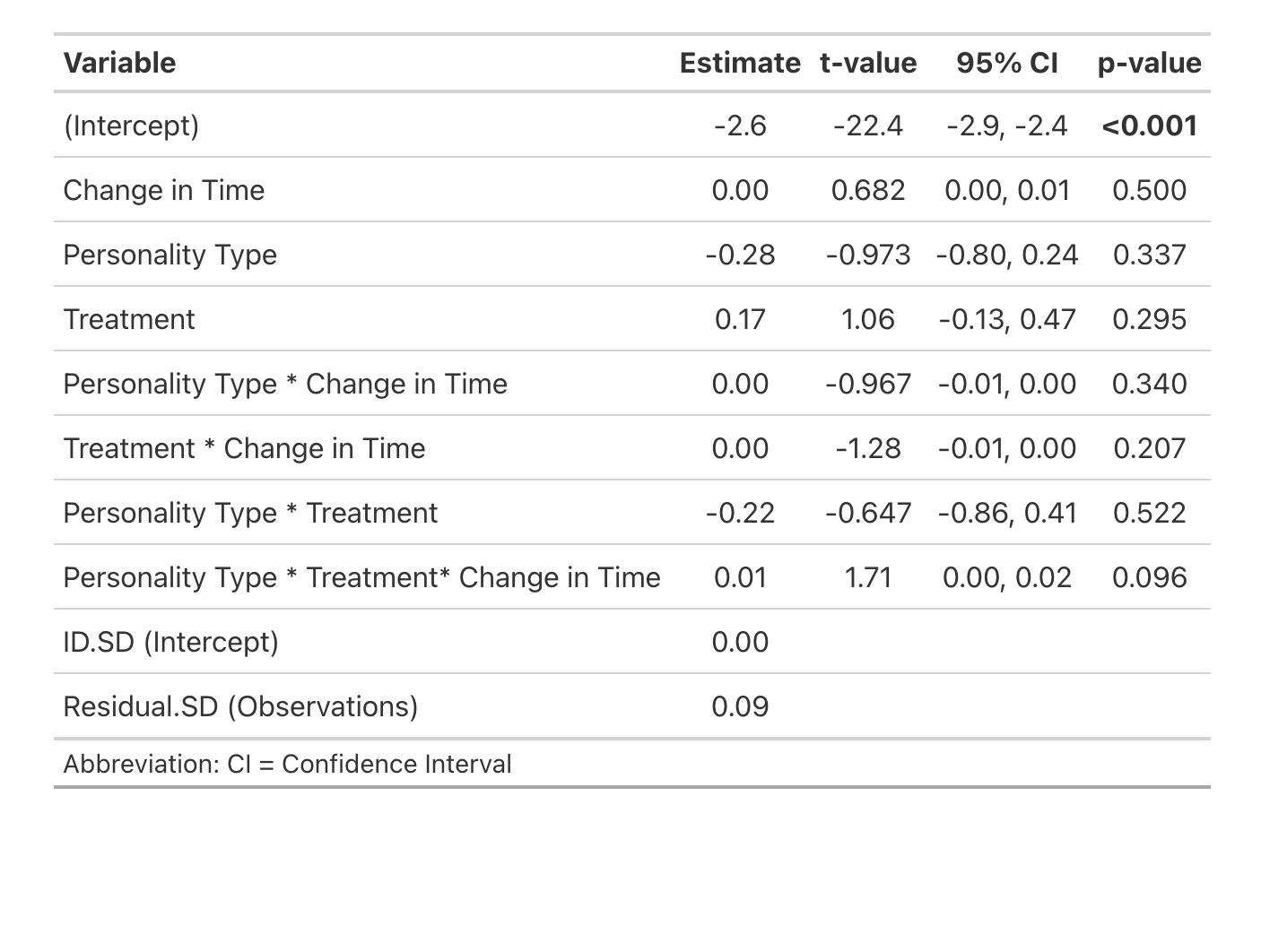

F. Active serotonergic Hd neurons (ps6 + 5HT labelled)
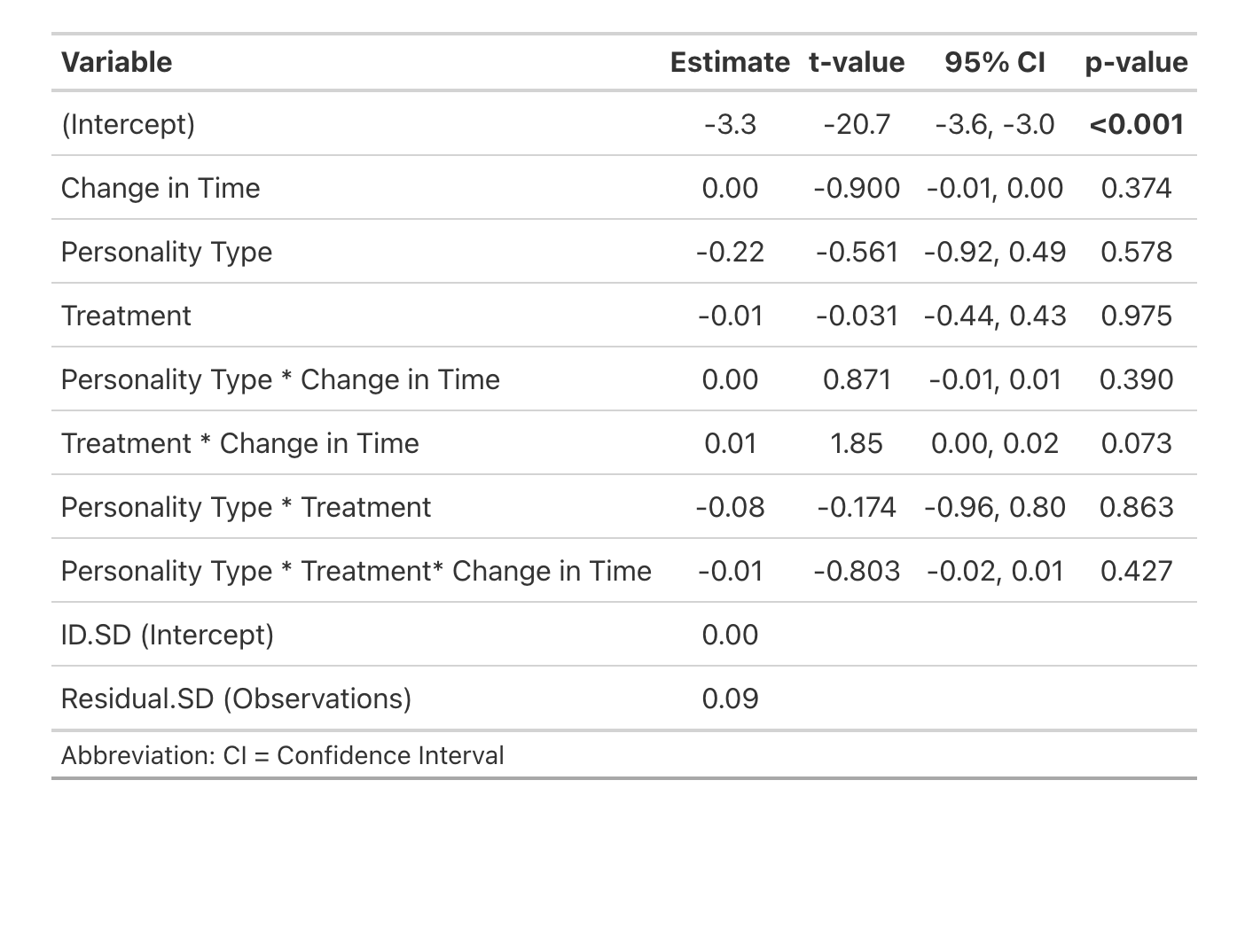

G. Active Hv neurons (ps6 labelled)
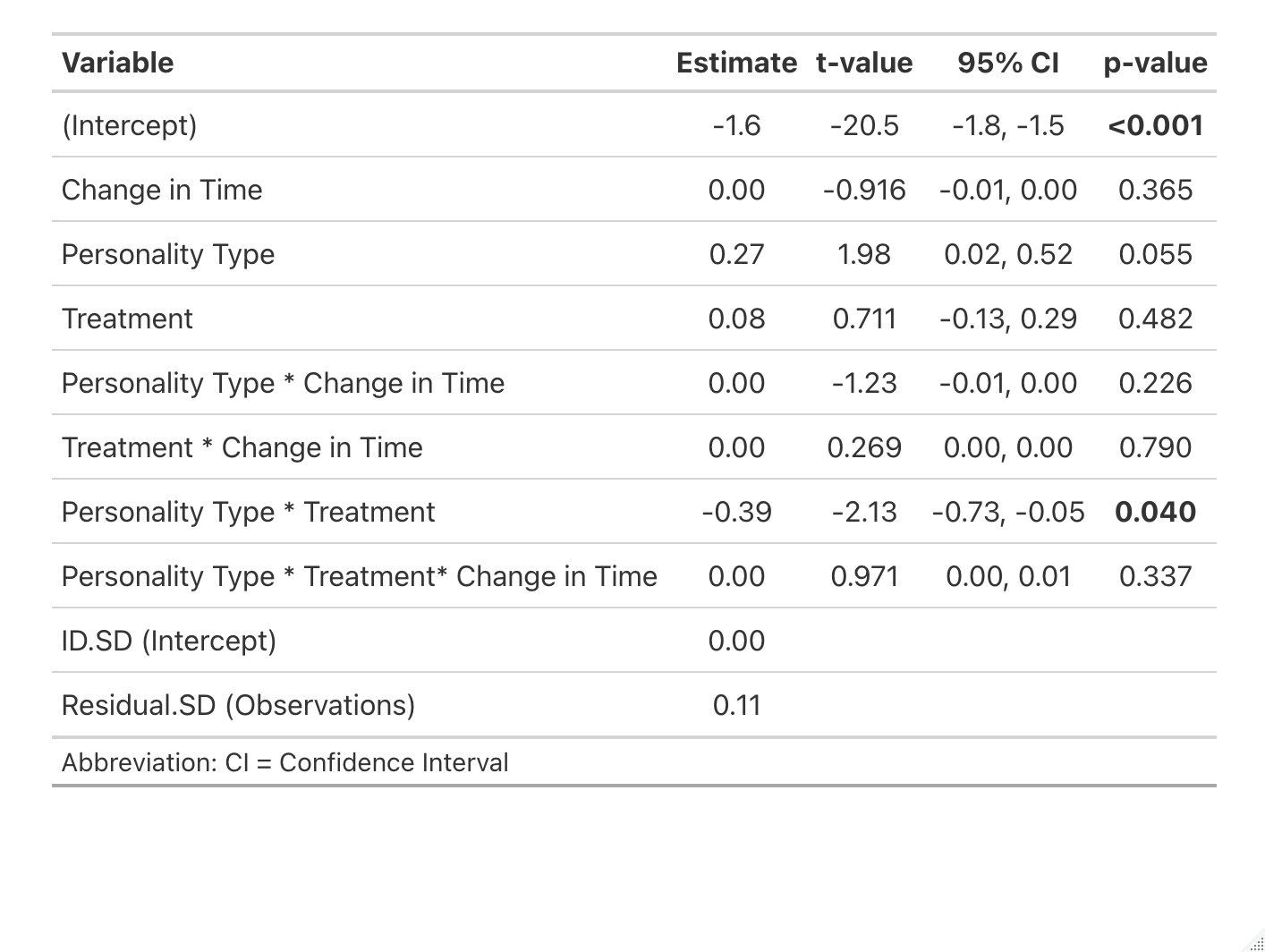

H. Active POA neurons (ps6 labelled)
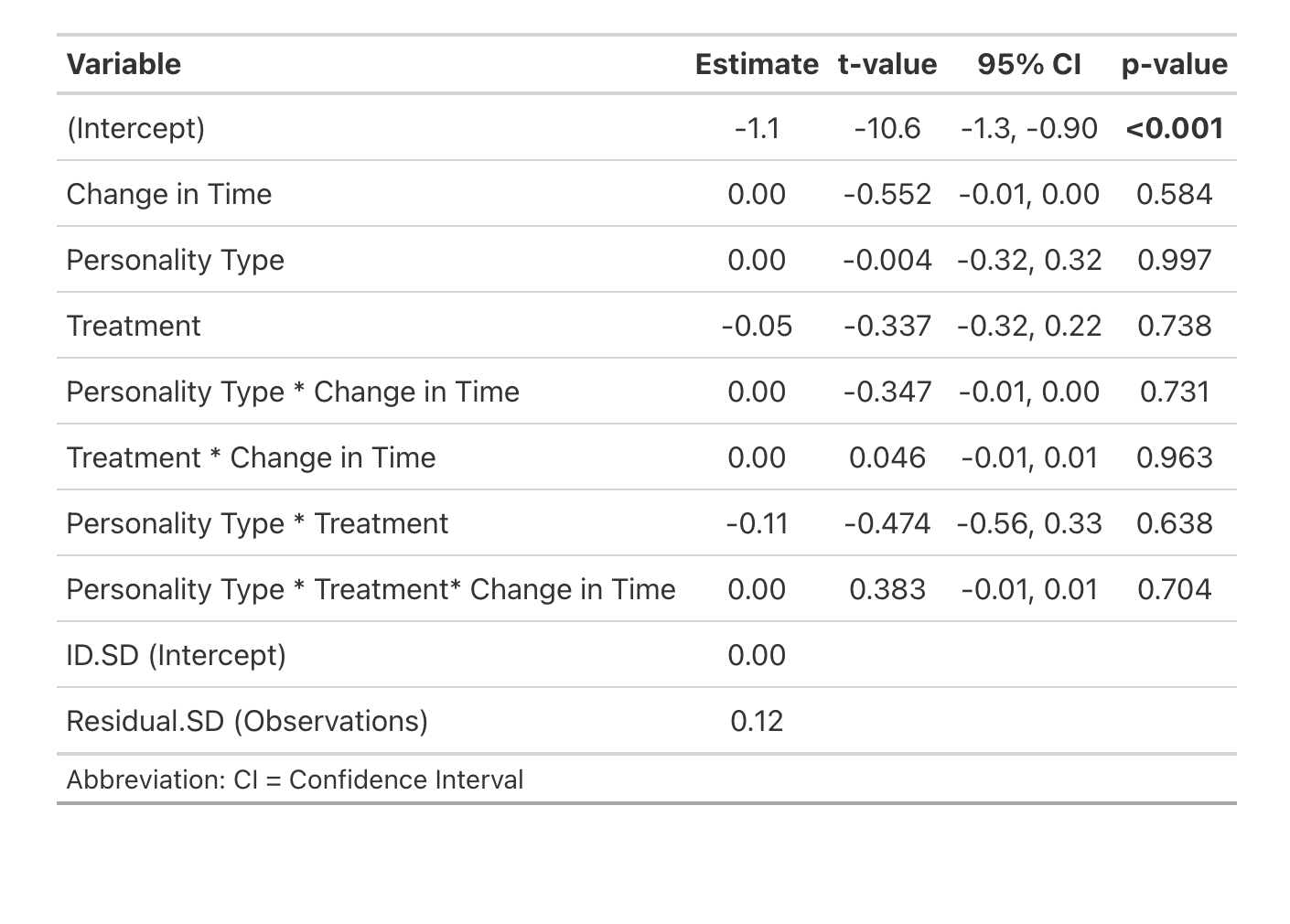

I. Active dopaminergic POA neurons (ps6 + TH double labelled)
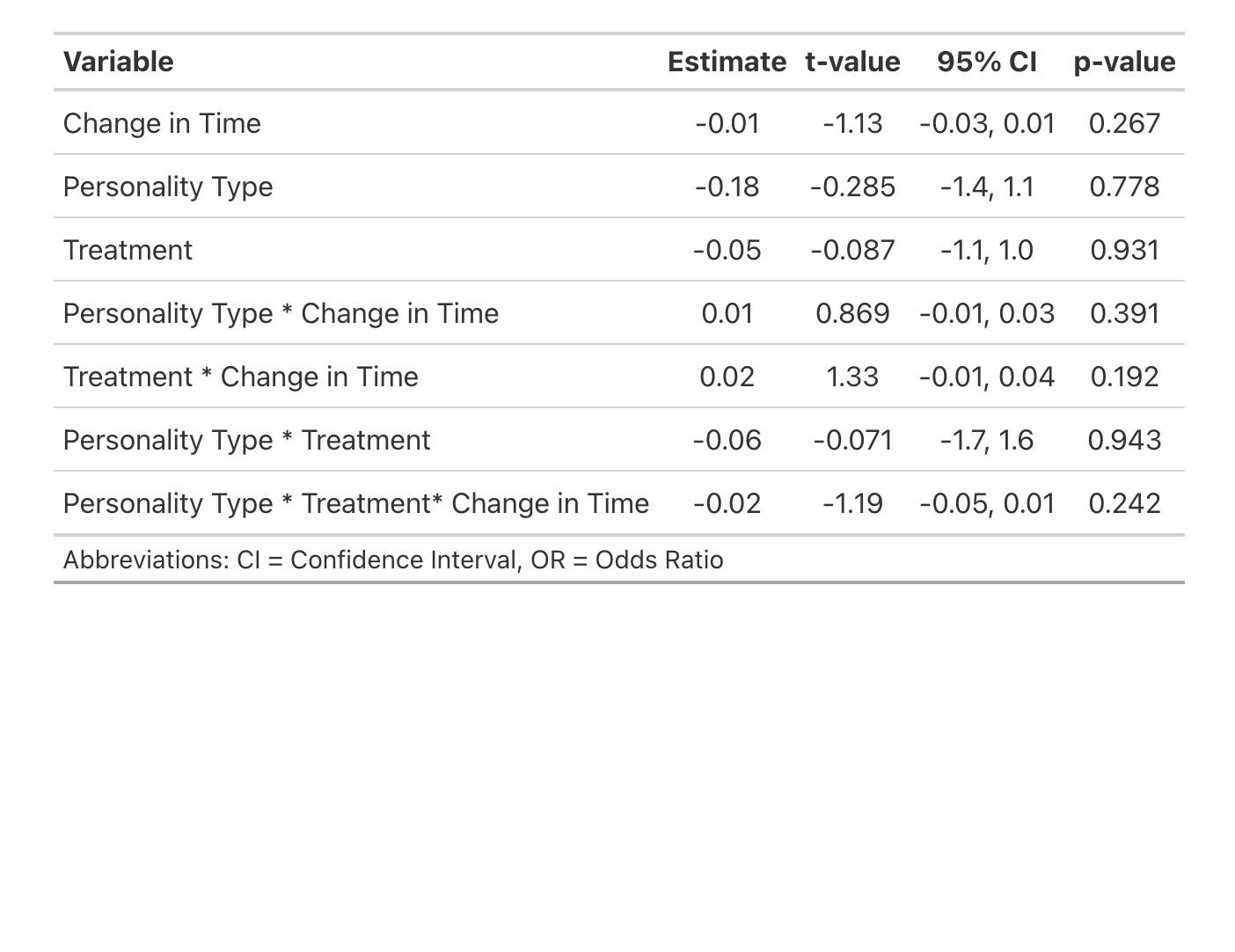

J. Active PT neurons (ps6 labelled)
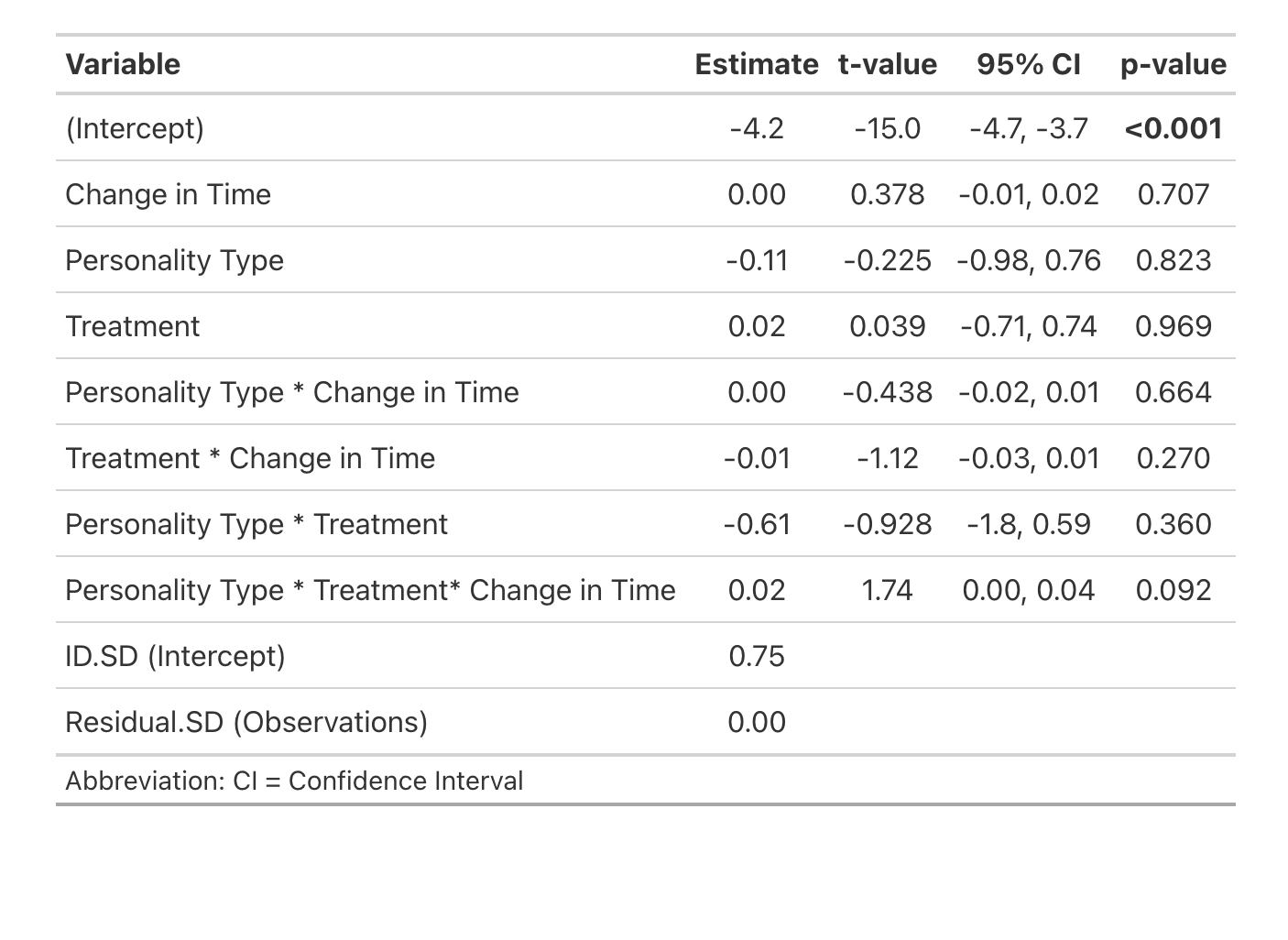

K. Active dopaminergic PT neurons (ps6 + TH double labelled)
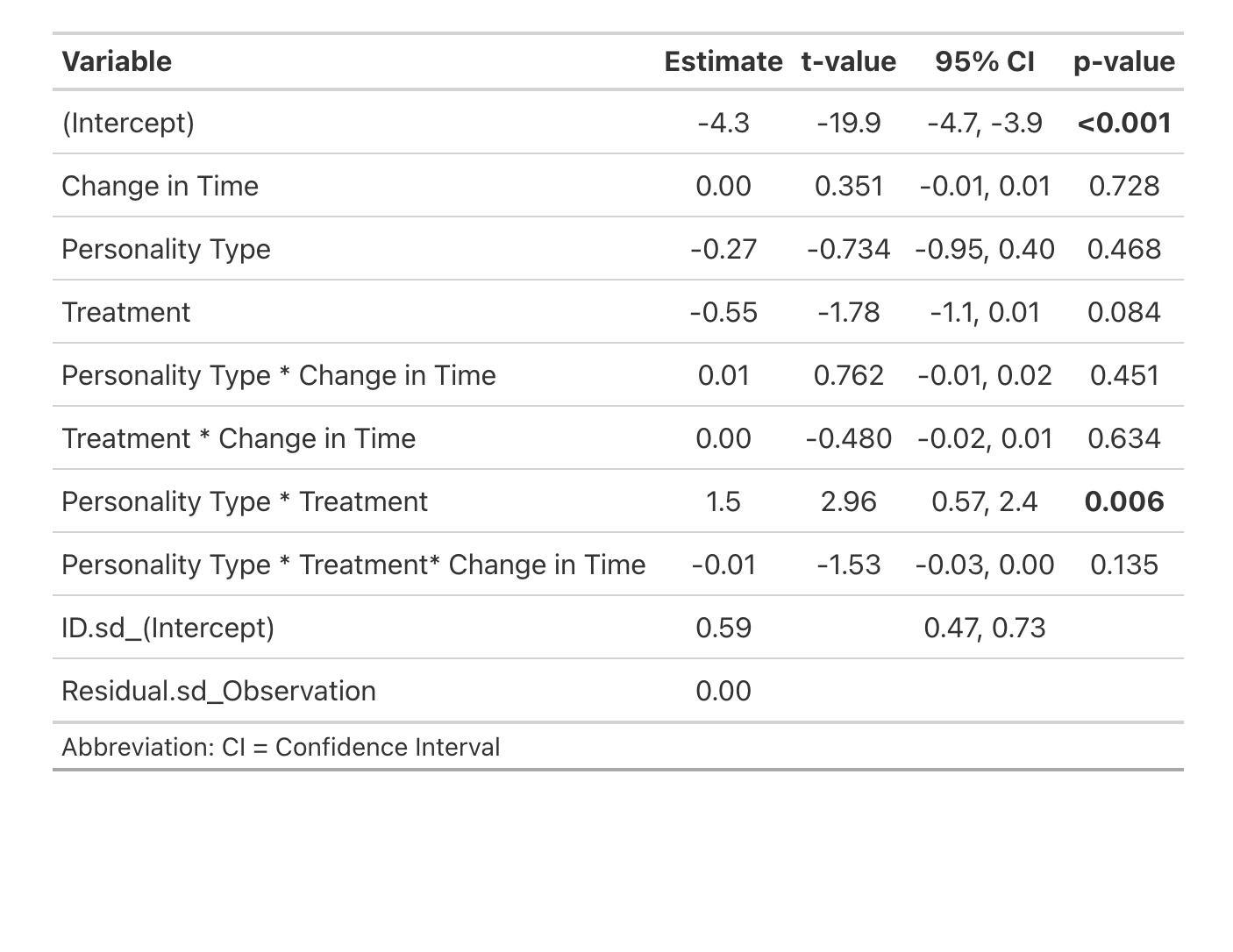

L. Active SR neurons (ps6 labelled)
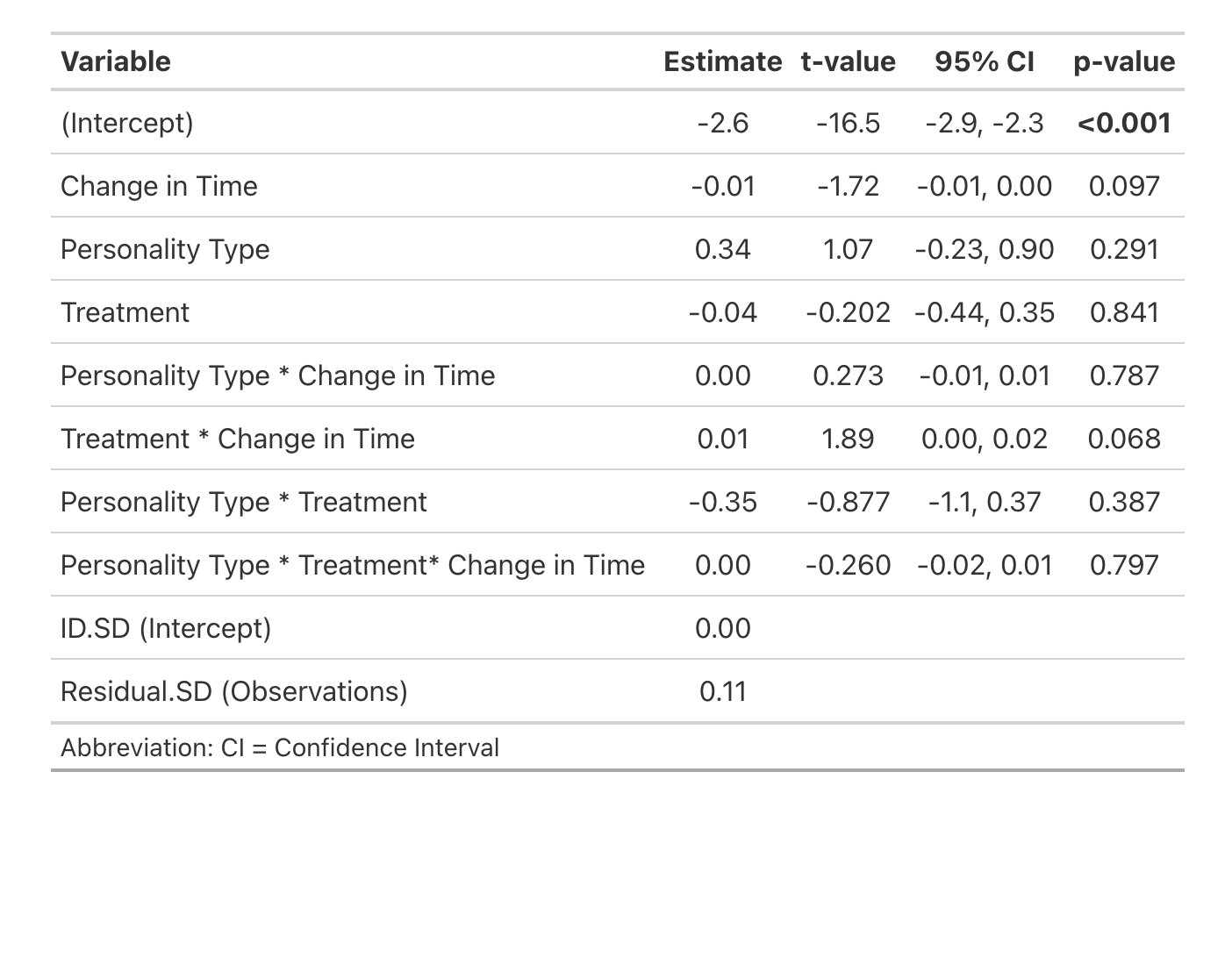

M. Active serotonergic SR neurons (ps6 + 5HT double labelled)
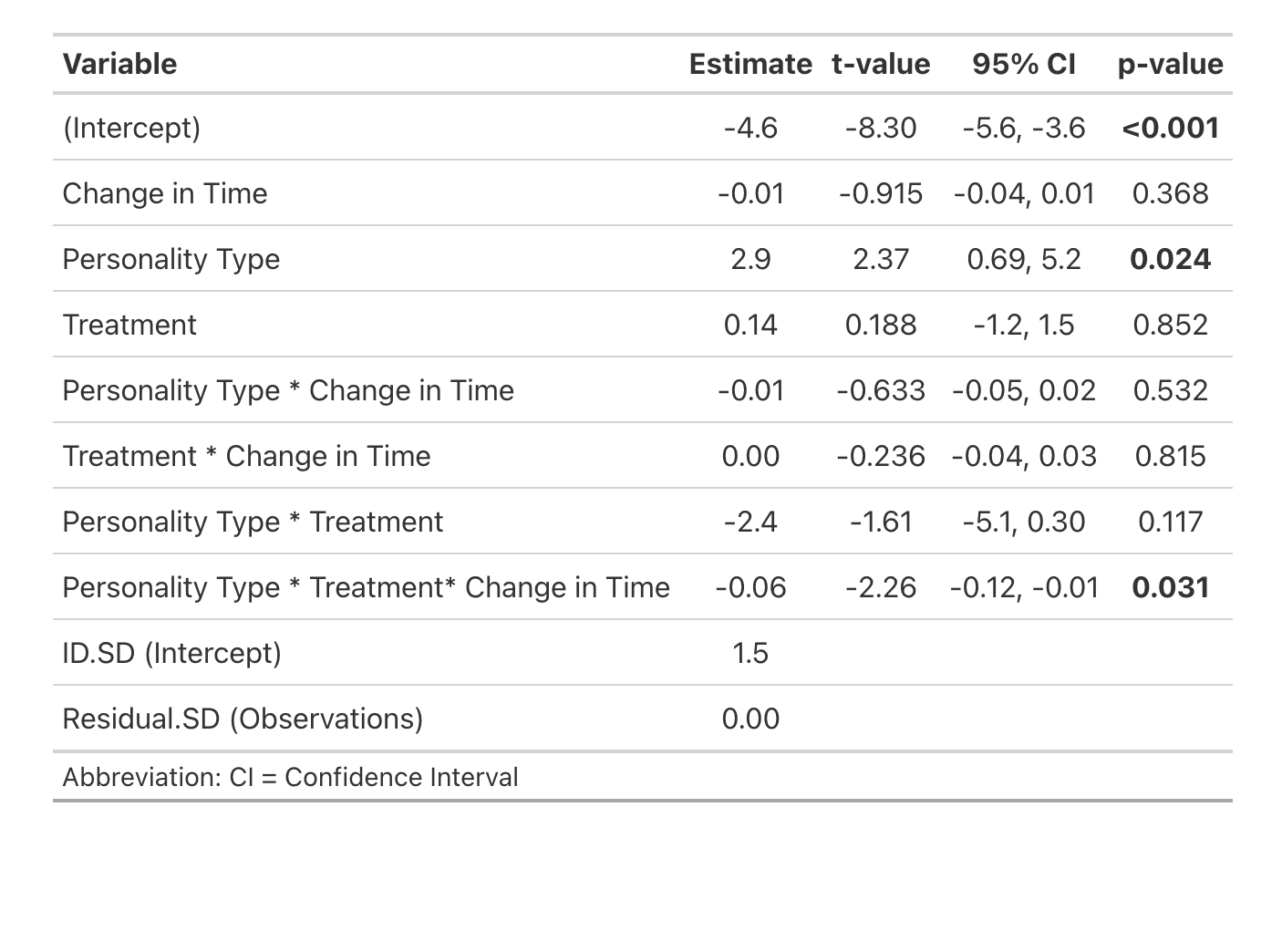

N. Active Tpp neurons (ps6 labelled)
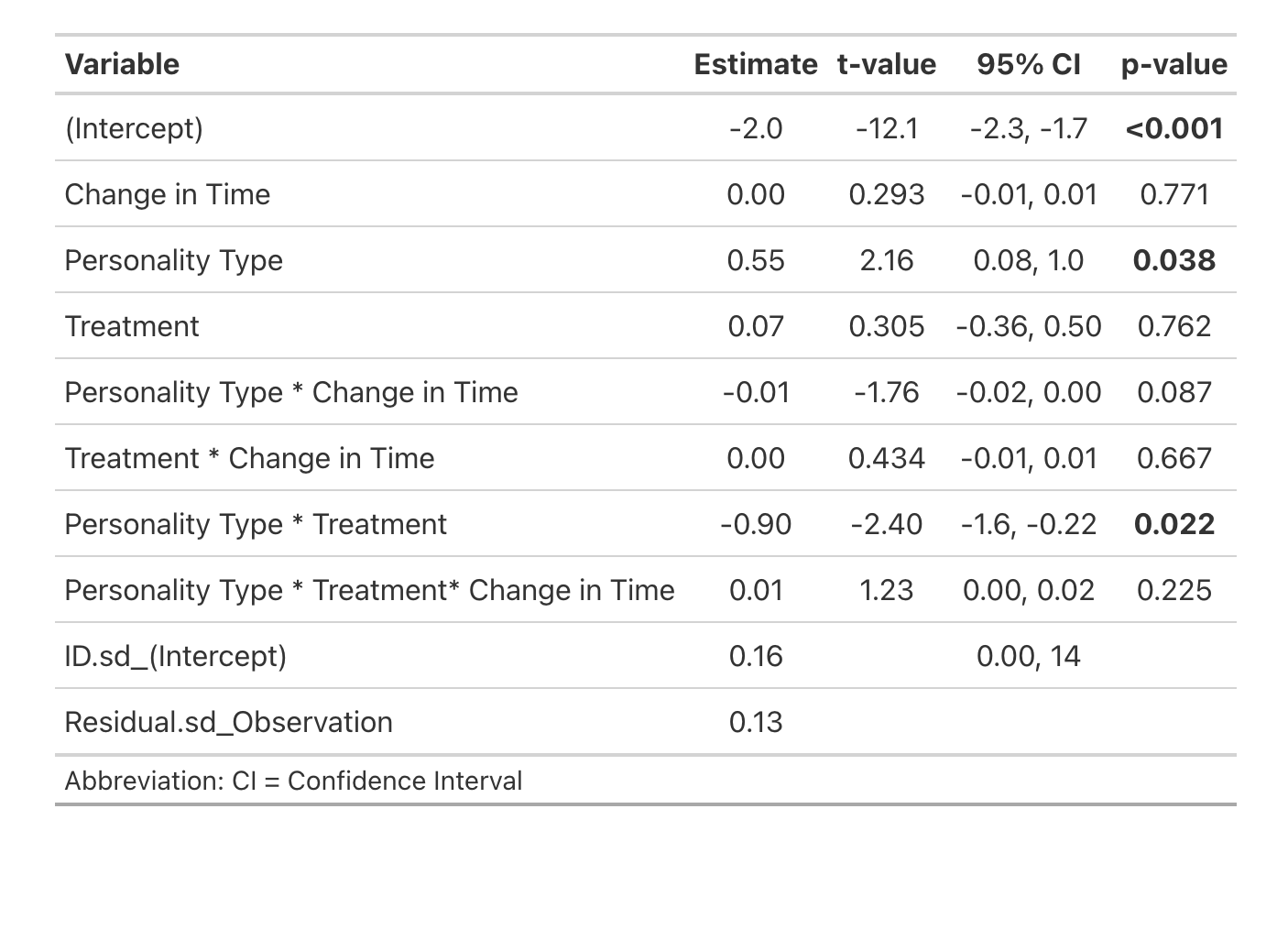

O. Active dopaminergic Tpp neurons (ps6 + TH double labelled)
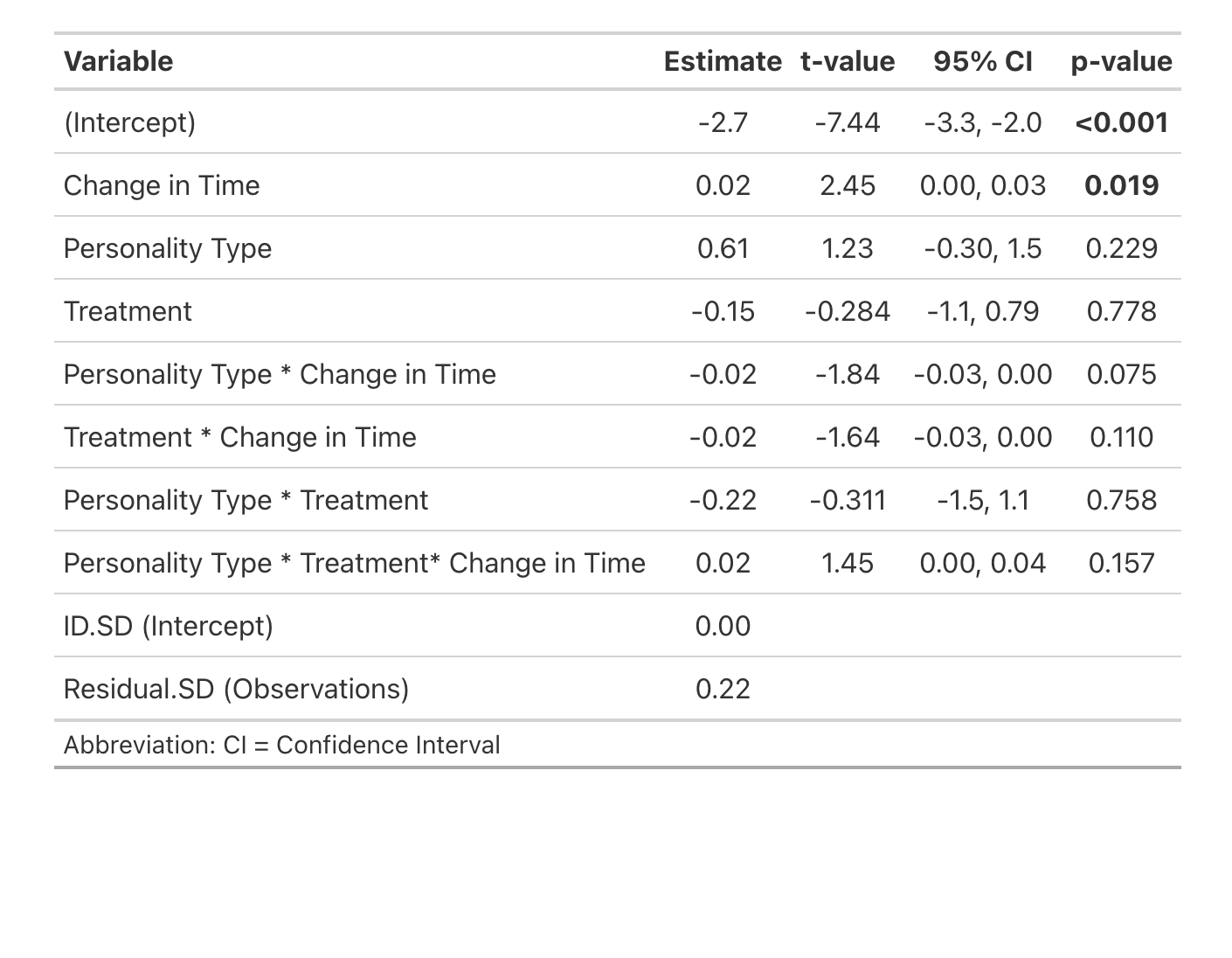

P. Active Vs neurons (ps6 labelled)
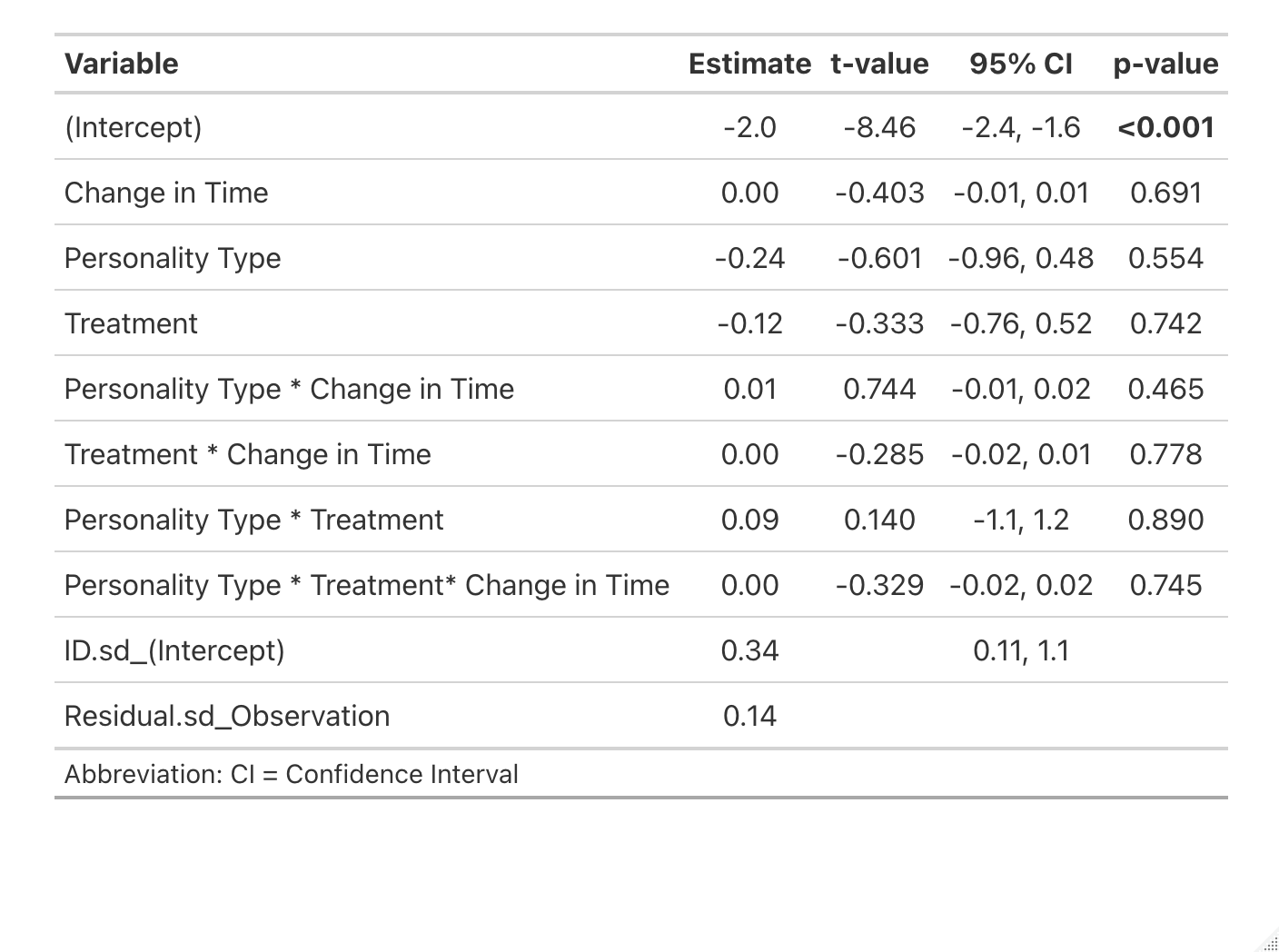

Q. Active dopaminergic Vs neurons (ps6 + TH labelled)
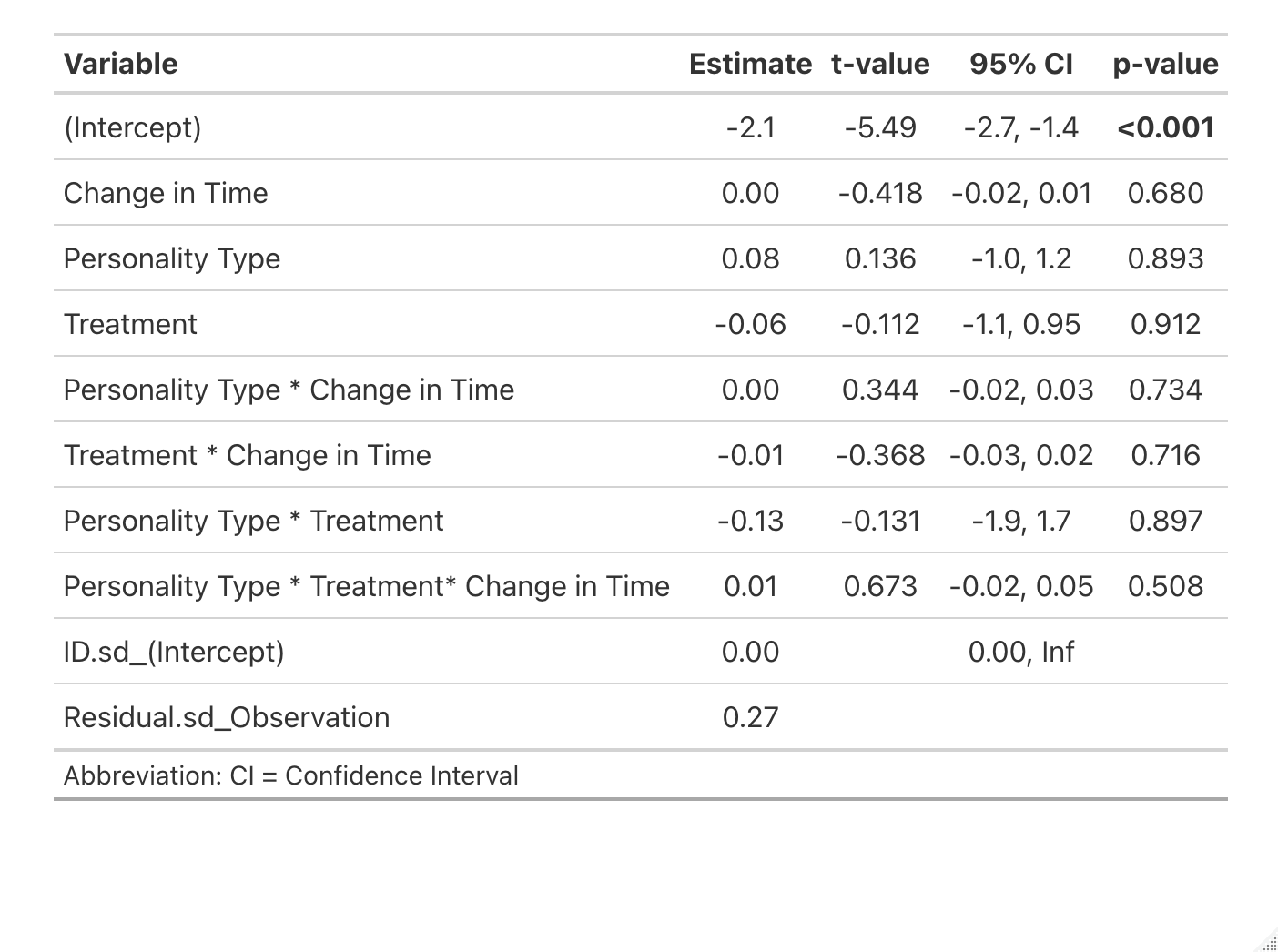

R. Active Vv neurons (ps6 labelled)
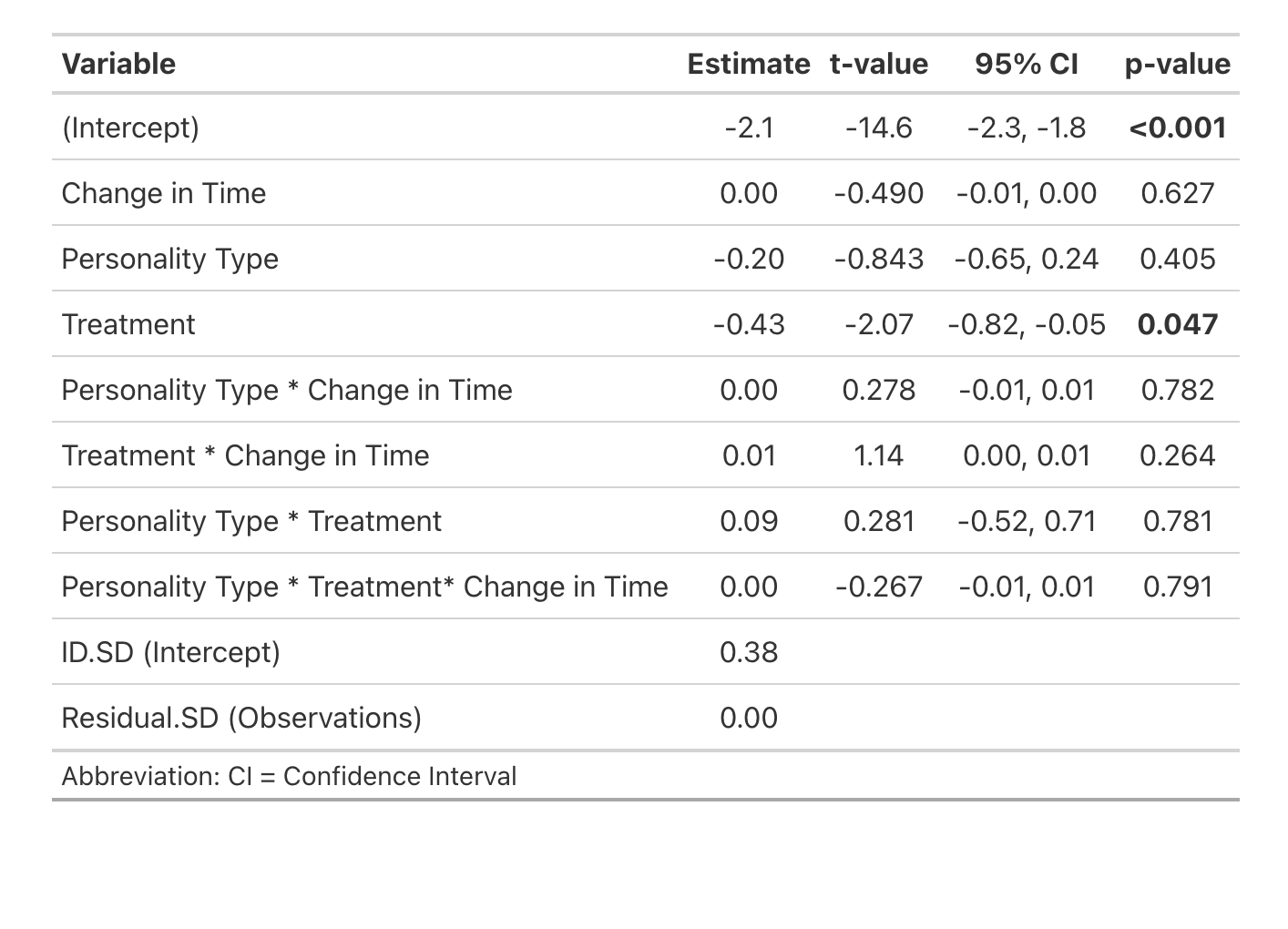

S. Active dopaminergic Vv neurons (ps6 + TH double labelled)
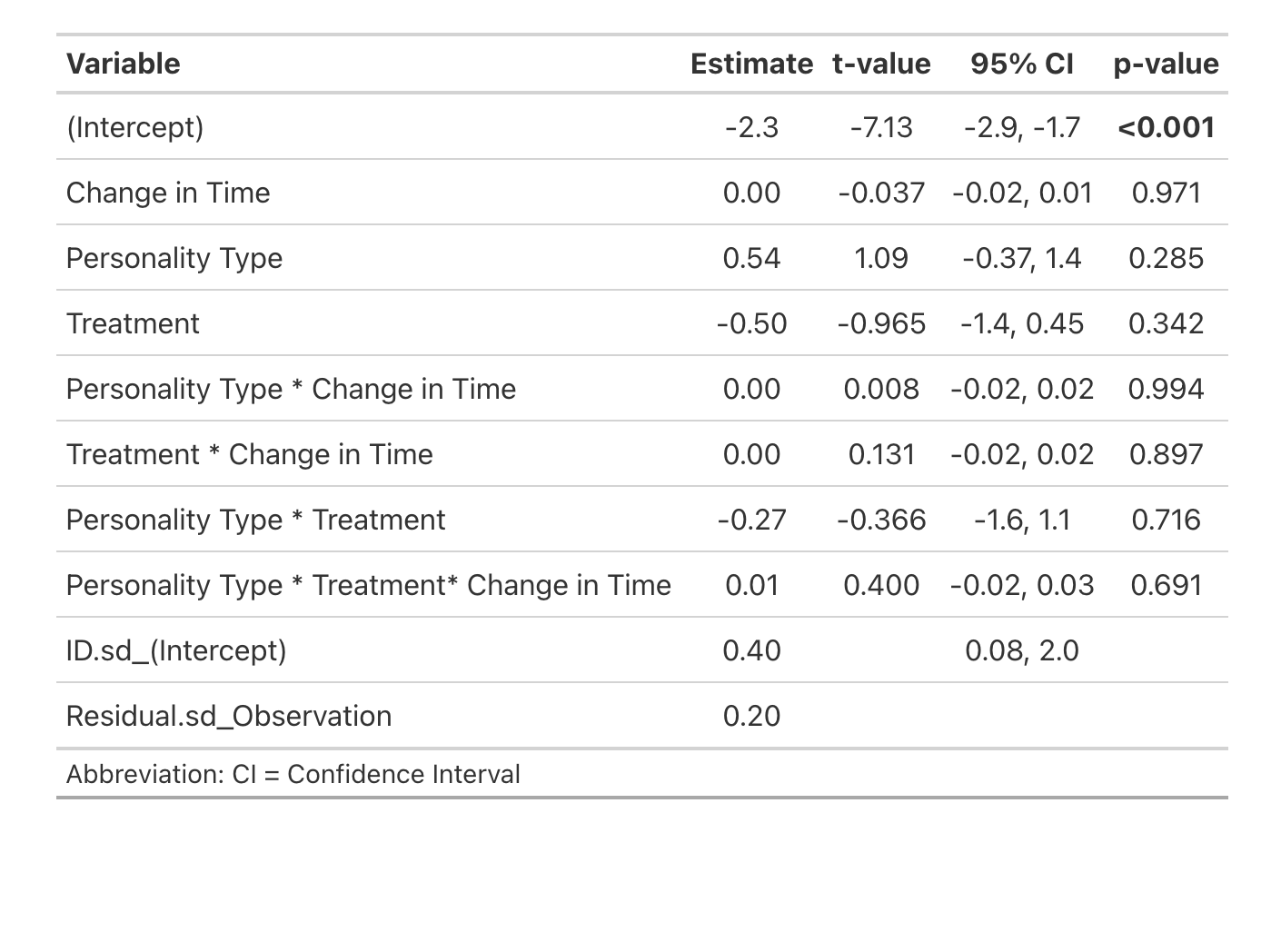

**Table 4. Model of the effects of conditioning day, treatment, personality type and interactions on the time spent in the conditioned zone in the conditioned place preference task**

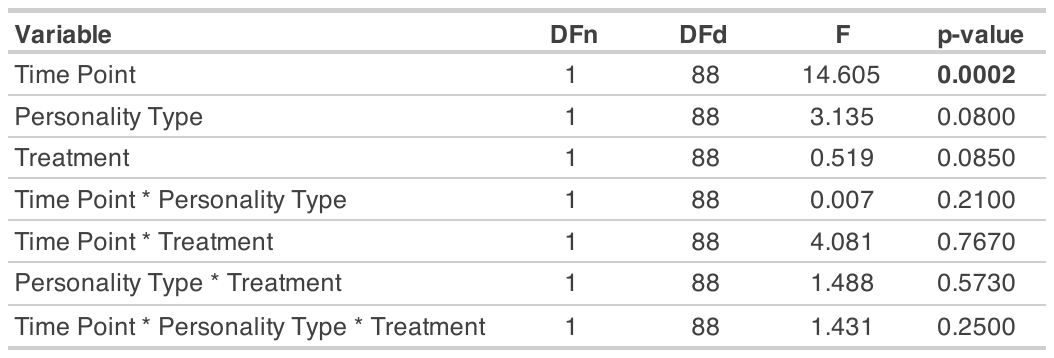

**Table 5. Model of the effects of conditioning day, treatment, personality type, and interactions on the time spent in the conditioned zone in the conditioned place preference task from Corcoran et al., (2025).**

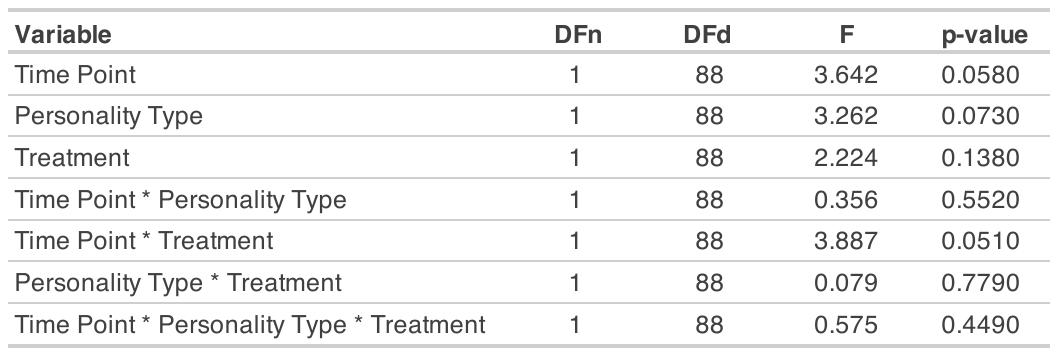

**Table 6. Model of the effects of time point, personality type, and treatment on the distance moved in conditioned place preference task**

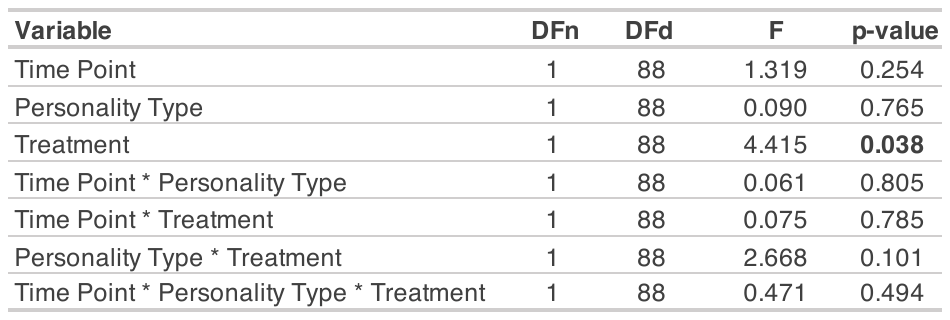

**Table 7. Raw Behavior Data**

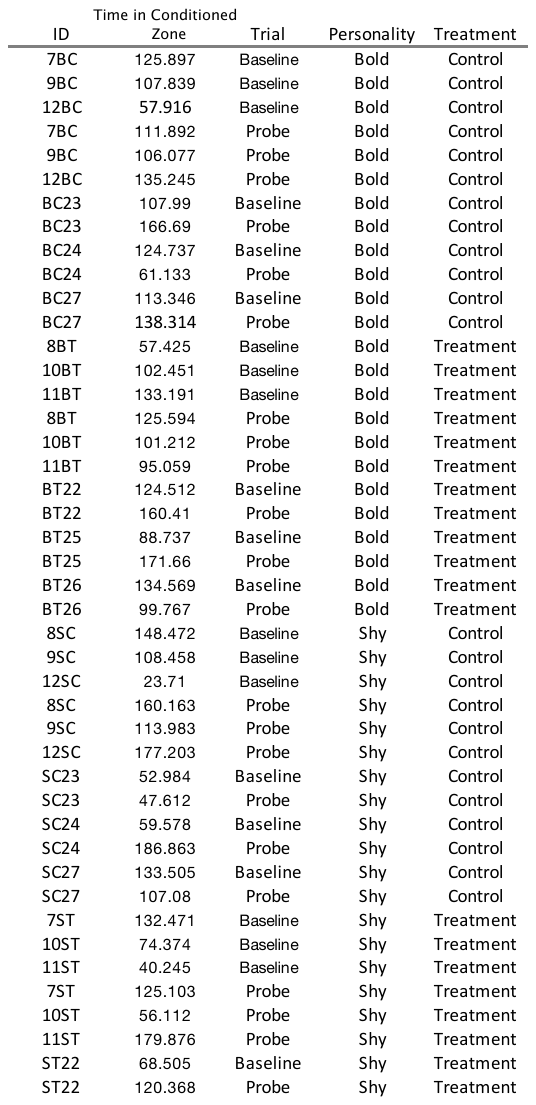

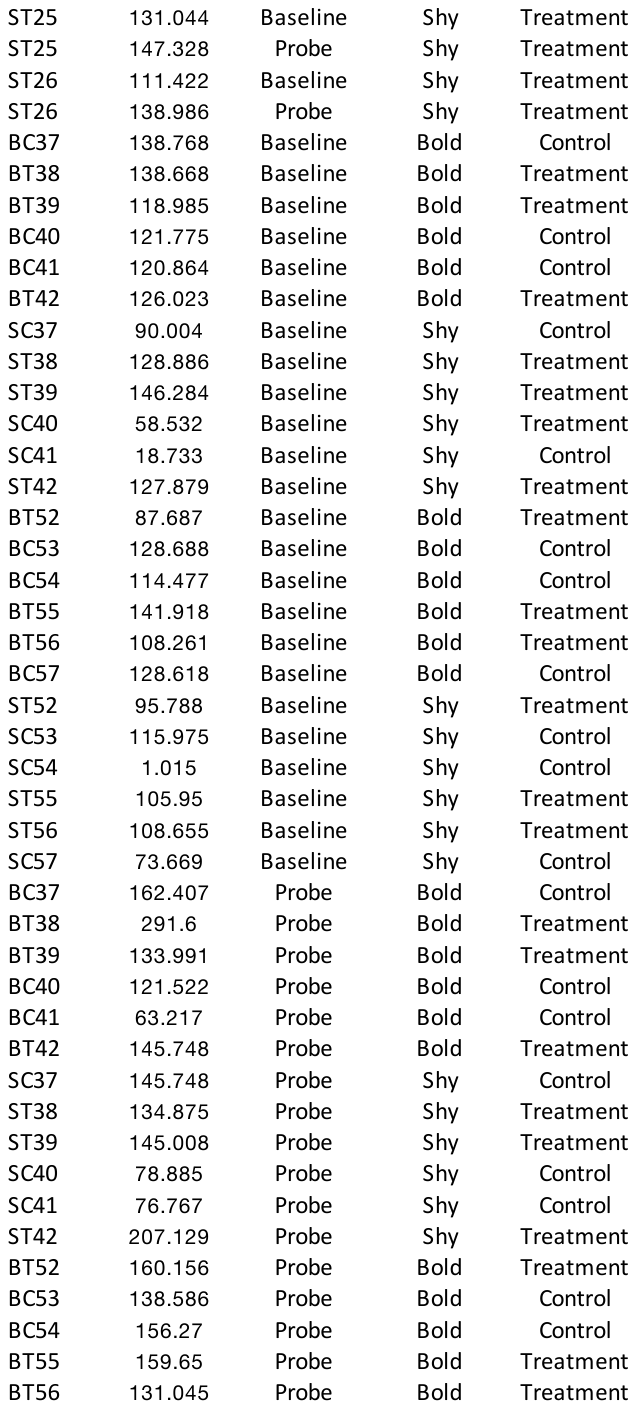

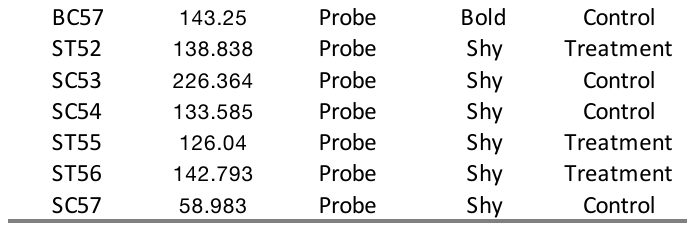

**Table 8. Raw Cell Count Data**

A. Dm
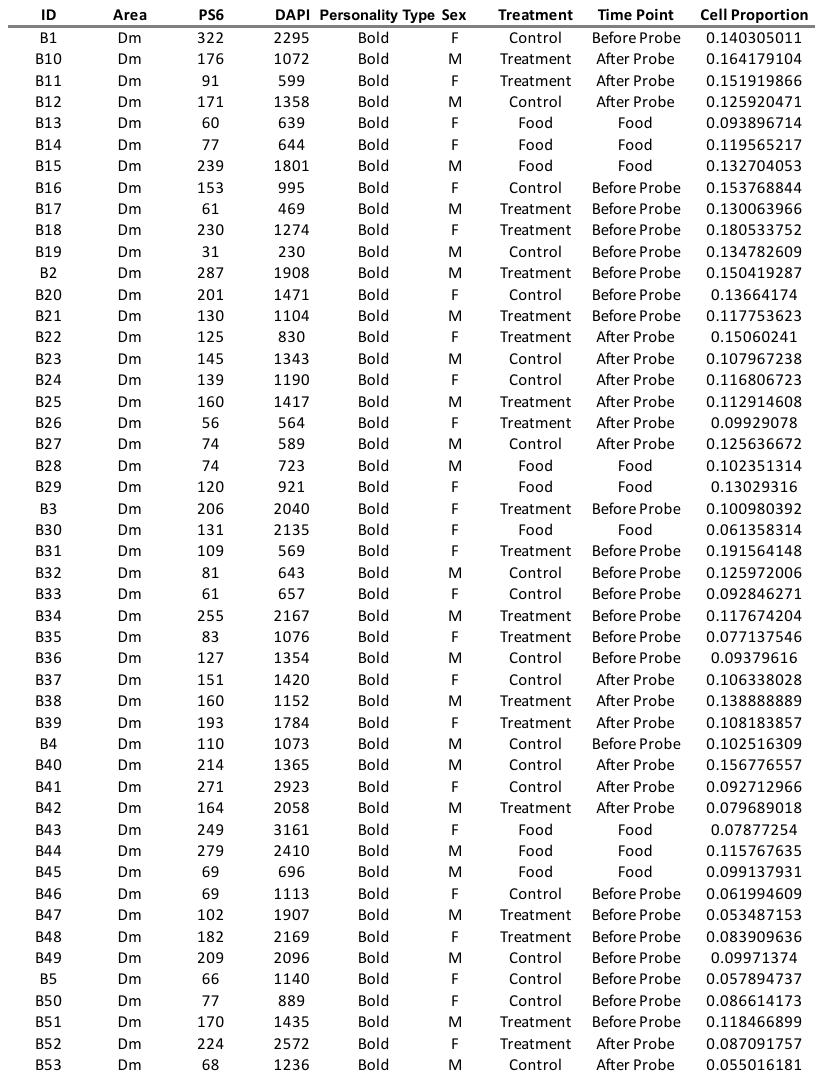

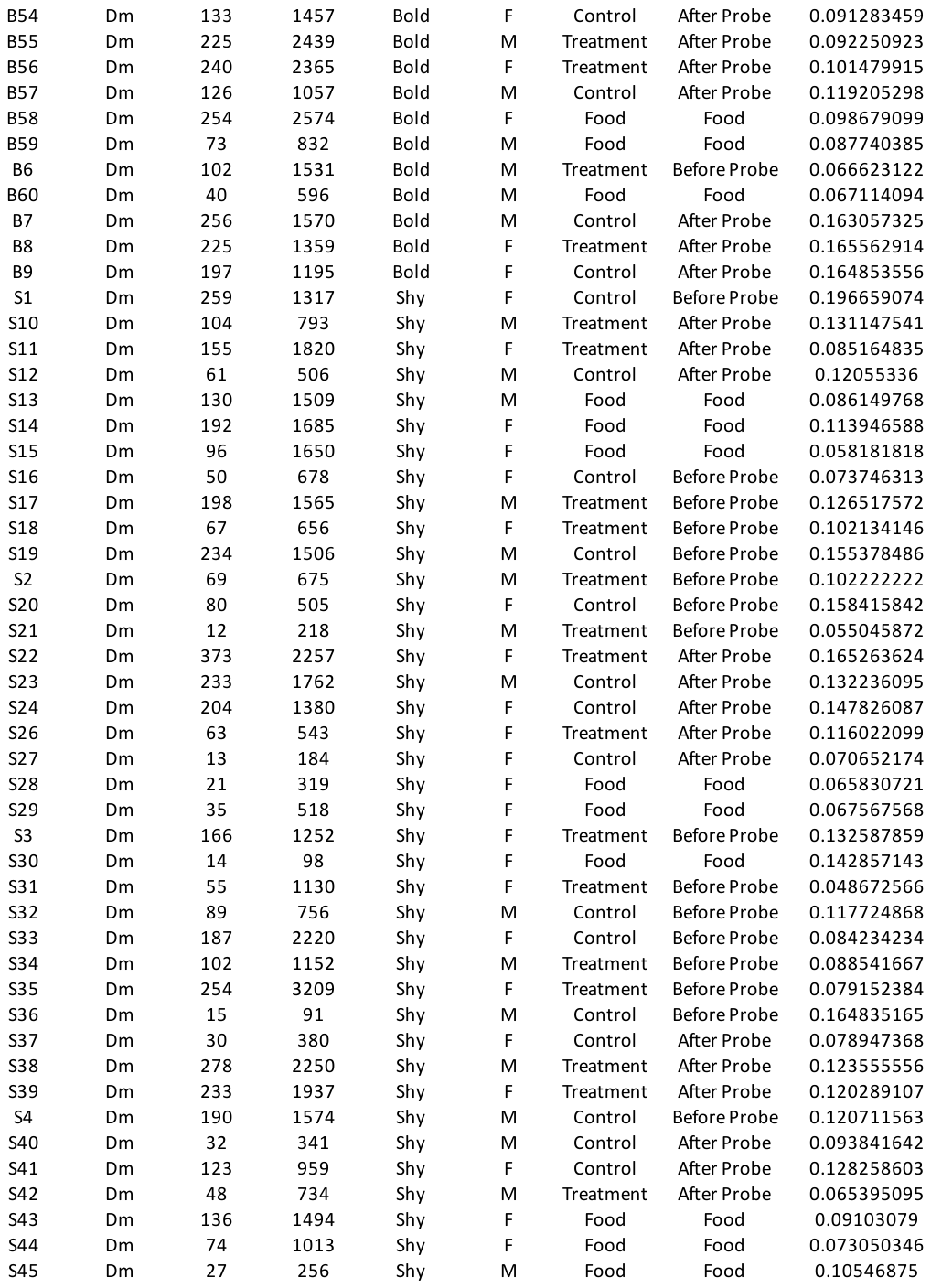

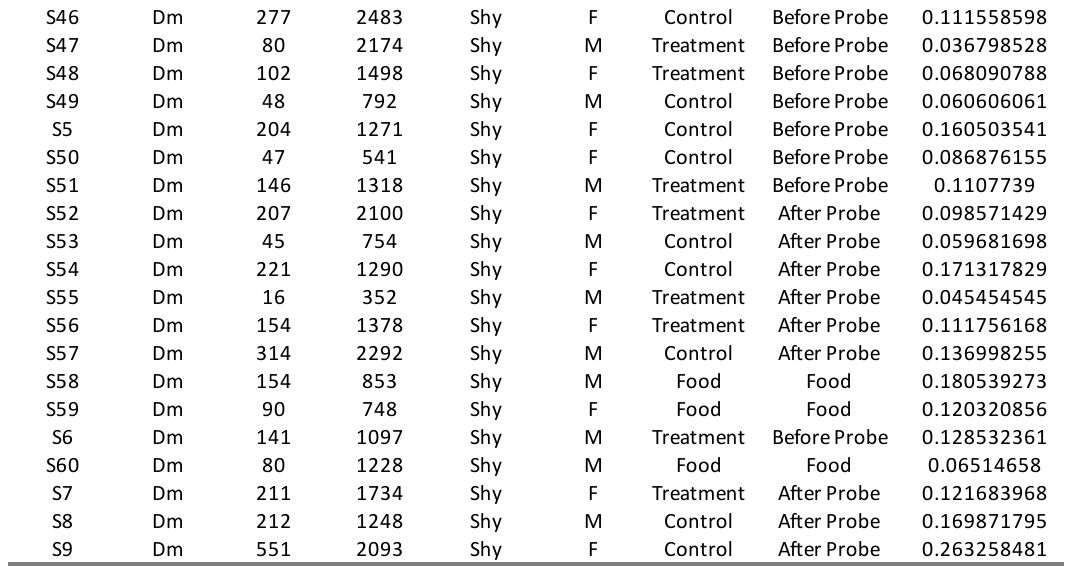

B. Dl

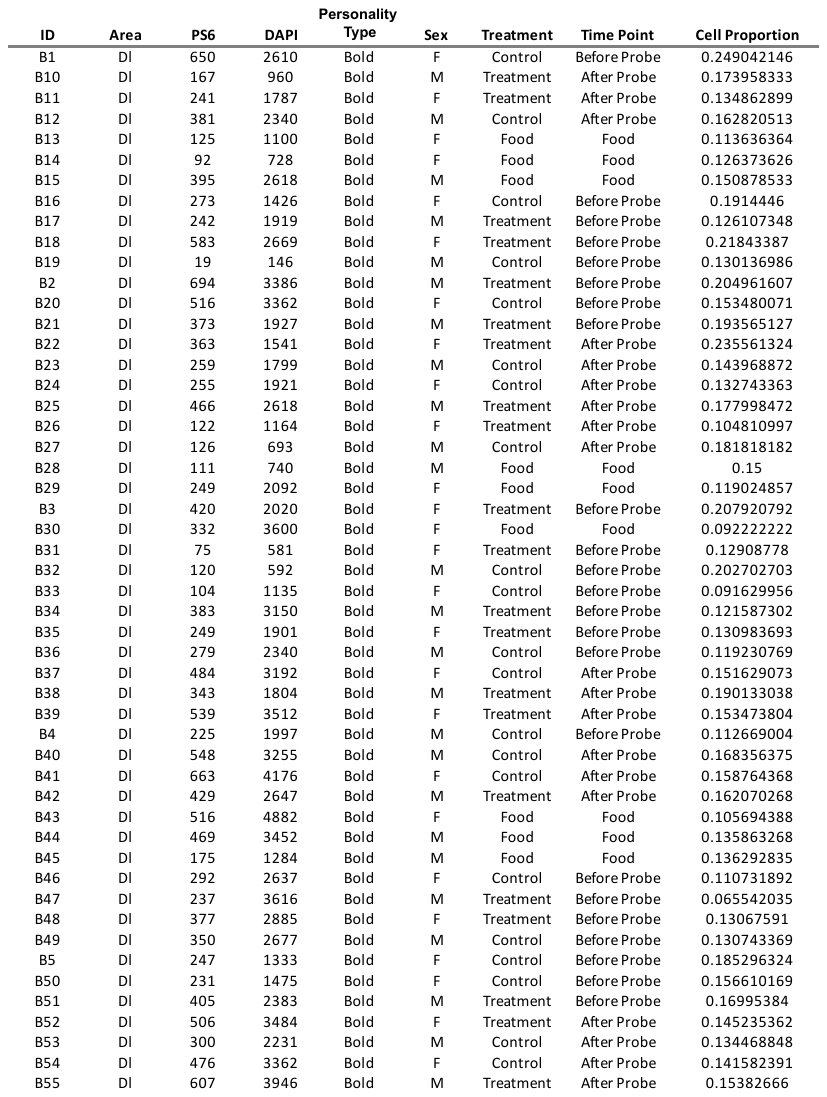

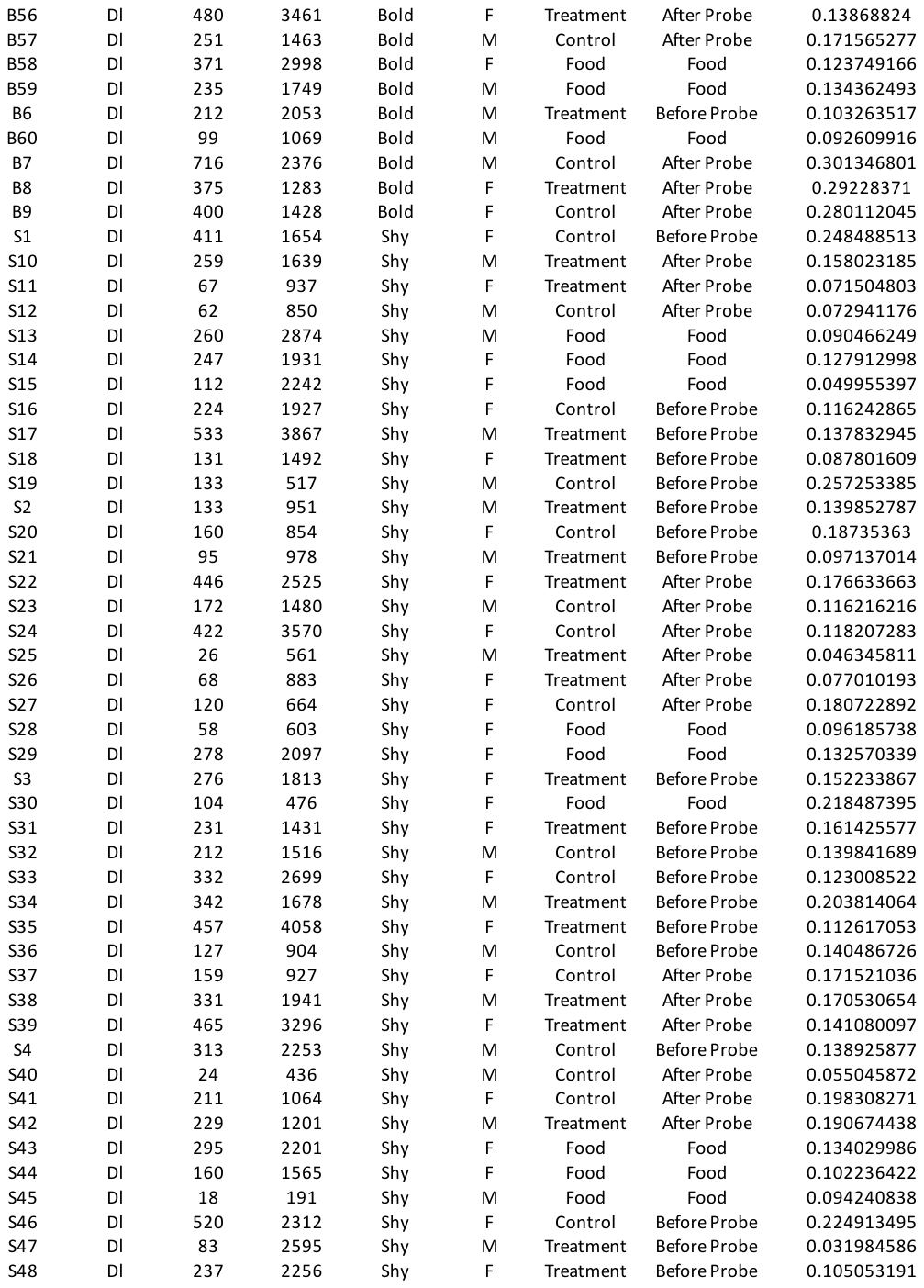

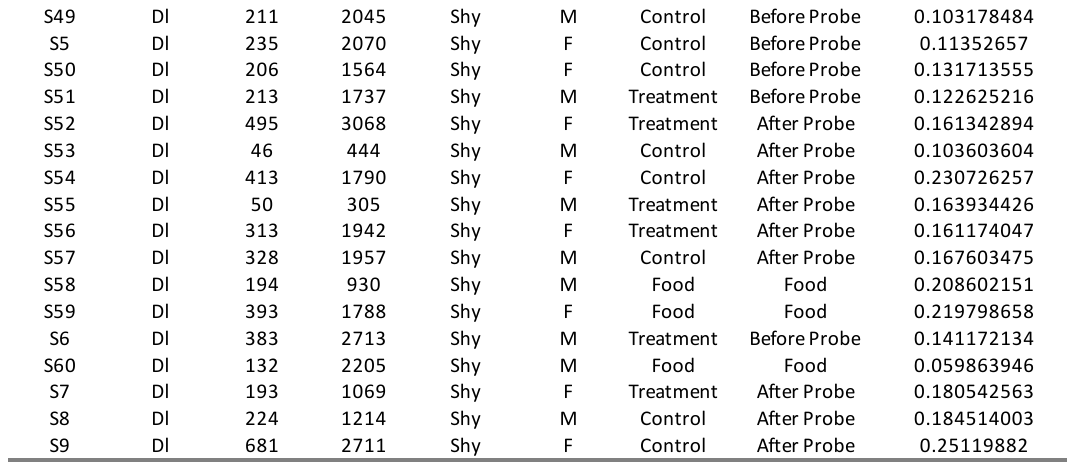

C. Dp

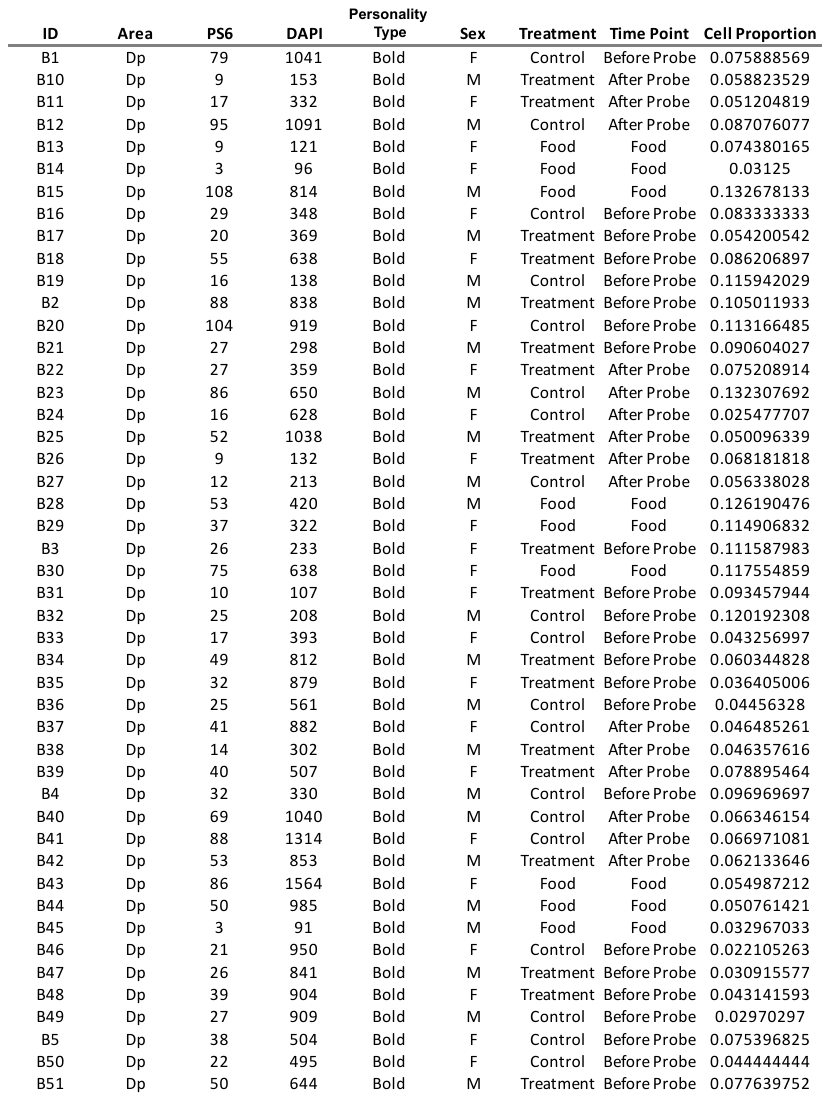

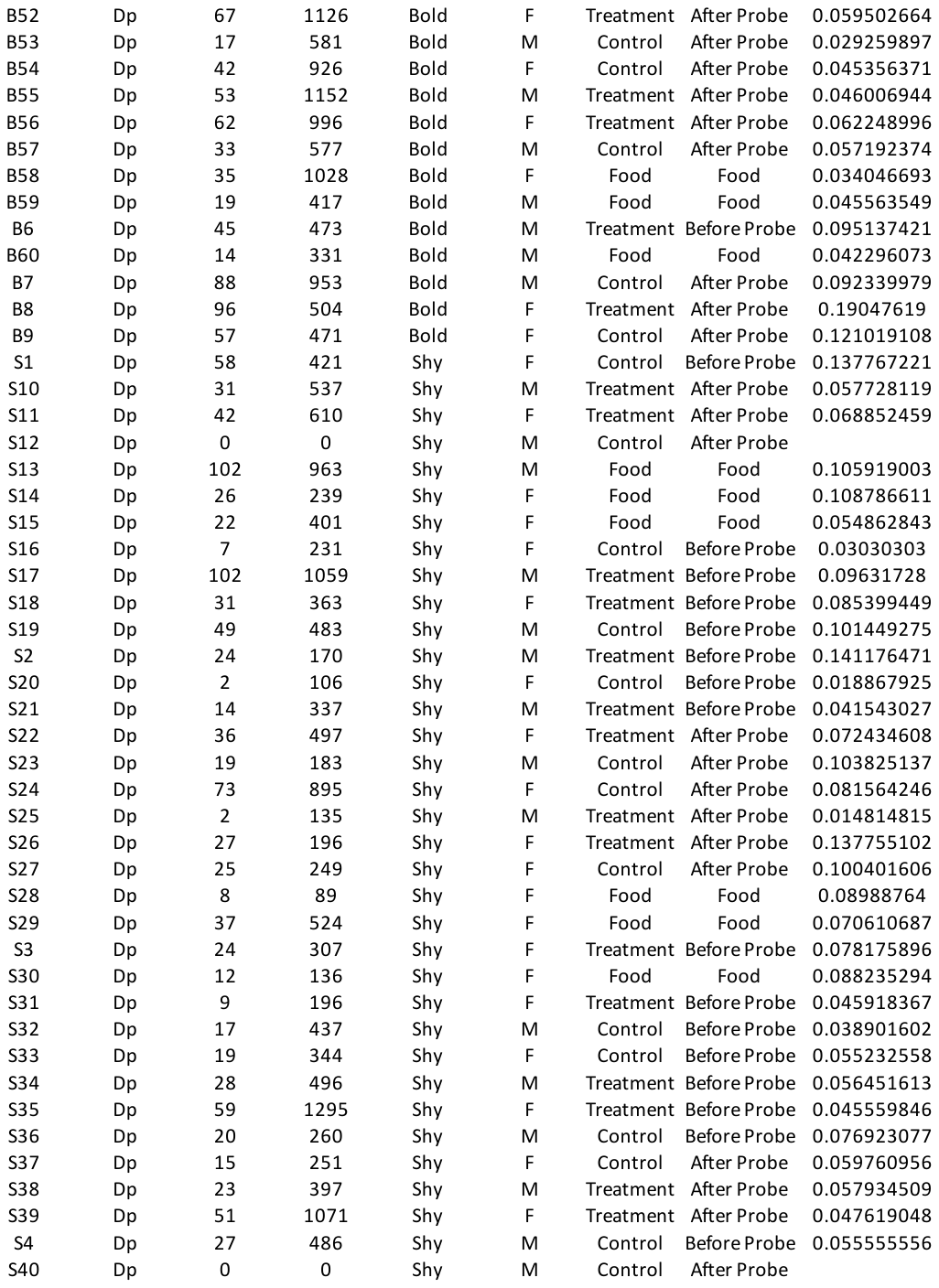

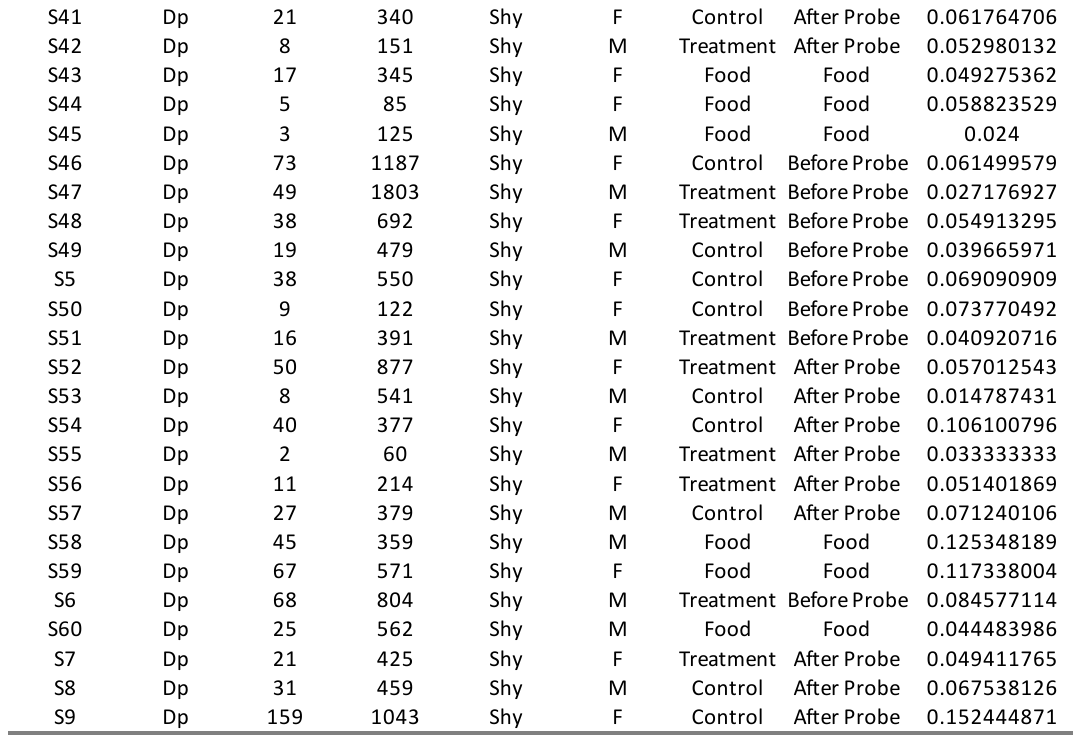

D. Hb

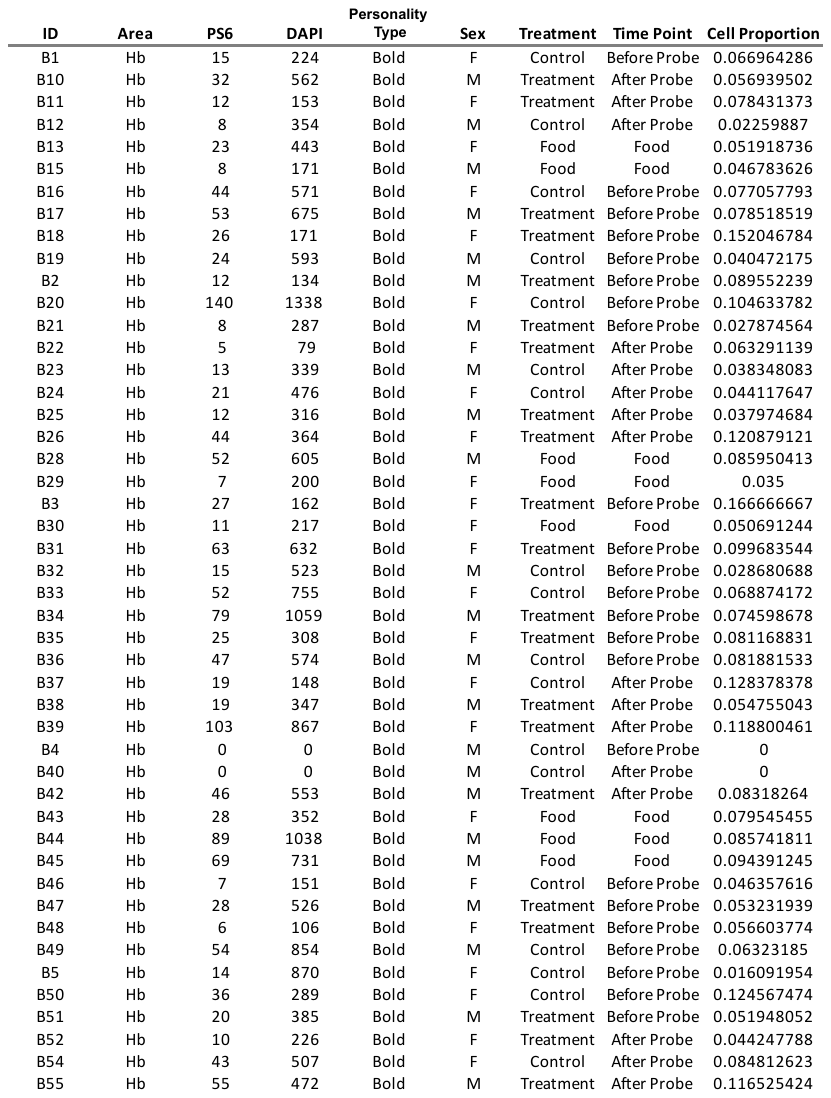

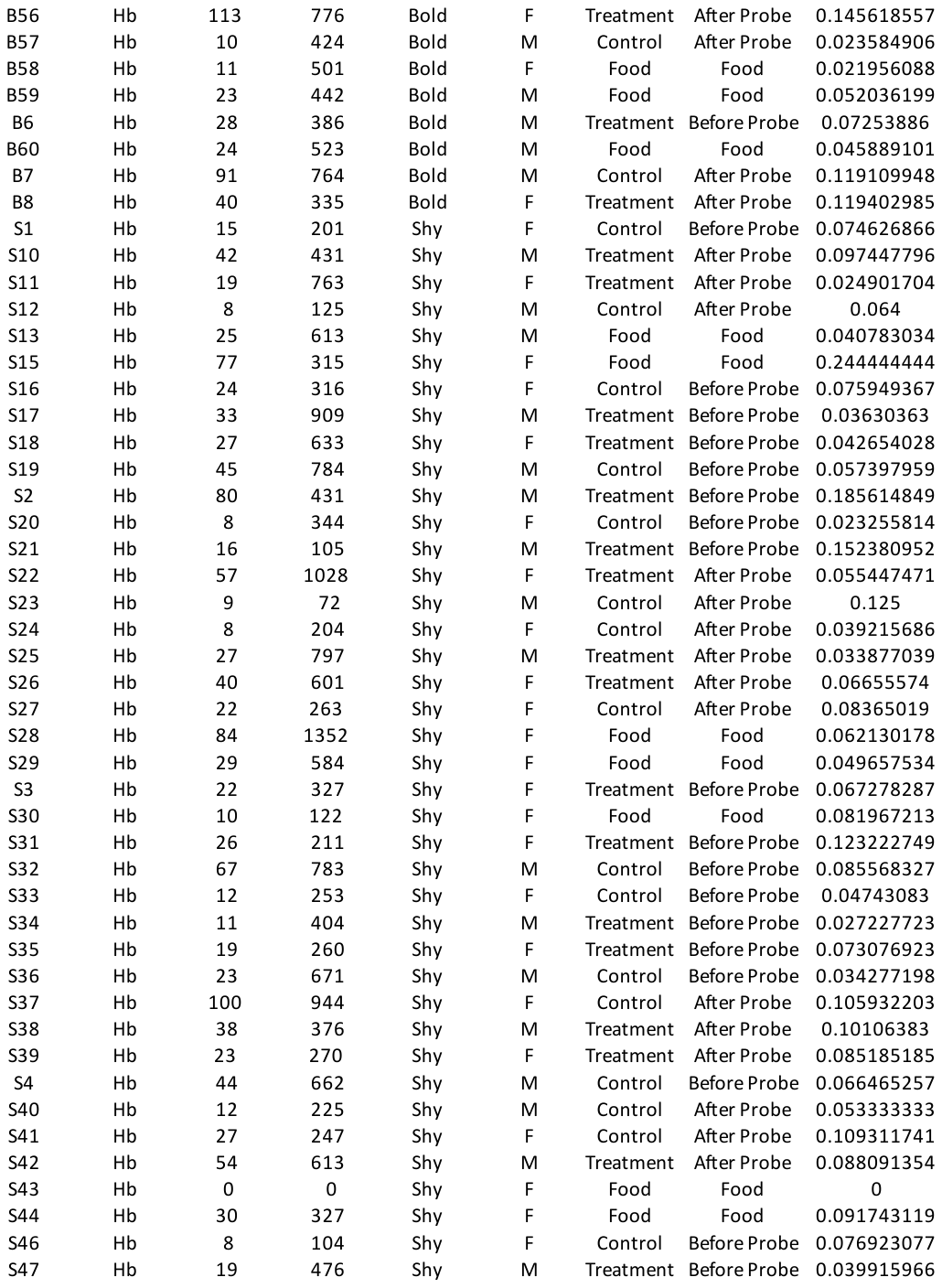

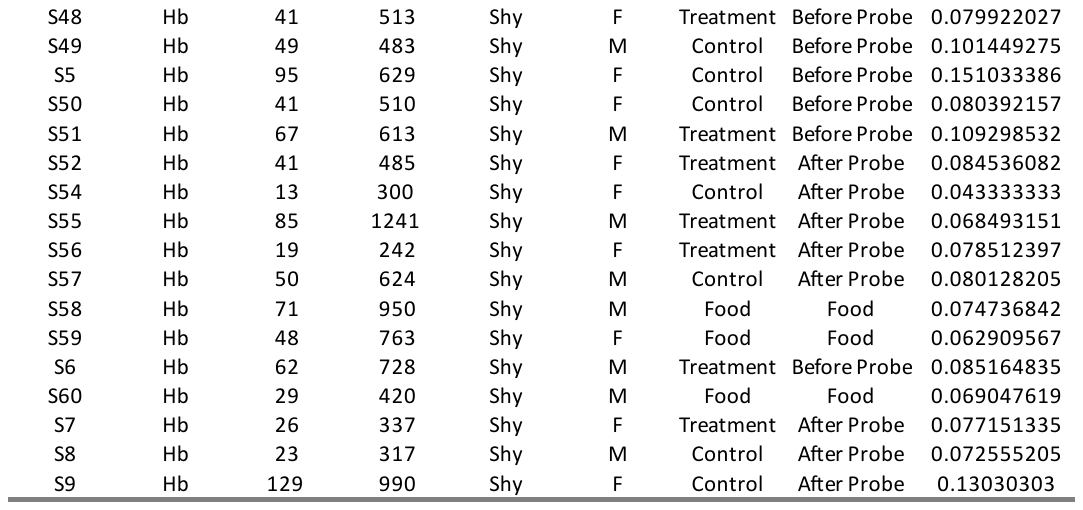

E. Hd

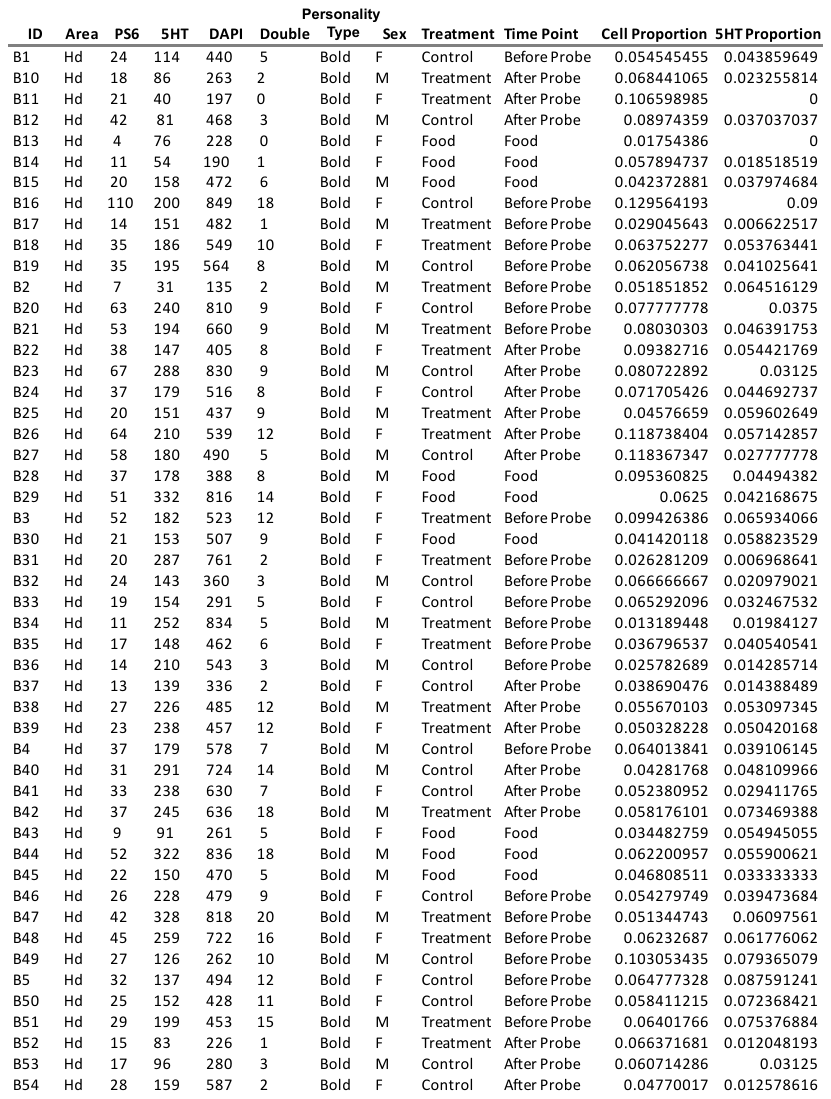

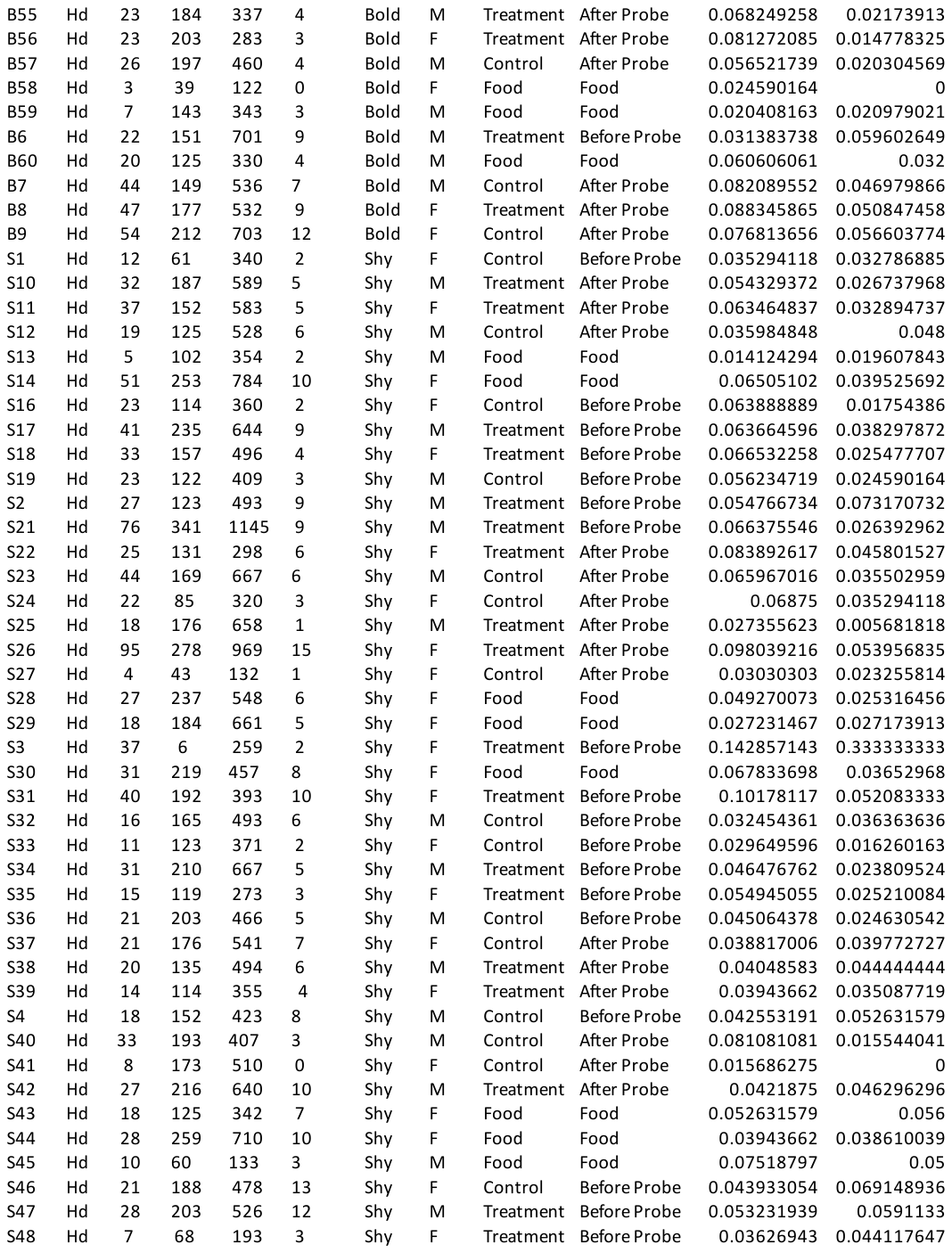

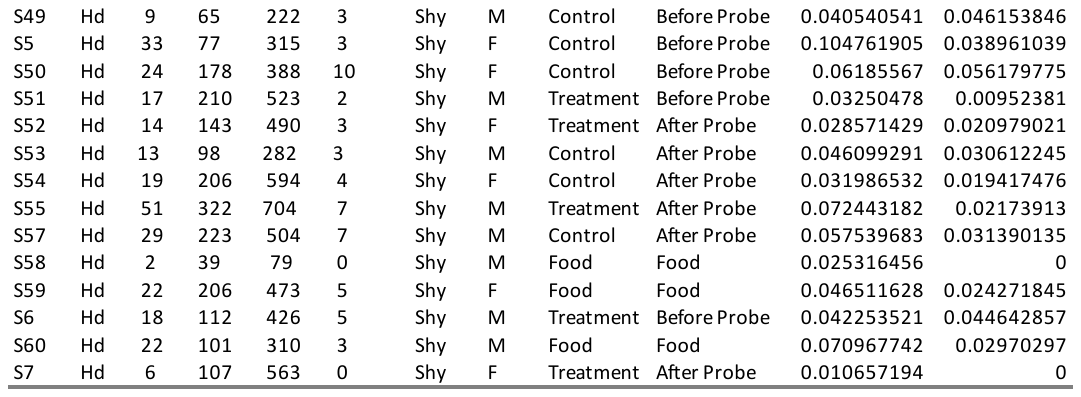

F. Hv

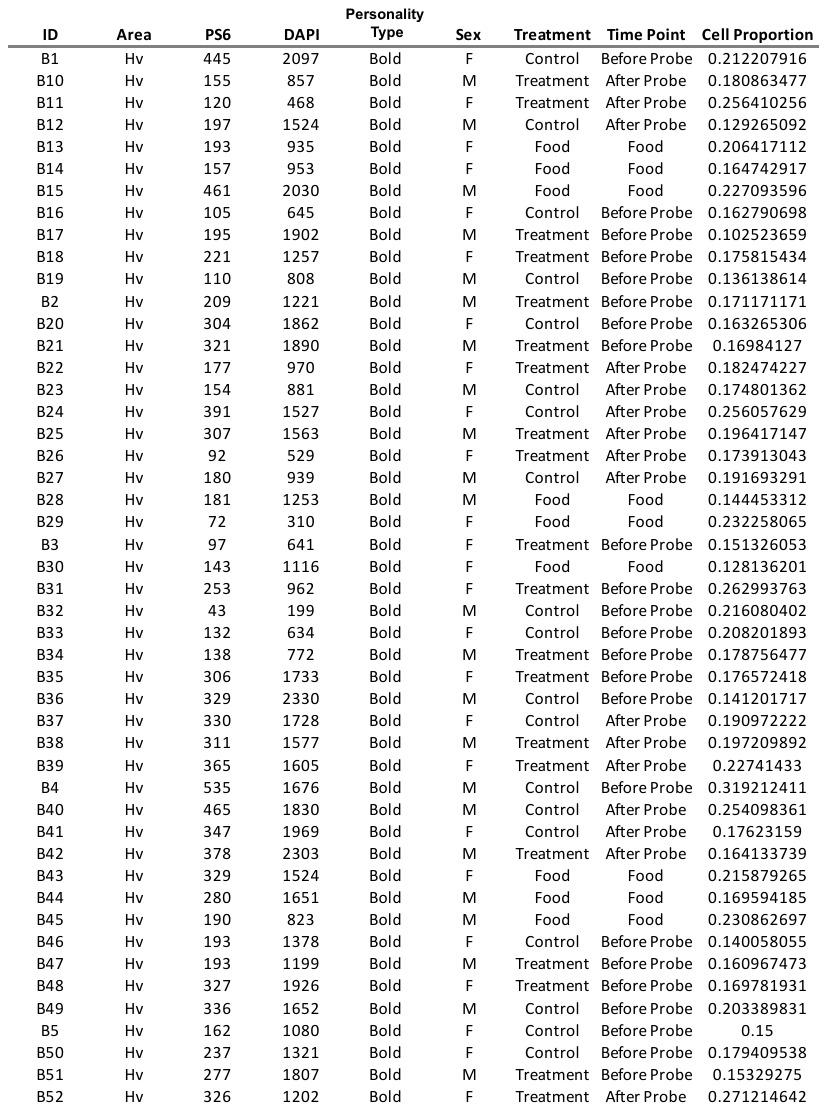

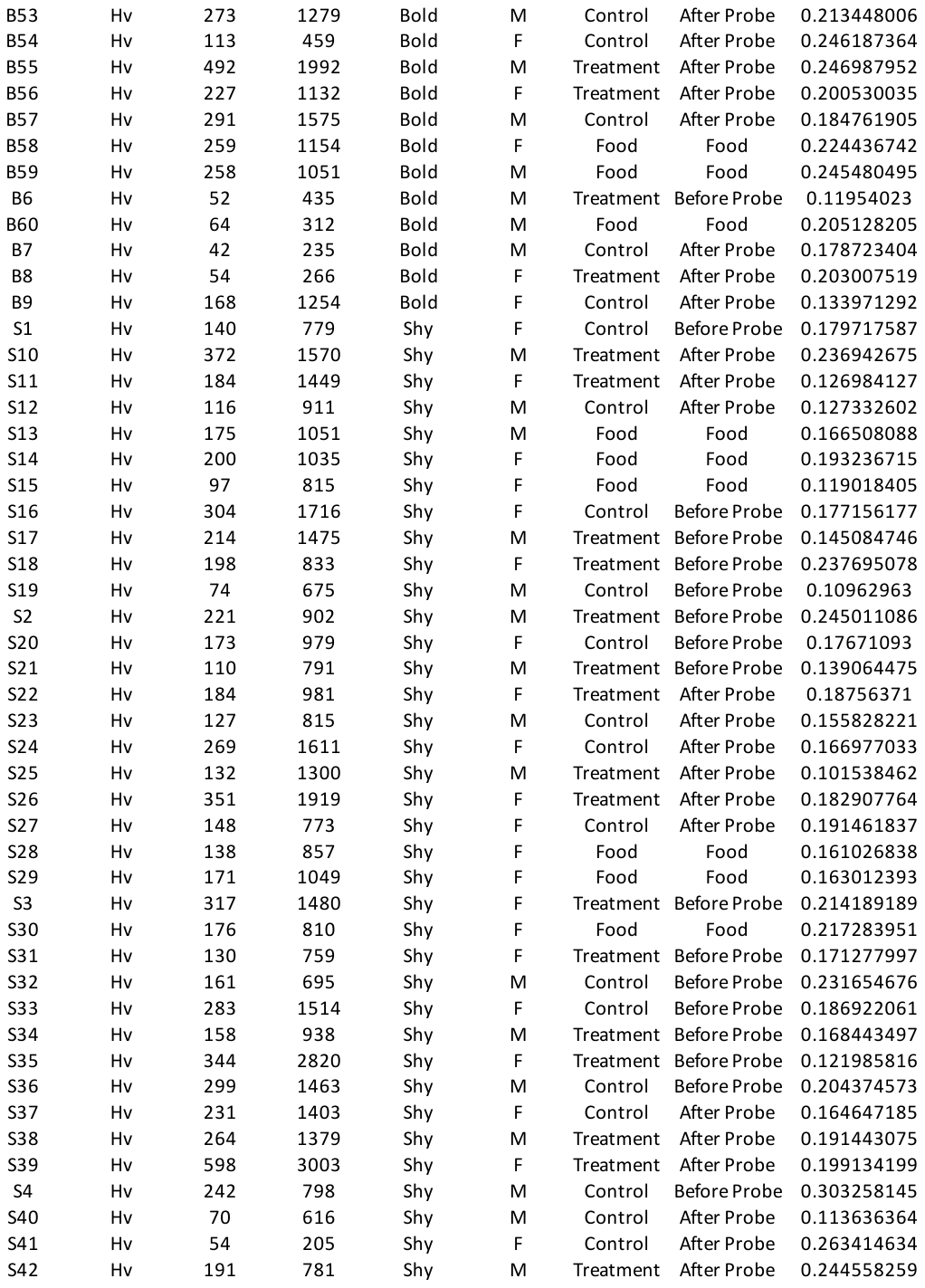

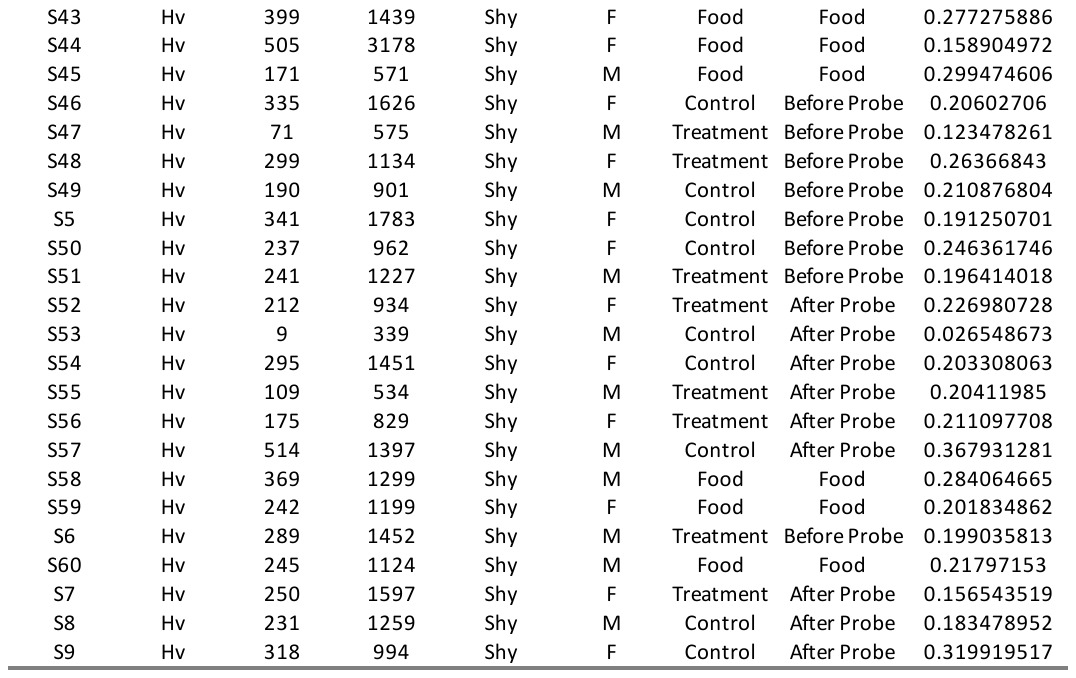

G. POA

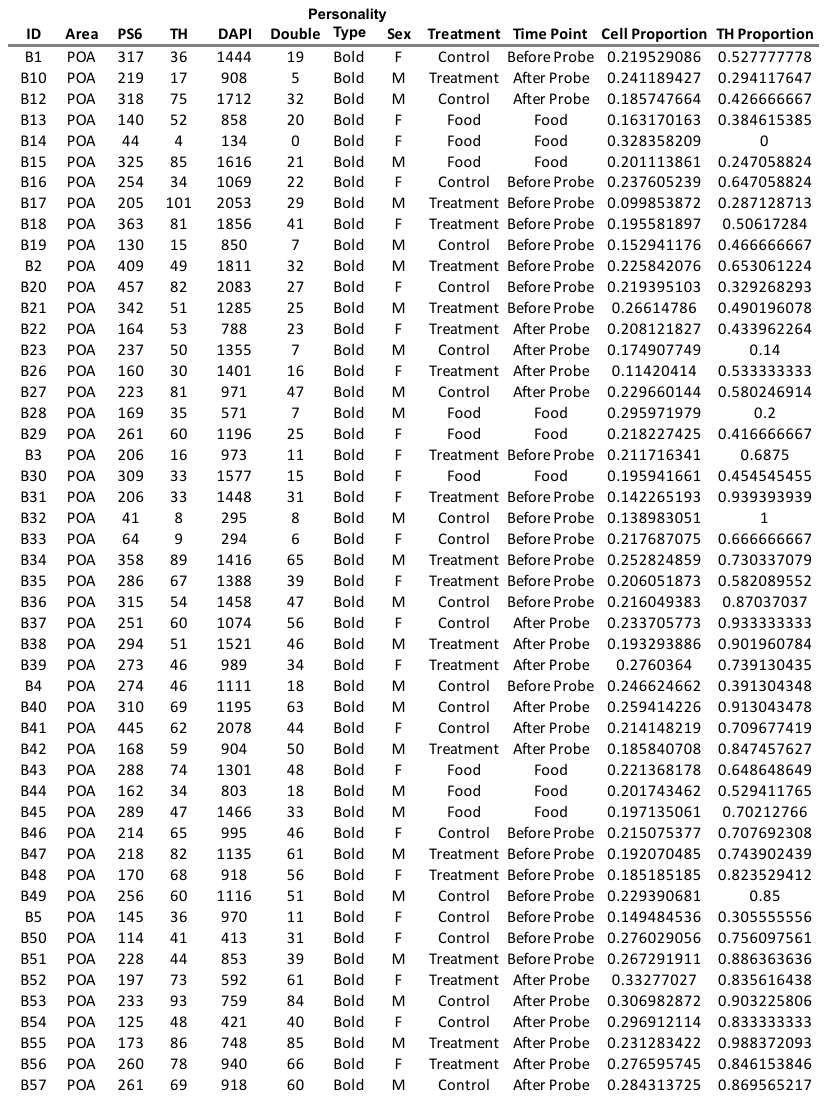

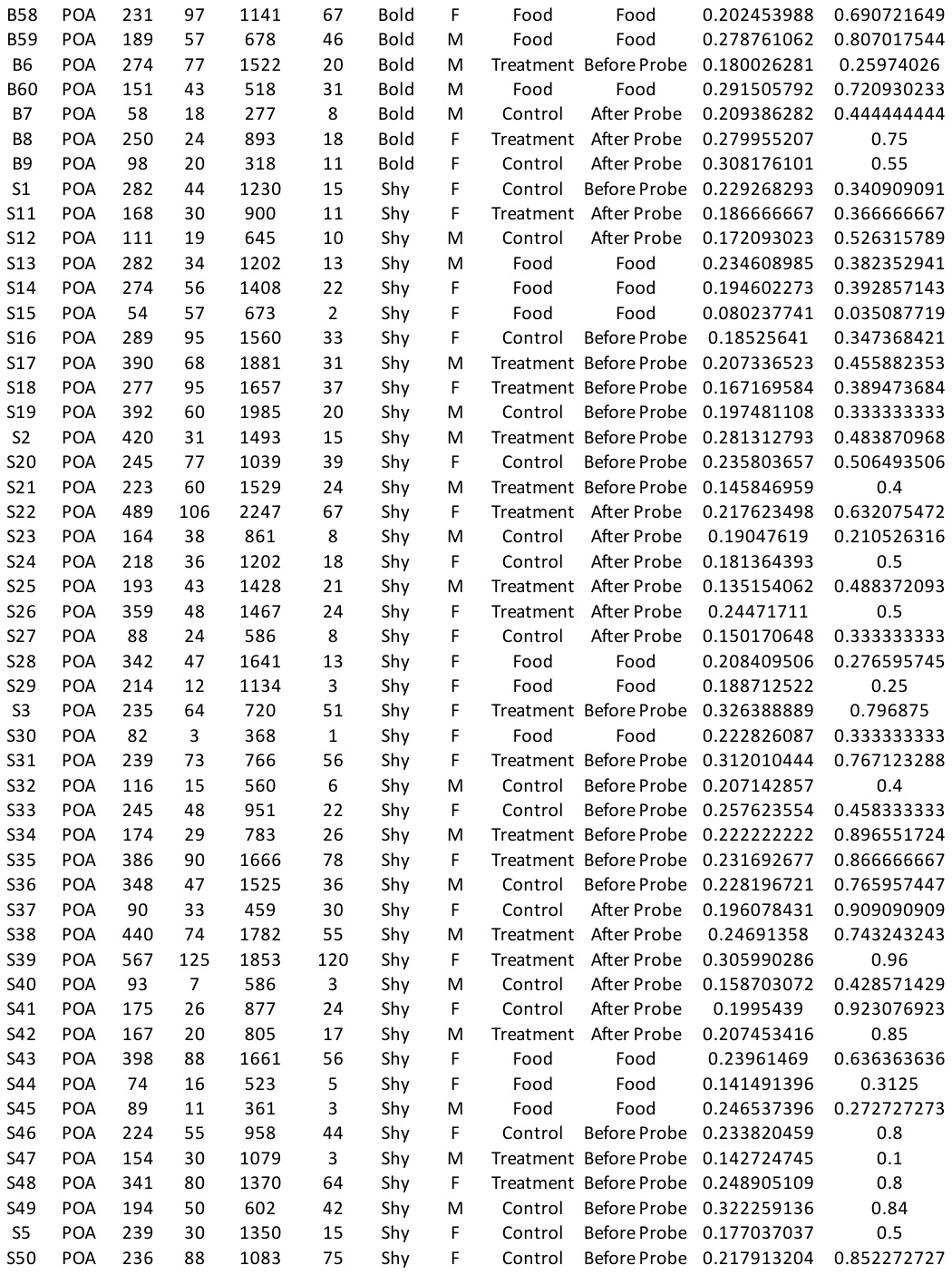

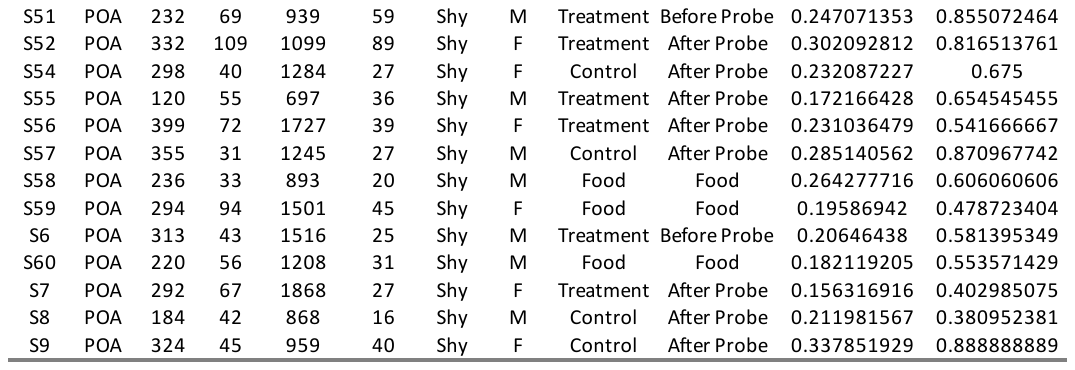

H. PT

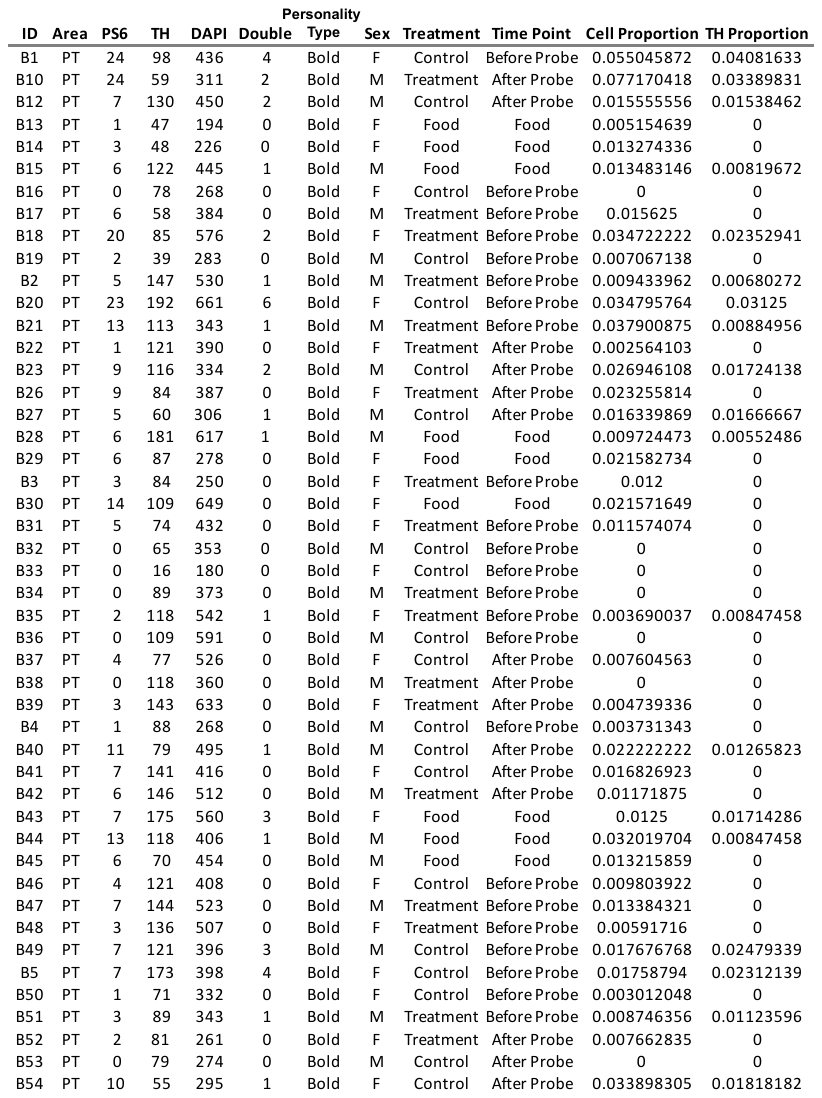

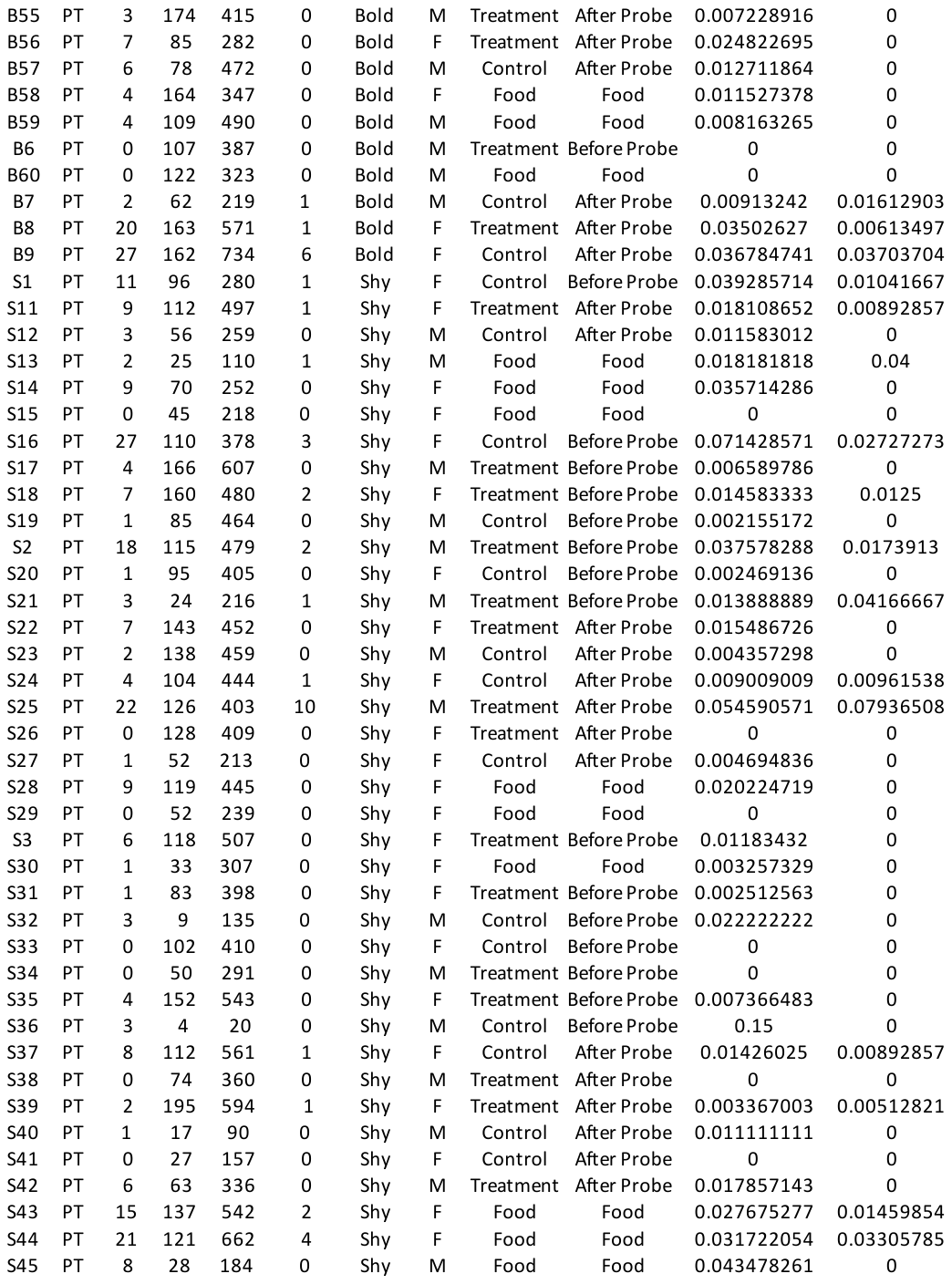

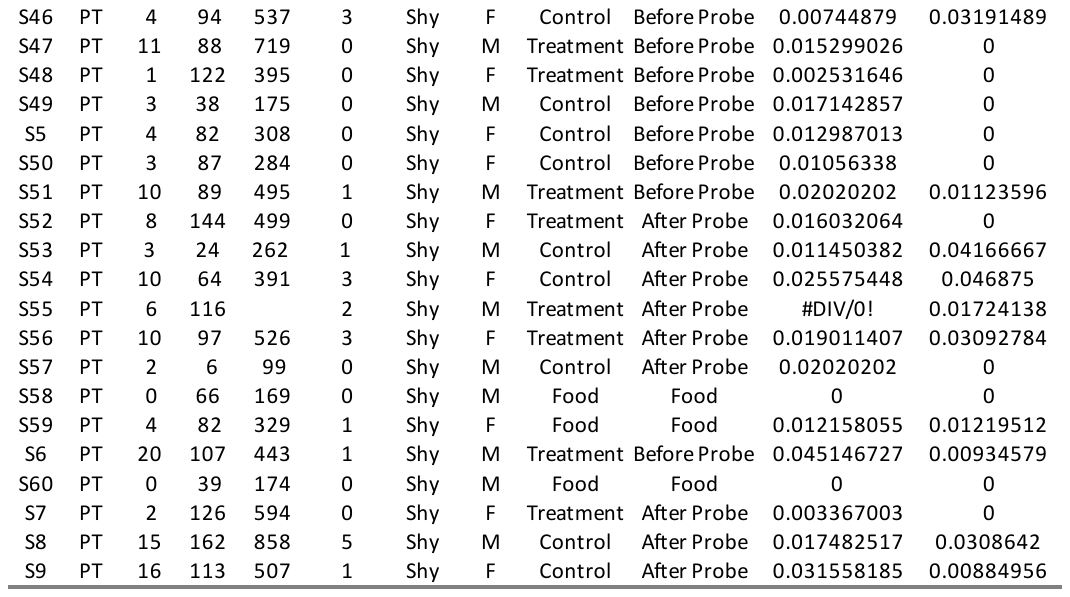

I. SR

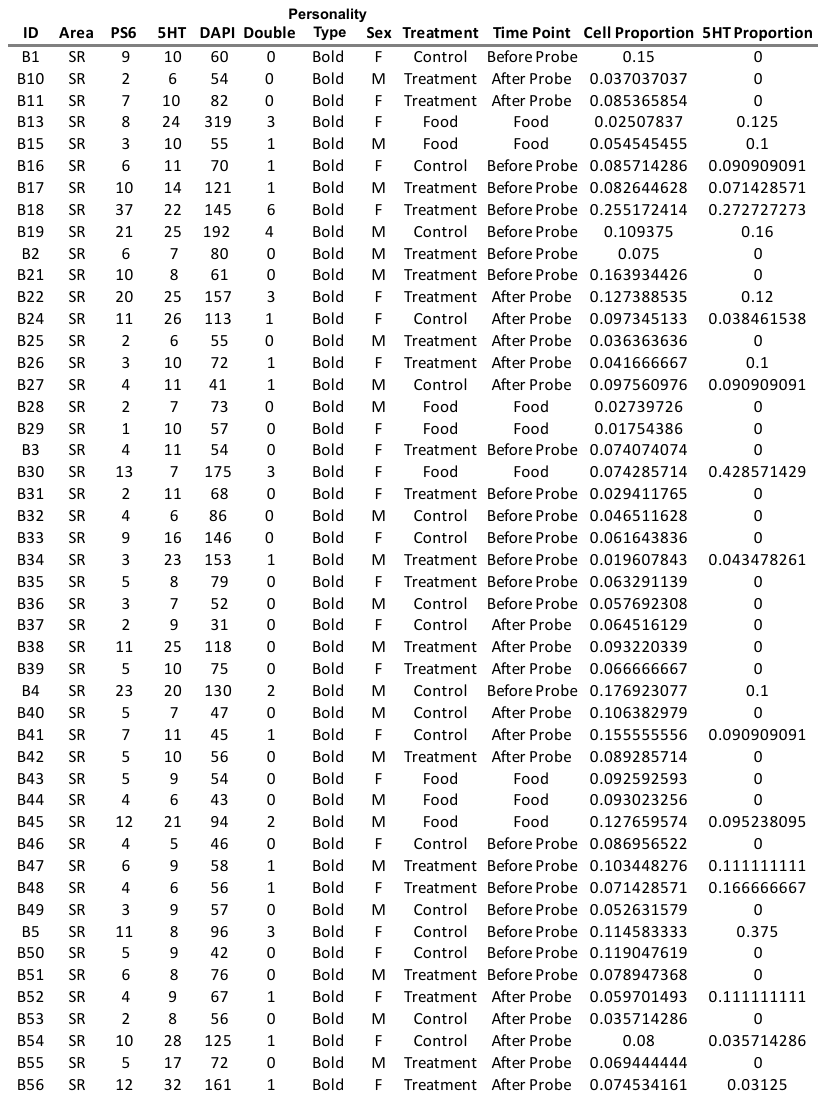

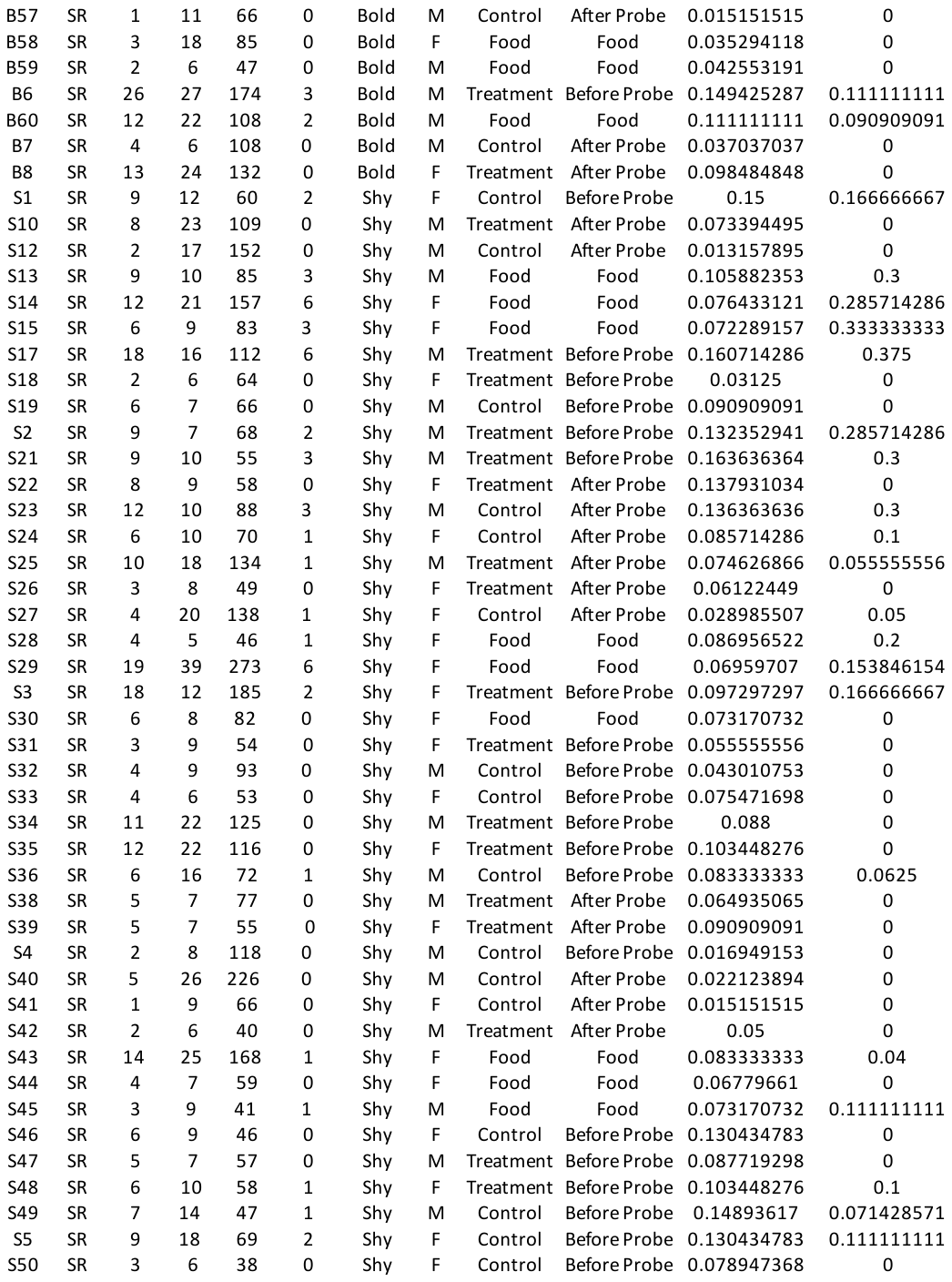

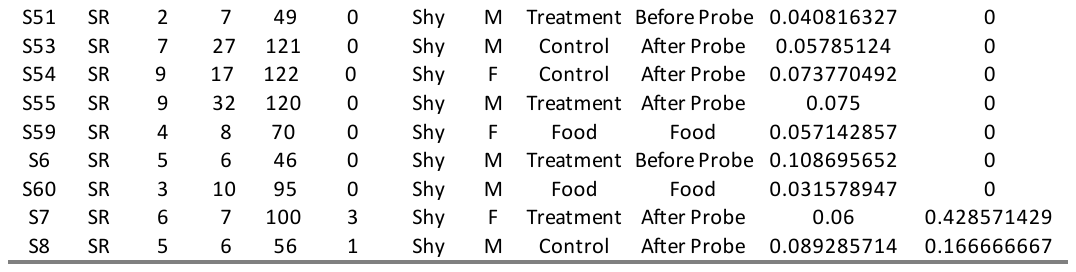

J. Tpp

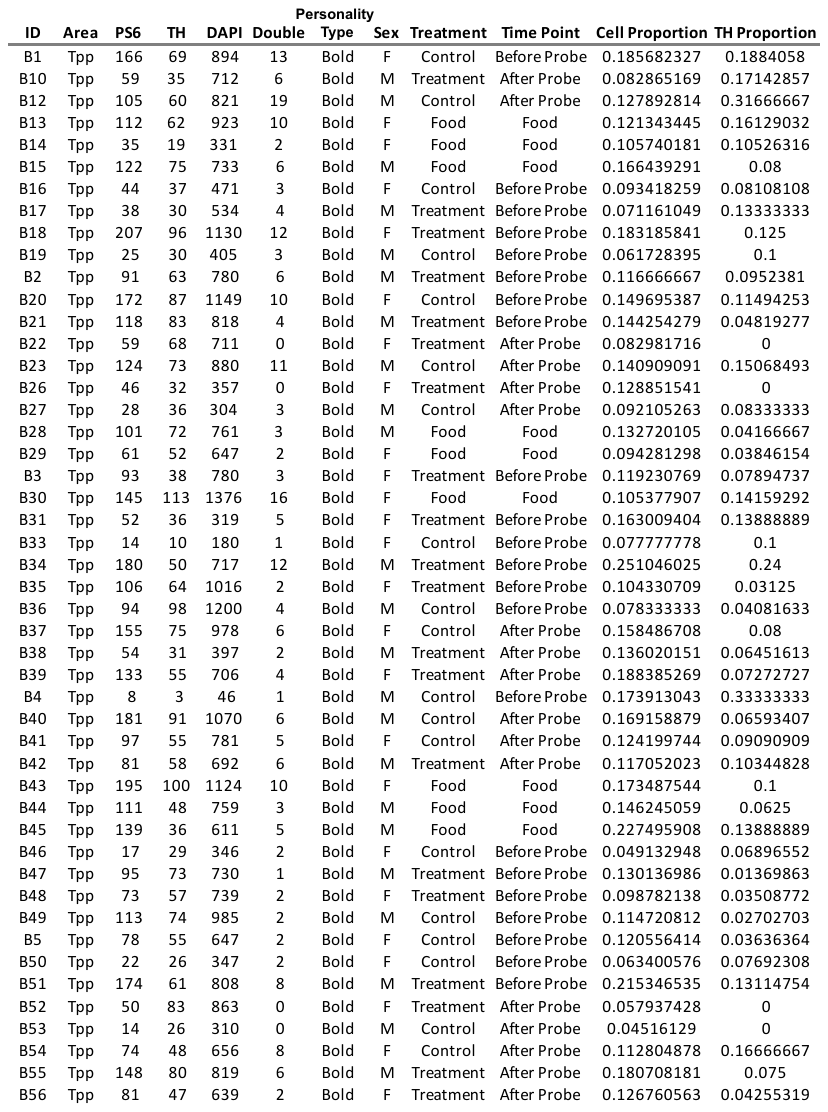

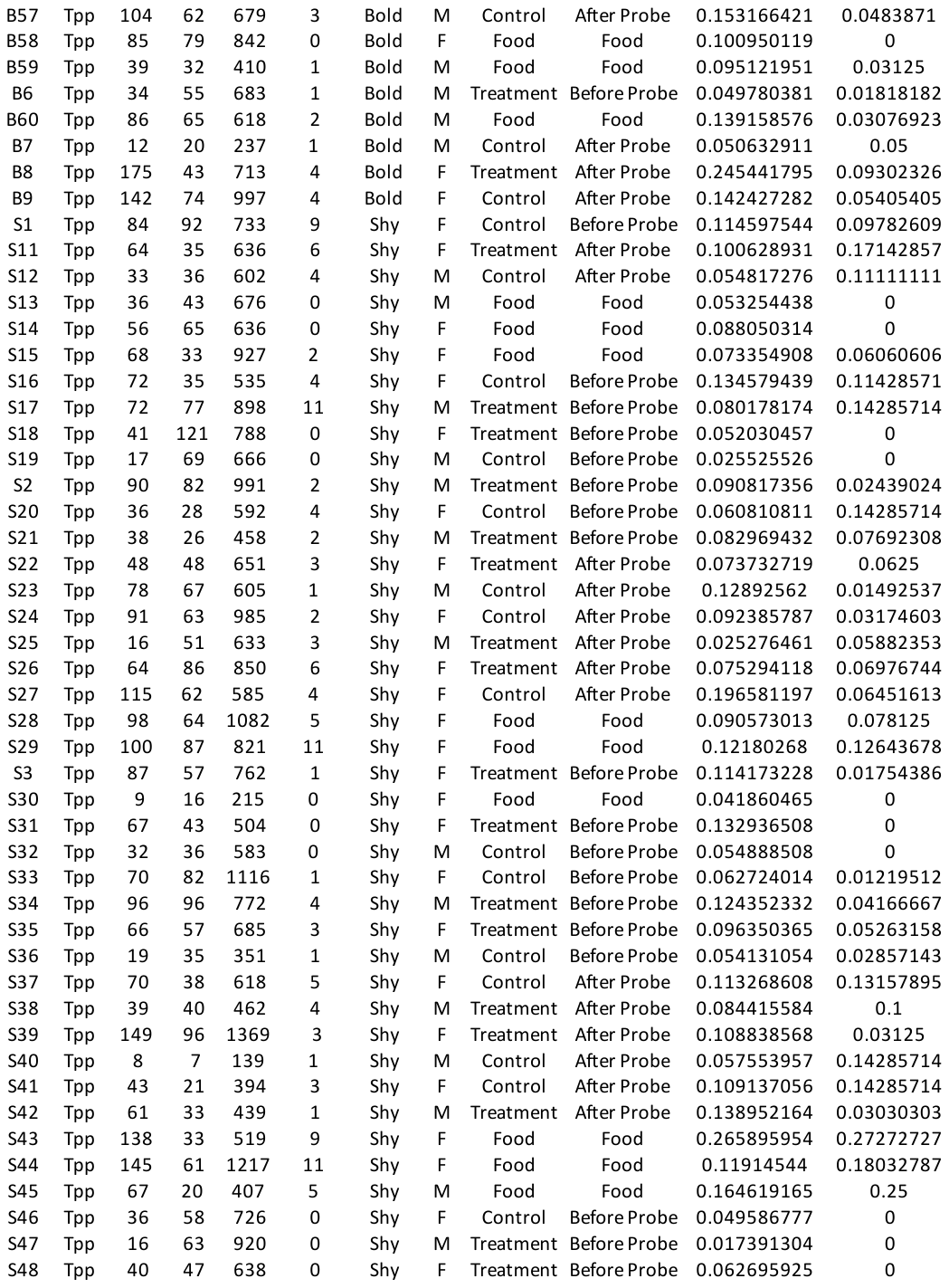

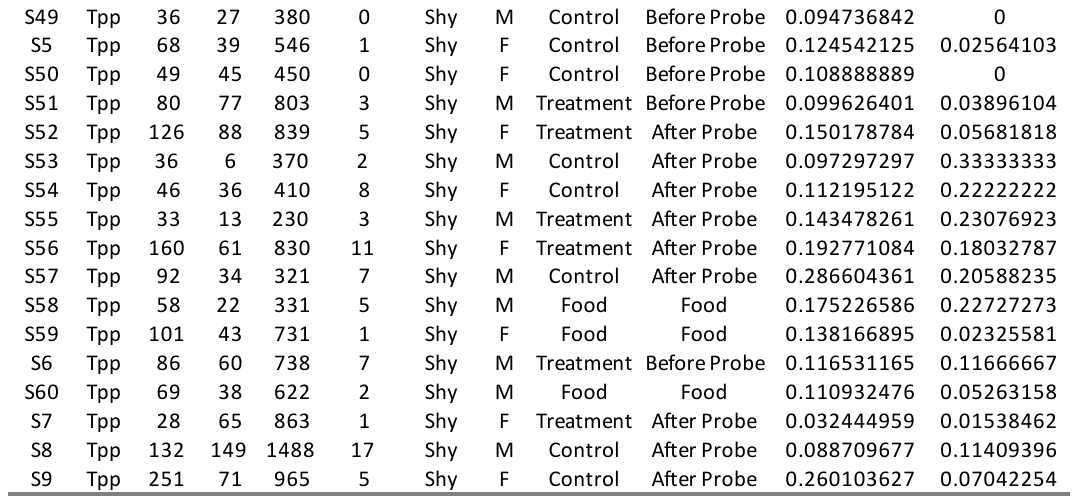

K. Vs

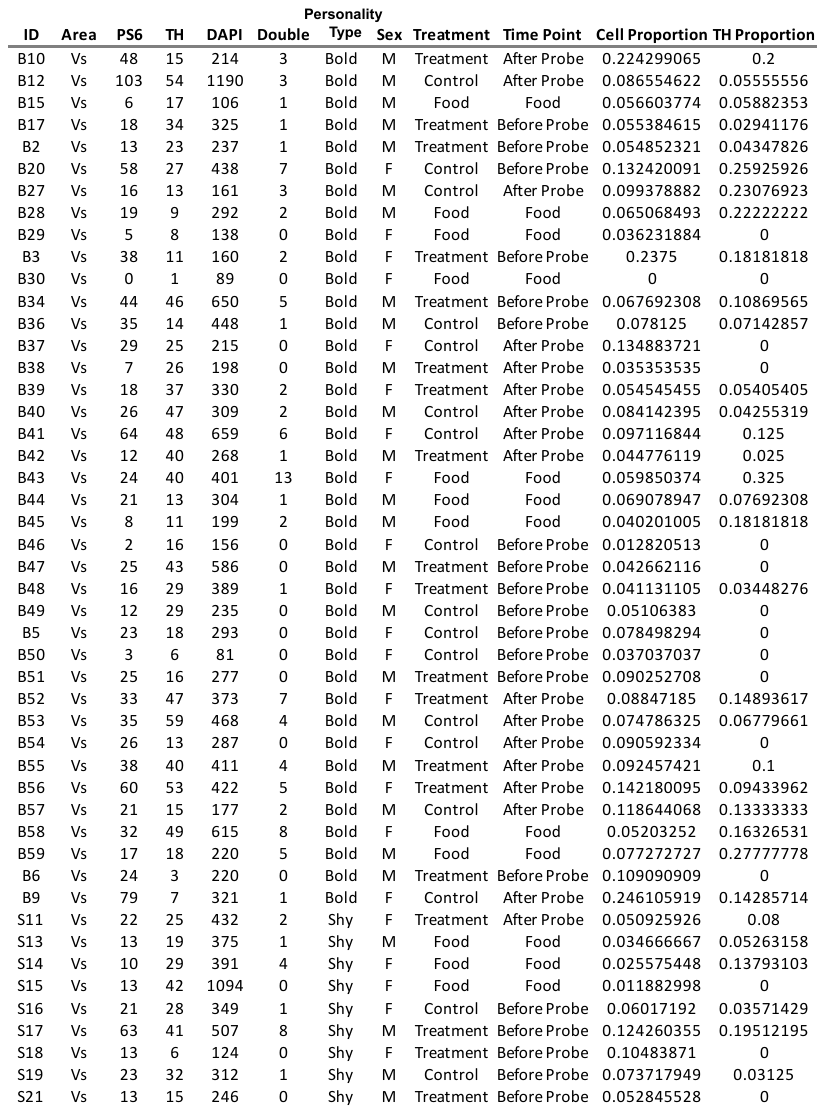

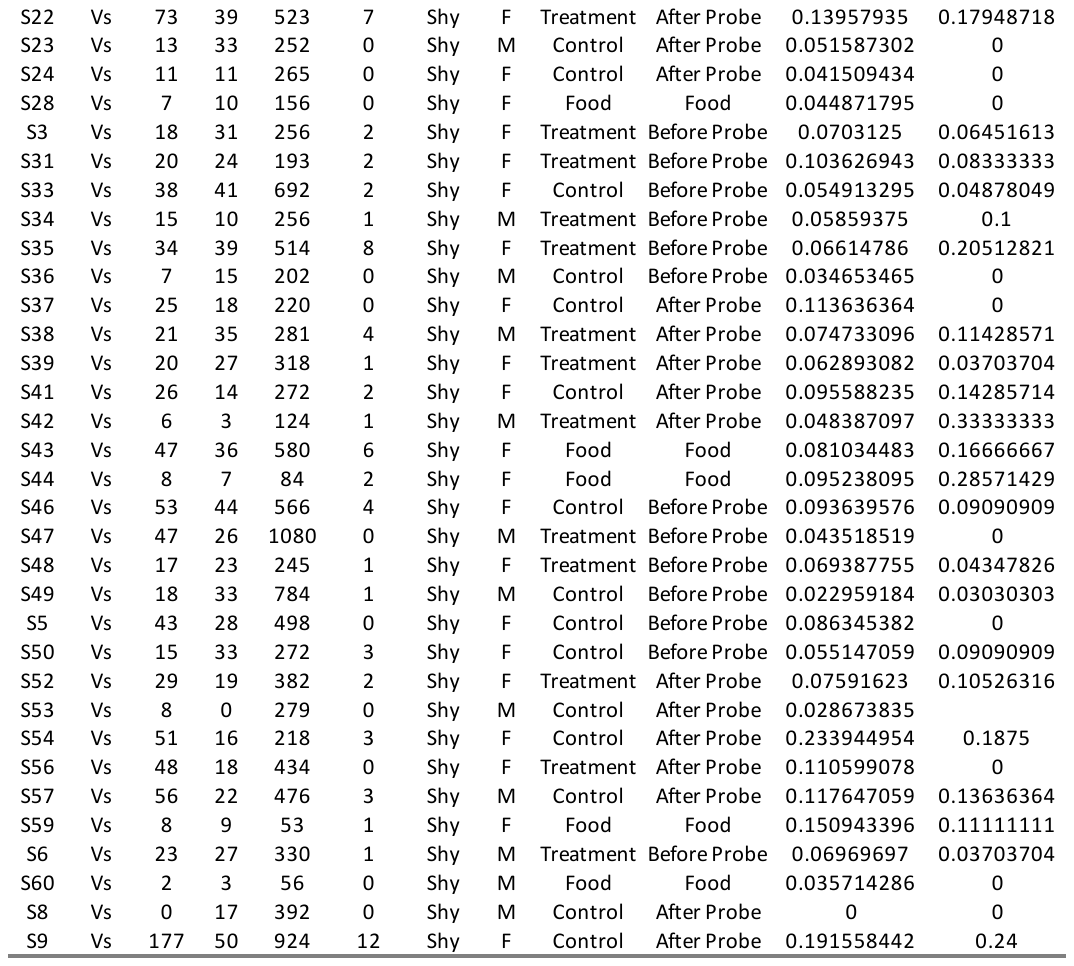

L. Vv

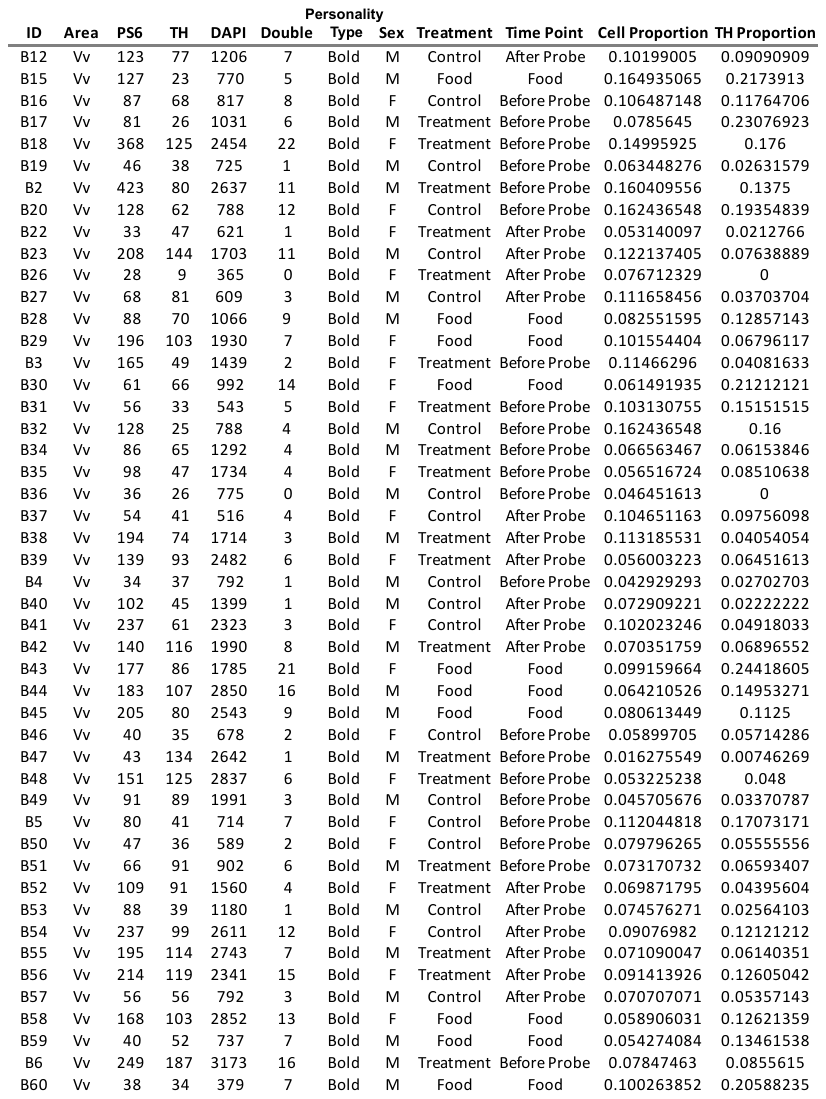

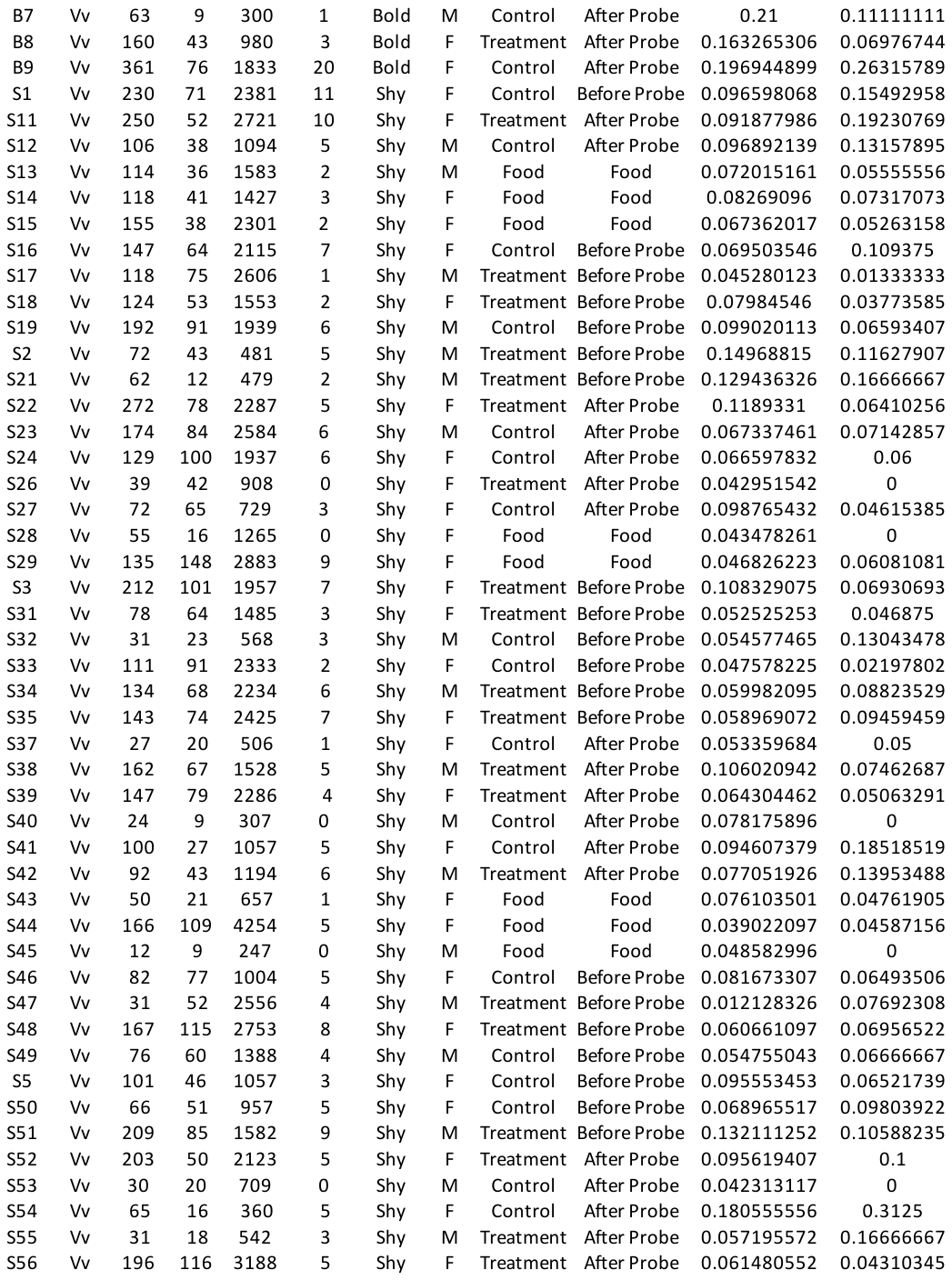

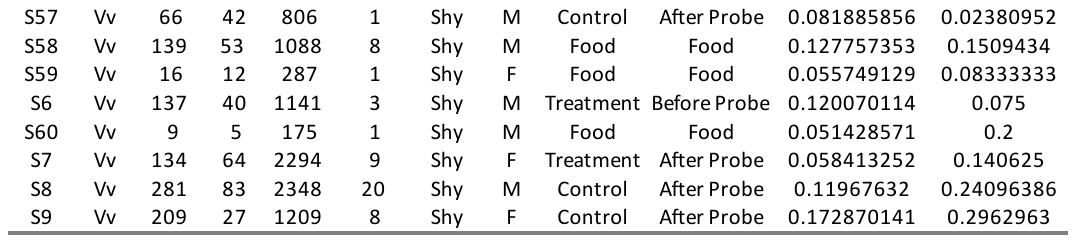
